# A multi-omics characterization reveals distinct molecular signatures in the human motor cortex and lumbar spinal cord in ALS

**DOI:** 10.64898/2026.07.24.740361

**Authors:** Natalie Barretto, Benjamin T. Fullerton, Aidan C. Daly, Obadele Casel, Olena Kuksenko, Kristy Kang, Joana Petrescu, Maya Xia, Jacqueline Eschbach, Matthew Leung, Shruti Khiste, Brhan Gebremedhin, Colin Smith, Christopher A. Jackson, Hemali Phatnani

**Affiliations:** Department of Neurology, Columbia University, New York, NY, USA; Motor Neuron Center, Columbia University Irving Medical Center, New York, NY, USA; Center for Genomics of Neurodegenerative Disease, New York Genome Center, New York, NY, USA; Centre for Clinical Brain Sciences, University of Edinburgh, Edinburgh, EH16 4TG, UK; UK Dementia Research Institute, University of Edinburgh, Edinburgh, EH16 4SB, UK

**Author notes:** Corresponding Authors (CAJ) and (HP).

## Abstract

Amyotrophic lateral sclerosis (ALS) is a debilitating neurodegenerative disease characterized by the loss of upper motor neurons in the motor cortex (MTC) and lower motor neurons in the spinal cord, leading to muscle atrophy and ultimately respiratory failure. While motor neurons (MNs) are the selectively vulnerable cell type, their interactions with glia contribute to the progression of ALS pathology. However, it remains unclear whether the site of ALS symptom onset influences the molecular alterations underlying MN and glial dysfunction and whether these alterations are shared between the MTC and lumbar spinal cord (LSC). To address these questions, we constructed spatially-resolved gene expression maps of the MTC and LSC by combining spatial and single-nucleus transcriptomic profiles from a cohort of non-neurological controls and ALS donors clinically stratified by site of symptom onset. In the ventral horn of the LSC, we see a decrease in genes associated with MNs and synaptic signaling in ALS donors. We also identify region-specific alterations in endothelial- and glial-related functions. Notably, the severity of these MN deficits and endothelial-related functions is influenced by the site of symptom onset, whereas alterations in glial function largely are not. In contrast to the LSC, we observe layer-specific increases in synaptic signaling in the MTC of ALS donors. Comparing the molecular and cellular changes within the LSC and MTC in ALS indicates that they are predominantly non-overlapping, and have different molecular signatures.

## INTRODUCTION

Amyotrophic lateral sclerosis (ALS) is a debilitating neurodegenerative disease characterized by the loss of upper motor neurons in the motor cortex (MTC) and lower motor neurons in the spinal cord, leading to muscle atrophy and ultimately respiratory failure. The clinical presentation of ALS can be stratified by site of symptom onset: bulbar-onset patients experience muscle weakness beginning in the face, neck, or tongue, resulting in difficulties with speaking and swallowing, whereas limb-onset patients exhibit muscle weakness beginning in the hand, arm, or leg. Disease progression often affects anatomically adjacent regions, suggesting that the underlying pathology begins focally but propagates through the nervous system in an organized manner^1–4^. In support of this, motor neuron (MN) loss is greatest in regions proximal to site of symptom onset^5^. However, it is unclear whether the site of symptom onset influences all aspects of ALS pathology or primarily MN function.

While MNs are the selectively vulnerable cell type, their interactions with glia contribute to the progression of ALS pathology. Microgliosis^6–9^ and astrogliosis^10–13^ are commonly observed in post-mortem MTC and spinal cord tissue from ALS patients. Specifically, reactive microglia and astrocytes are found near the corticospinal tracts^12,14,8^, which are composed of upper MN axons, and are abundant in the ventral horn of the spinal cord^8,12–14^, where lower MNs reside. In cell culture experiments, introducing microglia^15,16^ or astrocytes^13,17,18^ isolated from ALS patients or mouse models induces MN toxicity, indicating that glial dysfunction promotes MN dysfunction. Importantly, it has not been fully characterized whether the molecular alterations underlying MN and glial dysfunction in ALS are consistent between the MTC and lumbar spinal cord (LSC).

To address these questions, we constructed spatially-resolved gene expression maps of the MTC and LSC by combining spatial and single-nucleus transcriptomic profiles from a cohort of non-neurological controls and ALS donors clinically stratified by site of symptom onset. In ALS donors there is a decrease in MN gene expression and synaptic signaling in the ventral horn of the LSC. There are also region-specific alterations in endothelial- and glial-related functions. The site of symptom onset (bulbar or limb) strongly influences the severity of MN defects and changes to endothelial tissue, but not glial composition and function. In the MTC, we observe layer-specific increases in synaptic signaling-associated genes in ALS donors. Molecular and cellular changes linked to ALS differ within the LSC and MTC, and are non-overlapping, distinct molecular signatures.

## RESULTS

### Experimental design and cohort overview

We sampled LSC tissue from 42 donors (14 non-neurological controls, 28 ALS) and sampled MTC tissue from 50 donors (12 controls, 38 ALS) (Figure 1A, Supplemental Table 1). ALS donors were clinically stratified by site of symptom onset. We measured individual cell transcriptomes and chromatin accessibility with the 10x Genomics multiomics system on nuclei isolated from LSC (20 donors: 9 controls; 11 ALS) and MTC (24 donors: 8 controls; 16 ALS) (Supplemental Figure 1). Single-nucleus RNA sequencing (snRNA-seq) transcriptomes were annotated for cell types based on gene expression profiles and established markers from the literature. After excluding low-quality nuclei and doublets, we assigned ten broad CNS cell types to 121,881 high-quality LSC nuclei: excitatory neurons (EN), inhibitory neurons (IN), motor neurons (MN), oligodendrocytes (OLIGO), oligodendrocyte progenitor cells (OPC), astrocytes (AST), microglia (MIG), immune (IMM), endothelial (ENDO), and meninges (MENING) (Figure 1Bi). The distribution of LSC cell types across donors revealed a significant increase in both the number and proportion of astrocytes in ALS donors compared to controls (Figure 1Bii-iii), a likely consequence of astrogliosis, the proliferation of reactive astrocytes in response to ALS-related pathology.

**Figure 1.**
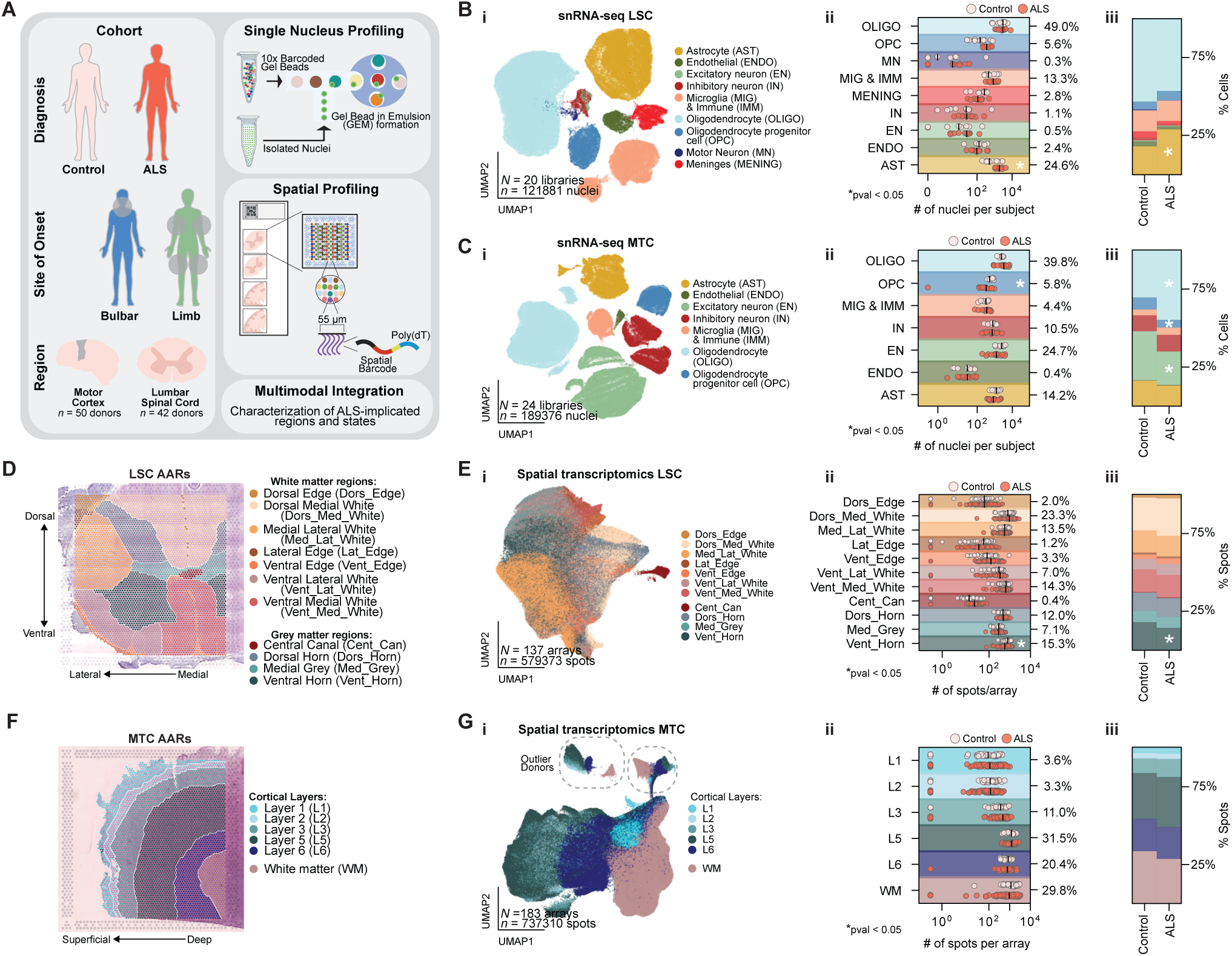
Experimental design and cohort overview. **A)** Overview of experimental design for generating spatially-resolved gene expression maps of the MTC and LSC. **B)** i) Uniform manifold approximation and projection (UMAP) plot of LSC snRNA-seq data. Each data point represents one nucleus, color-coded by cell type. ii) Number of nuclei annotated per cell type across all LSC snRNA-seq libraries, color-coded by disease status. The proportion of nuclei assigned to each cell type is shown as a percentage. The total number of nuclei per library is after quality control and doublet exclusion. Black bars mark median nuclei counts; asterisks denote significant differences between ALS and control donors (Welch’s t-test, p < 0.05). iii) Relative proportion of annotated cell types in LSC snRNA-seq libraries stratified by disease group. Asterisks denote significant differences between ALS and control donors (Welch’s t-test, p < 0.05). **C)** Same as (Bi–iii), but for MTC snRNA-seq data. **D)** Representative image of an annotated LSC Visium spatial transcriptomics array. Each spot in the LSC is given an annotated anatomical region (AAR) based on cellular composition, anatomical divisions, and H&E staining. **E)** i) UMAP plot of ST data. Each data point represents one spot from a Visium array, color-coded by AAR. ii) Number of spots annotated per spinal cord region across all Visium arrays, color-coded by disease status. The proportion of spots assigned to each region is shown as a percentage. The total number of spots per region is after quality control. Black bars mark median spots; asterisks denote significant differences between ALS and control donors (Welch’s t-test, p < 0.05). iii) Relative proportion of annotated Visium spots across spinal cord regions, stratified by disease group. Asterisks denote significant differences between ALS and control donors (Welch’s t-test, p < 0.05). **F)** Representative image of an annotated MTC Visium spatial transcriptomics array. **G)** Same as (Ei–iii), but for MTC ST data.

We similarly assigned seven broad CNS cell types to 190,367 high-quality MTC nuclei: EN, IN, OLIGO, OPC, AST, MIG, IMM, and ENDO (Figure 1Ci, Supplemental Figure 2). ALS donors showed a significant decrease in OPC number and proportion, along with an increased proportion of oligodendrocytes and a decreased proportion of excitatory neurons compared to controls (Figure 1Cii-iii).

We characterized the cytoarchitecture of the LSC and MTC across healthy and ALS-disease states with the 10x Genomics Visium platform (Figure 1A), generating spatially resolved (55 µm barcoded spots within a 6.5 × 6.5 mm array) transcriptomic maps. From 35 LSC donors (8 controls, 27 ALS) and 50 MTC donors (12 controls, 38 ALS), we sectioned multiple adjacent tissue sections for each donor (median of four arrays per donor), totaling 137 LSC and 183 MTC spatial transcriptomes for analysis.

From 137 LSC arrays, we captured 579,373 spatial spots, with an average of 2,130 unique RNA molecules and 909 genes per spot (Supplemental Figure 3). Each spatial spot was manually annotated with one of eleven anatomical regions based on hematoxylin and eosin (H&E) staining, cellular composition, and anatomical landmarks (Figure 1D). Preprocessed spatial spots were embedded into a UMAP plot, highlighting separation between the central canal (Cent_Can), where cerebrospinal fluid circulates, and the surrounding grey and white matter regions (Figure 1Ei). The distribution of spatial spots across LSC regions revealed a significant decrease in both the number and proportion of ventral horn spots in ALS donors compared to control donors, possibly reflecting a reduction in ventral horn area in disease (Figure 1Eii-iii).

From 183 MTC arrays, we captured 737,373 spatial spots, with an average of 4,756 unique RNA molecules and 1,914 genes per spot (Supplemental Figure 4A-F). Each spatial spot was manually annotated with one of five cortical layers (L1-6) or white matter based on H&E staining and cellular composition (Figure 1F, Supplemental Figure 4G). We did not include L4, as the MTC is classified as an agranular cortex^19^. Preprocessed spatial spots were embedded into a UMAP plot, highlighting separation between grey and white matter regions of the MTC (Figure 1Gi). There was no difference in the number or proportion of spatial spots across MTC regions between ALS and control donors (Figure 1Gii-iii). Two ALS donors showed notable separation in the UMAP embedding, one of which demonstrated high expression of heat shock-related genes, and the other showed elevated expression of immune response genes (Supplemental Figure 4H). These two donors were excluded from downstream analyses, along with one control donor who was diagnosed with multiple sclerosis and three donors that failed quality control, resulting in 8 control and 36 ALS donors for MTC spatial analysis.

### Glial changes surround the ventral horn in ALS

Given increased astrocyte abundance in our ALS LSC snRNA-seq data, and links between ALS and gliosis, we annotated glial cells for function using established marker genes. Astrocyte nuclei in the LSC were classified as residing in grey matter (GM) or white matter (WM) regions of the spinal cord based on their distinct transcriptomic signatures^20^. GM astrocytes regulate neuronal synapses and provide metabolic support, while WM astrocytes maintain and insulate myelinated axons^21^ (Figure 2A). Microglia were classified as homeostatic or reactive, or were re-annotated as other immune cells, including macrophages, monocytes, and lymphocytes^20,22^ (Figure 2B, Supplemental Figure 5). OPCs were classified into *OLIG1*^+^ differentiating states, *SOX10*^+^ myelin-promoting subpopulations^20,23,24^, or *NCOA1*^+^ reactive subpopulations (Figure 2C). Oligodendrocytes were divided into *OPALIN*^+^ or *RBFOX1*^+^ subpopulations, which are thought to represent intermediate and mature, stable states, respectively^25^ (Figure 2D).

**Figure 2.**
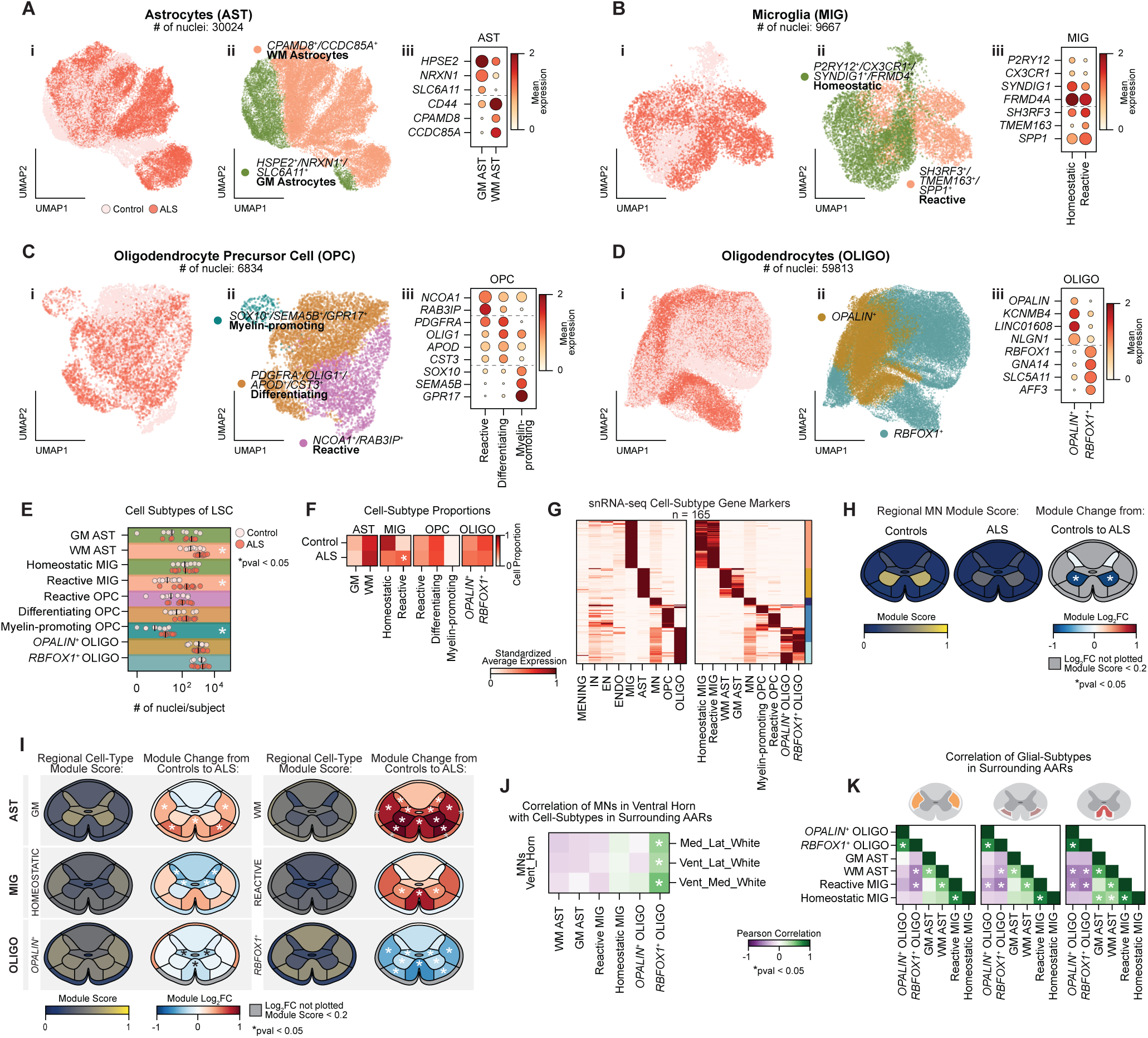
Glial changes surround the ventral horn in ALS. **A)** UMAP plot of astrocytes from snRNA-seq colored by i) disease status, ii) grey or white matter cell-subtype annotations, and iii) dot plot of marker genes. Dot plot is colored by mean gene expression and dot size denotes the proportion of cells with non-zero counts, from 0-100%. **B)** Annotation of microglia into homeostatic or reactive subtypes using the same plotting outline as A. **C)** Annotation of oligodendrocyte precursor cells into reactive, differentiating, or myelin-promoting subtypes using the same plotting outline as A. **D)** Annotation of oligodendrocytes into *OPALIN*^+^ or *RBFOX1*^+^ subtypes using the same plotting outline as A. **E)** Number of nuclei annotated per cell-subtype across all LSC snRNA-seq libraries, color-coded by disease status. Black bars mark median nuclei counts; asterisks denote significant differences between ALS and control donors (Welch’s t-test, p < 0.05, FDR-BH). **F)** Averaged proportions of cell-subtypes for controls and ALS. Asterisk indicates significant increase in reactive MIG in ALS donors relative to controls (Welch’s t-test, p < 0.05, FDR-BH). **G)** Min-maxed scaled expression of marker genes across LSC cell types (left) and cell-subtypes (right). **H)** Spatial expression of MN module scores across Visium arrays grouped by disease status (left and middle). Log_2_ fold change of MN module scores in ALS relative to controls (right). Asterisks indicate significant differences relative to control donors (Welch’s t-test, p < 0.05, FDR-BY). **I)** Spatial expression of glial subtype module scores across all Visium arrays (left). Log_2_ fold change of glial subtype module scores in ALS relative to controls (right). Asterisks indicate significant differences relative to control donors (Welch’s t-test, p < 0.05, FDR-BY). **J)** Pearson correlation matrix of glial subtype module scores in medial lateral white (top), ventral lateral white (middle), and ventral medial white (bottom) regions versus MN module scores in the ventral horn. Asterisks indicate a correlation significantly different from 0 (Wald test, p < 0.05, FDR-BH). **K)** Pearson correlation matrix of glial subtype module scores within medial lateral white (left), ventral lateral white (middle), and ventral medial white (right). Asterisks indicate a correlation significantly different from 0 (Wald test, p < 0.05, FDR-BH).

Among these subpopulations, we observed a significant increase in the number, but not the proportion, of both WM astrocytes and myelin-promoting OPCs in ALS donors, which we interpret as astrocyte and OPC proliferation (Figure 2E-F). This contrasts with an increase in both the number and proportion of reactive microglia, indicating a shift from homeostatic to reactive microglia in ALS (Figure 2E-F).

We next determined whether changes in cell subtype composition were global across the LSC or localized to distinct regions. We identified genes that were both preferentially expressed in each cell type and differentially expressed between its subpopulations (e.g. genes with higher expression in WM versus GM astrocytes) (Figure 2G, Supplemental Figure 6). For each set of genes, we calculated a functional module score (scaled average expression) for each spatial spot, allowing us to infer the spatial distribution and relative abundance of each cell subtype. For example, the MN module score was enriched in the ventral horn of control donors but significantly diminished in ALS (Figure 2H). GM and WM astrocyte module scores were predominantly expressed within the grey and white matter regions, respectively, but both populations were significantly increased in the white matter regions surrounding the ventral horn in ALS (Figure 2I). WM astrocyte scores were also increased within grey matter regions, particularly within the ventral horn, possibly reflecting WM astrocytes responding to localized atrophy or a shift in the functional state of GM astrocytes. Microglial module scores localize more strongly in white matter regions; in ALS donors, homeostatic microglial scores were significantly decreased in the dorsal horn and medial grey, which contrasts with reactive microglial scores that were significantly increased within the ventral horn and adjacent ventral medial white region (Figure 2I). *OPALIN*^+^ oligodendrocyte scores were present in both GM and WM, unlike *RBFOX1*^+^ oligodendrocyte scores that were predominantly localized to the WM regions. In ALS donors, there was a modest loss of *OPALIN*^+^ oligodendrocyte scores in medial grey and ventral medial white regions, but a significant decrease in *RBFOX1*^+^ oligodendrocyte scores within the ventral horn and surrounding regions (Figure 2I). Importantly, we did not detect any changes in the dorsal medial white region, which contains the sensory tracts, highlighting the specificity in these compositional changes. We conclude that alterations in cell subtype composition occur in spatially distinct regions rather than globally across the LSC.

Both a decrease in MN-associated gene expression in the ventral horn and changes in cell subtype composition in adjacent white matter regions are linked to ALS. These changes are not correlated in individual donors (Figure 2J), except for a within-donor correlation between MN module score and *RBFOX1*^+^ oligodendrocyte module score in the surrounding WM regions, indicating that these two cell types may be functionally linked. By contrast, within each WM region we find that reactive microglia scores correlate positively with WM astrocyte scores (but not GM astrocytes), and are anticorrelated with oligodendrocyte scores (Figure 2K). Together, these findings suggest that, at the donor level, there are two distinct patterns: a loss of lower MNs and *RBFOX1*^+^ oligodendrocytes in adjacent WM, and, separately, an increase in reactive microglia and astrocyte function that is linked to a decrease in oligodendrocyte function.

### LSC Spatial gene modules capture coordinated changes in ALS

Changes in MN and glial composition in the LSC are quantifiable because of our prior knowledge of these cell types and functions. To determine if there are other systematic changes in ALS, we fit a Bayesian hierarchical model for spatial transcriptomics, Splotch^26^, to our spatial transcriptomic data (Figure 3A). Splotch models gene expression at each spatial spot as a zero-inflated Poisson distribution, with spot-level gene expression λ informed by spot-and donor-level metadata (spinal cord region and site of symptom onset), spatial autocorrelation with neighboring spots, and spot-specific noise (Methods). Following Bayesian inference, we quantified both the magnitude (log_2_ fold-change) and significance (Bayes factor) of gene expression changes between control and ALS donors for each LSC region through analysis of posterior distributions over model parameters. We then calculate the correlation distance between modeled gene expression λ for each transcript, to construct a k-Nearest Neighbors (k-NN) graph, and cluster transcripts which are co-occurring together into 24 spatial gene modules (Figure 3A, Supplemental Figure 7). Modules M19 (chromosome M) and M23 (chromosome Y) contain transcripts which correlate from a single chromosome; the other modules are autosomal transcripts which have similar spatial co-expression patterns.

**Figure 3.**
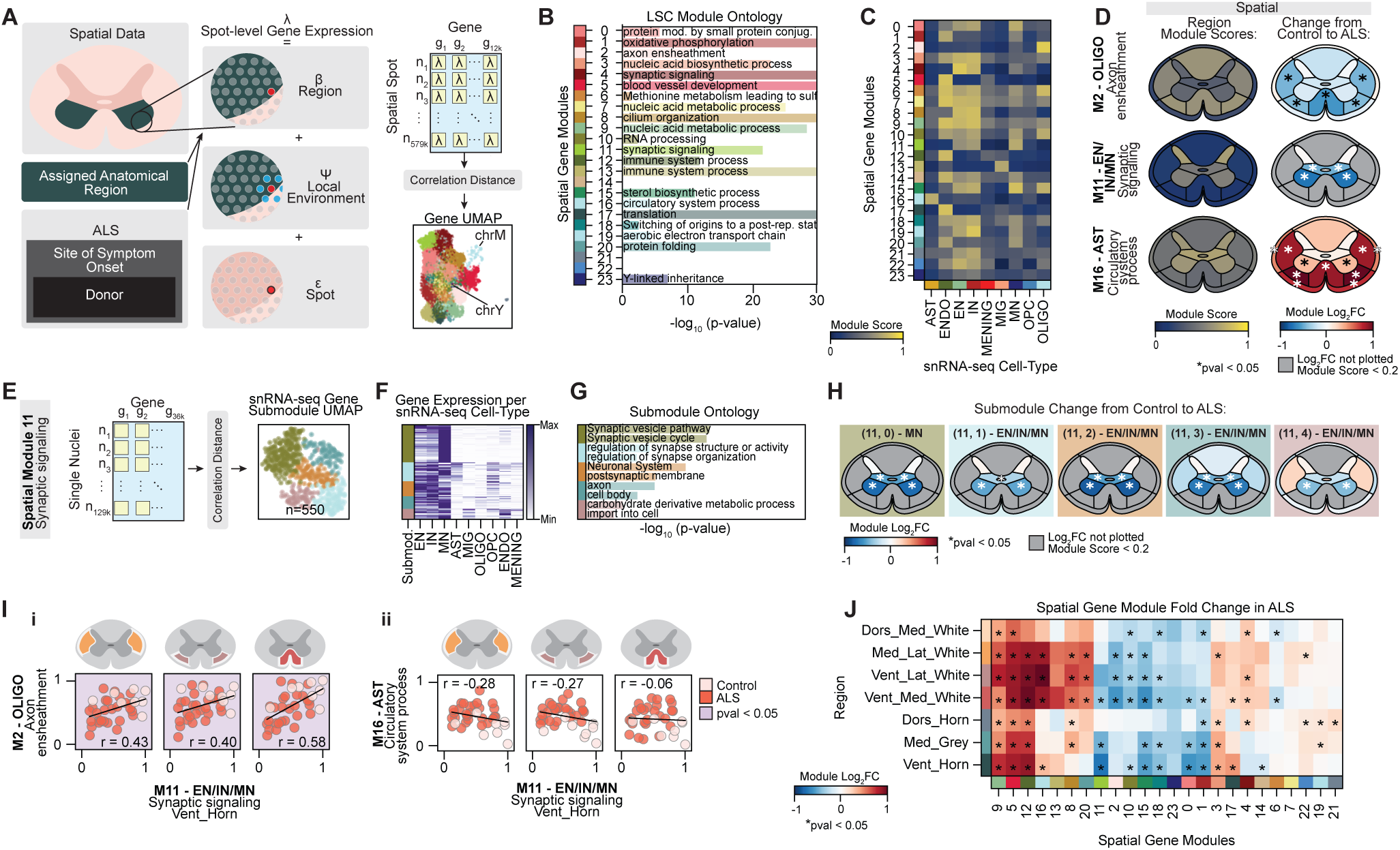
LSC Spatial gene modules capture coordinated changes in ALS. **A)** Schematic of the Splotch model for spatial transcriptomics data: spot-level gene expression (λ) is modeled as a combination of ALS disease state, site of symptom onset, and donor ID, specific region of interest (β), a spatial autoregressive component (ψ), and a spot noise (ε) component. Pearson correlation coefficients are calculated for all gene pairs across all spots. The distance between genes is derived from the correlation distance between modeled spot-level gene expressions (λ). Genes are embedded into a k-Nearest Neighbors (k-NN) graph, and clustered with the Leiden community clustering algorithm to identify spatially correlated gene modules. Gene-gene k-NN is embedded into a UMAP plot where each dot represents a gene, colored according to the assigned spatial gene module. **B)** Most significant gene ontology terms associated with each spatial gene module. Length of underlying bar plot indicates -log_10_(*p*-value). Empty lines indicate no enriched terms for that spatial gene module. **C)** Average spatial gene module score for each LSC snRNA-seq cell type. **D)** The average spatial gene module scores per spinal cord region (left). Log_2_ fold change of spatial gene modules in ALS donors relative to controls (right). Regions with module scores less than 0.2 are masked in log₂ fold change panels. Asterisks indicate significant differences in ALS relative to control donors (Welch’s t-test, p < 0.05, FDR-BY). **E)** Schematic demonstrating genes from spatial gene module 11 (M11) are assigned into submodules through correlation within snRNA-seq data. M11 gene-gene k-NN is embedded into a UMAP plot where each dot is a gene, colored according to the assigned submodule. **F)** Average expression of each gene in M11 for each LSC snRNA-seq cell type. Expression values are min-max scaled per gene for visualization, and submodule assignment is annotated on the left. **G)** GO terms associated with each submodule in M11, with length of underlying bar plot indicating -log_10_(*p*-value). **H)** Log_2_ fold change of submodule scores for M11 in ALS donors relative to controls. Regions with submodule scores less than 0.2 are masked in log₂ fold change panels. Asterisks indicate significant differences relative to control donors (Welch’s t-test, p < 0.05, FDR-BY). **I)** Donor-level correlation between M11 in the ventral horn and M2 (i) or M16 (ii) in surrounding white matter regions: medial lateral white (left), ventral lateral white (middle), and ventral medial white (right). Each dot represents an individual donor. Plots are annotated with least squares regression line, Pearson correlation coefficient, and shaded in violet if correlation is significantly different from zero (Wald test, p < 0.05, FDR-BH). **J)** Log_2_ fold change of LSC spatial gene modules in ALS relative to control donors, organized by similarity of changes. Asterisks indicate significant differences in ALS relative to control donors (Welch’s t-test, p < 0.05, FDR-BY).

Many of these spatial modules are enriched for transcripts annotated to gene ontology (GO) terms, or are linked to transcripts of a specific cell type (Figure 3B-C). Module M2 is linked to axonal ensheathment and module M16 is linked to circulatory system process, M4 and M11 are linked to synaptic signaling, and M12 and M13 are linked to immune response (Figure 3B). For each spatial spot, and for each snRNA-seq nuclei, we calculate a module score for each spatial module, which represents the scaled average expression of all transcripts in the module. For example, in single nuclei, oligodendrocytes showed the highest module score for M2, indicating they drive expression of genes associated with myelination, while astrocytes were enriched for genes of M16, likely reflecting the role of astrocytes in regulating blood flow^27^. Additionally, we found modules that were predominantly associated with endothelial cells (M12 and M17), microglia (M13), and neurons (M0, M4, and M11).

Many modules displayed distinct region-specific spatial expression patterns (Supplemental Figure 7C). For example, M2 and M11 were predominantly localized to the white and grey matter regions, respectively, whereas M16 was expressed diffusely across the LSC (Figure 3D). In our ALS donors, we observed a significant loss of M2 oligodendrocyte axonal ensheathment in regions surrounding the ventral horn, along with significant upregulation in M16 astrocyte circulatory system processes in the ventral horn and surrounding regions. We found a significant reduction in the neuronal-enriched M11 synaptic signaling module within the ventral horn (Figure 3D). Importantly, only M9 correlated with donor post-mortem interval. No other spatial gene modules or snRNA-seq module scores correlated with donor metadata (Supplemental Figures 8-9). These findings indicate that disease-associated changes are region-specific.

Each spatial spot can encompass a mixture of cell types, and spatial gene modules may reflect contributions from multiple cellular populations. We further divided our spatial modules into submodules by calculating separately for each module the Pearson correlation distance between each gene within our snRNA-seq expression data, constructing a k-NN graph, and then clustering the module genes into submodules (Figure 3E, Supplemental Figure 10). We were particularly interested in M11 as it was linked to genes enriched in MNs and was downregulated within the ventral horn of ALS donors. Upon dividing M11 into submodules, we found a MN-specific submodule (M11,0), while the other three submodules were broadly neuronal (Figure 3F). GO analyses showed that M11,0 is enriched for genes associated with the synaptic vesicle life cycle, while the other submodules are linked to synapse and axon structure, the postsynaptic membrane, and synaptic activity (Figure 3G). Each of the M11 submodules was decreased in the ventral horns of ALS donors (Figure 3H, Supplemental Figure 11).

At the donor level, downregulation of neuronal module M11 in the ventral horn significantly correlated with downregulation of oligodendrocyte module M2 in adjacent white matter, indicating coordinated impairments in synaptic signaling and axonal ensheathment (Figure 3Ii). However, it remains unclear whether these myelination impairments are a cause of synaptic deficits, a consequence of MN degeneration and the resulting loss of axonal substrate to myelinate, or reflect a combination of both processes. Notably, loss of M11 synaptic signaling was not significantly correlated with astrocyte module M16 in the nearby white matter, suggesting that, at this stage of disease, functional changes in synaptic signaling and astrocytes are not linked (Figure 3Iii).

While neuronal module M11 does not correlate with astrocyte module M16, we do find that two endothelial modules (M5 and M12) and a microglial module (M13) change similarly to M16 in ALS (Figure 3J). In addition, M11 and M2 cluster together and share patterns of change with M10 (RNA processing), M15 (sterol biosynthetic process), and M1 (oxidative phosphorylation). Taken together, these analyses show that the LSC in ALS undergoes many region-specific changes in cellular composition and function, with some of these changes occurring in a coordinated manner.

### Corticospinal tract loss correlates with glial and endothelial modules

Corticospinal tract (CST) loss is a hallmark of ALS pathology, and is clearly visible by the decrease in H&E staining in the medial lateral white matter (Figure 4A). Upper MN axons form the CST and enter the LSC through medial lateral white matter, synapsing onto lower MNs in the ventral horn, whose axons exit through ventral lateral white matter to innervate muscle; both upper and lower MNs and their axons are degraded in ALS.

**Figure 4.**
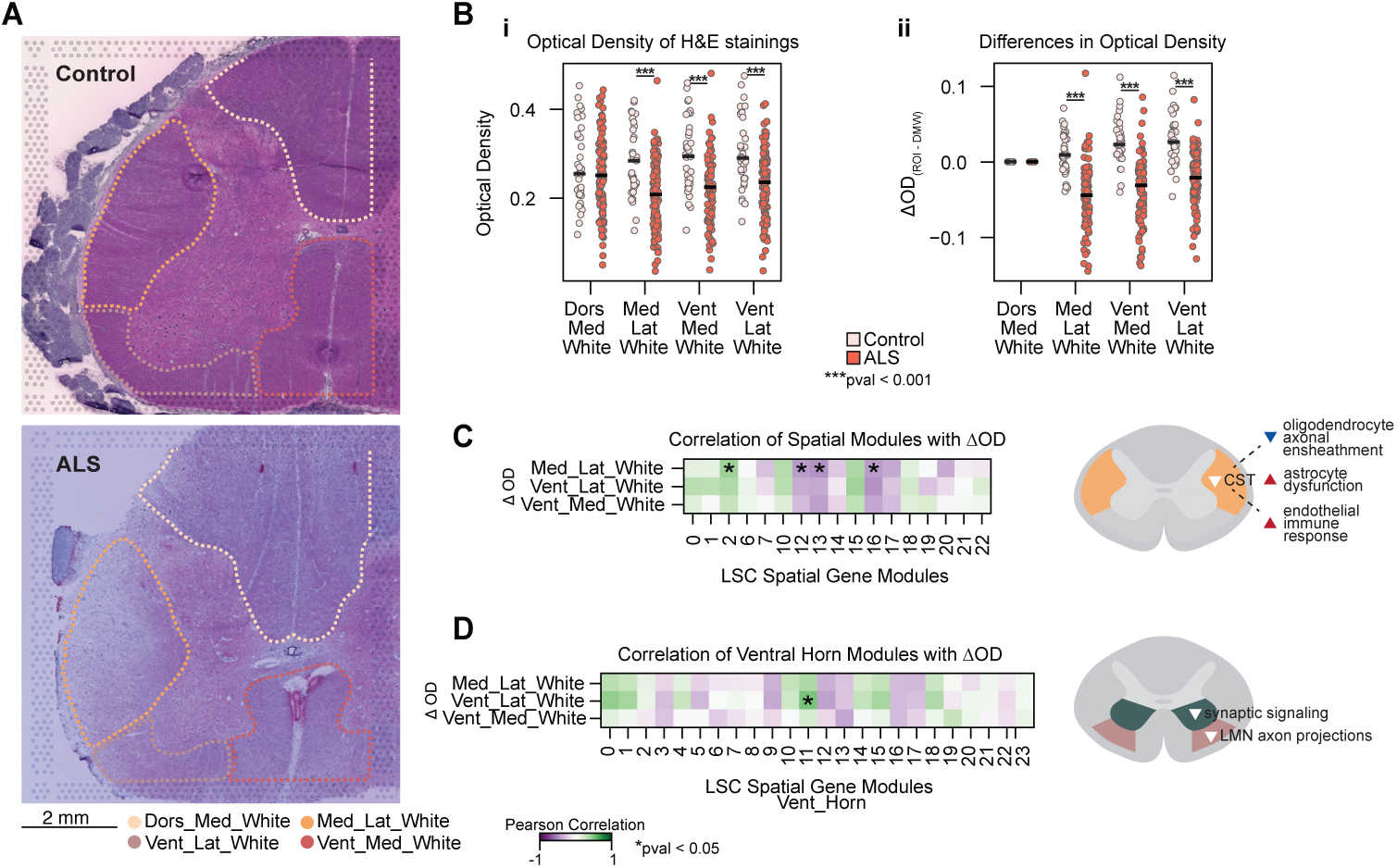
Corticospinal tract loss correlates with glial and endothelial modules. **A)** Representative H&E image demonstrating corticospinal tract loss with white matter AARs outlined. **B)** i) Average scaled optical density (OD) measurement per specified LSC anatomical region. ii) Average scaled OD relative to DMW, indicating the magnitude of OD loss for that region of interest (ROI). Each dot represents an individual array, colored by disease status. The black line indicates the median OD for each group. (Welch’s t-test, p < 0.05, FDR-BY). **C)** Pearson correlation matrix of spatial gene module scores within medial lateral white, ventral lateral white, and ventral medial white and their corresponding ΔOD. Asterisks indicate a correlation significantly different from 0 (Wald test, p < 0.05, FDR-BH). **D)** Pearson correlation matrix of spatial gene module scores in the ventral horn and the ΔOD within medial lateral white, ventral lateral white, and ventral medial white. Asterisks indicate a correlation significantly different from 0 (Wald test, p < 0.05, FDR-BH).

To quantify this loss in the H&E stained tissue sections used for Visium spatial transcriptomics, we measured the optical density (OD) for the medial lateral, ventral lateral, and ventral medial white matter regions which we expect to be degraded in ALS, as well as the dorsal medial white matter, which contains the sensory tracts and is relatively unaffected in ALS. OD in the dorsal medial white matter did not differ between control and ALS donors, whereas OD in the medial lateral, ventral lateral, and ventral medial white matter regions was significantly reduced in ALS (Figure 4Bi). This effect is more pronounced when we control for differences in staining by calculating ΔOD, relative to dorsal medial white matter staining in each section. This decrease in ΔOD reflects a loss of CSTs and lower MN axons (Figure 4Bii), and does not correlate with disease duration, age at death, or postmortem interval (Supplemental Figure 12).

When we compared CST loss, quantified by ΔOD in the medial lateral white matter regions, to the spatial modules with high expression in white matter regions, we find a significant correlation with oligodendrocyte-myelination spatial module M2, likely reflecting a decrease in myelination associated with CST loss. This contrasts with endothelial and microglia-immune response spatial modules M12 and M13, and astrocyte spatial module M16 which are anticorrelated with ΔOD, such that expression of these modules increases with CST loss (Figure 4C). In addition, ΔOD in the ventral lateral white matter region is significantly correlated with a transcriptomic loss of spatial module M11 in the ventral horn. We interpret this as the loss of synaptic signaling in the ventral horn and the loss of lower MN axons in the ventral lateral white matter are linked (Figure 4D).

### Site of symptom onset influences MN and endothelial functions

Given that MN loss is greater in the regions proximal to site of symptom onset^5^, we examined whether ALS-associated changes in the LSC are more severe in limb-versus bulbar-onset donors. We found a significant decrease in MN-associated gene expression in the ventral horn of limb-onset donors compared to bulbar-onset donors (Figure 5A). We directly quantified the number of MNs in the ventral horn from the Visium H&E images, finding significantly fewer MNs in limb-compared to bulbar-onset donors, with the number of MNs per donor correlating with MN-associated gene expression (Figure 5Bi-ii, Supplemental Figure 13A). Critically, neither measure correlated with disease duration, indicating that greater MN loss is related to symptom onset (Figure 5Biii, Supplemental Figure 13B-C). In contrast, site of onset had minimal influence on the spatial composition of astrocyte, microglia, or oligodendrocyte subpopulations in the LSC (Supplemental Figures 13D). Together, these findings suggest that site of symptom onset influences the severity of MN loss, but not necessarily glial response.

**Figure 5.**
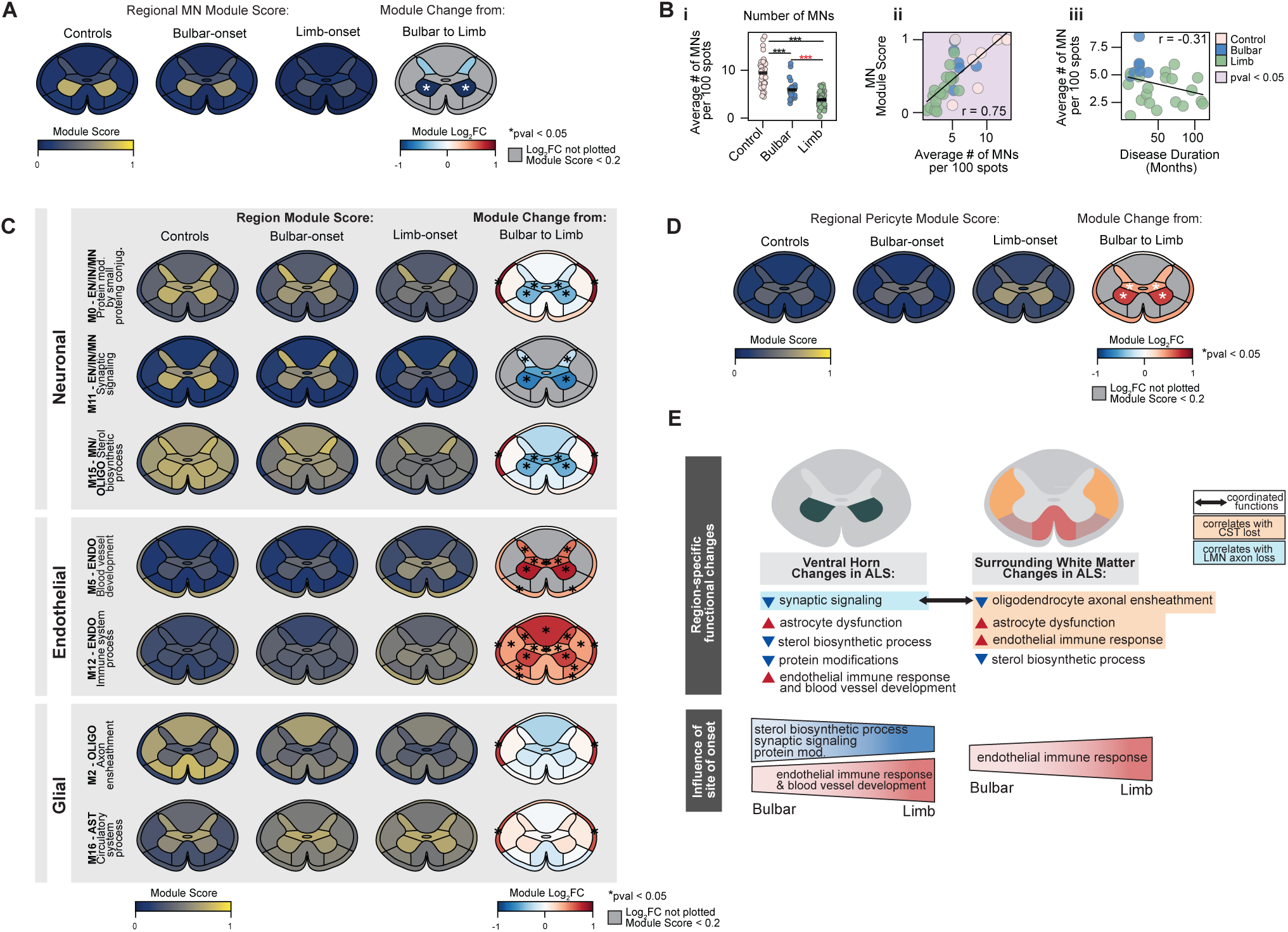
Site of symptom onset influences motor neuron and endothelial functions. **A)** Spatial expression of MN gene module scores across Visium arrays, grouped by site of symptom onset. Log_2_ fold change of MN module scores in limb-onset donors relative to bulbar-onset donors (right). Asterisks indicate significant differences in limb-relative to bulbar-onset donors (Welch’s t-test, p < 0.05, FDR-BY). **B)** i) Number of MNs per 100 spots in the ventral horn for each Visium array. The black lines indicate the mean for each group. (Welch’s t-test, p < 0.05). ii) Donor-level correlation between the average ventral horn MN count and the average MN gene module score. iii) Donor-level correlation between disease duration and average ventral horn MN count. Each dot represents an individual donor. Plots are annotated with least squares regression line, Pearson correlation coefficient, and shaded in violet if correlation is significantly different from zero (Wald test, p < 0.05). **C)** Spatial expression of neuronal-, endothelial-, and glial-enriched spatial gene module scores across Visium arrays grouped by site of symptom onset. Log_2_ fold change of spatial gene modules in limb-onset relative to bulbar-onset donors. Regions with module scores less than 0.2 are masked in log₂ fold change panels. Asterisks indicate significant differences relative to bulbar-onset donors (Welch’s t-test, p < 0.05, FDR-BY). **D)** Spatial expression of pericyte module score across Visium arrays grouped by site of symptom onset. Log_2_ fold change of pericyte module scores in limb-onset relative to bulbar-onset donors. Regions with module scores less than 0.2 are masked in log₂ fold change panels. Asterisks indicate significant differences relative to bulbar-onset donors (Welch’s t-test, p < 0.05, FDR-BY). **E)** LSC summary schematic highlighting region-specific functional changes, coordinated alterations, and functions influenced by site of symptom onset.

Several spatial gene modules associated with neuronal and grey matter expression are decreased in limb-onset donors, compared to bulbar-onset. Neuronal modules M0 and M11 (synaptic signaling) were significantly decreased in grey matter, as was neuronal module M15 (sterol biosynthetic process). M15 expression is decreased in both limb- and bulbar-onset donors across grey and white matter, but is decreased further in grey matter in limb-onset donors (Figure 5C, Supplemental Figure 14). Notably, endothelial spatial gene modules M5 and M12, associated with blood vessel development and immune system processes, were significantly upregulated in ALS, with greater increases observed in limb-onset relative to bulbar-onset. In contrast, glial modules such as M2 (oligodendrocyte axonal ensheathment) and M16 (astrocyte circulatory system processes) did not differ between bulbar- and limb-onset donors (Figure 5C). Among endothelial-associated cell types, we observed a significant increase in pericyte-associated gene expression in the ventral horn of limb-onset donors compared to bulbar-onset donors, which did not correlate with disease duration (Figure 5D). Together, these findings suggest that MN and endothelial response is influenced by the proximity to the site of symptom onset, whereas glial functions remain largely unaffected (Figure 5E).

### Synaptic signaling-associated genes are increased in L5 of MTC in ALS

Following the characterization of ALS-linked cell type- and region-specific changes in the LSC, we assessed whether these alterations are shared with the MTC. To facilitate the analysis of neurons, particularly rarer subtypes like L5 extratelencephalic neurons, we integrated our snRNA-seq dataset with a publicly available MTC snRNA-seq dataset^25^, resulting in 564,885 MTC nuclei (Figure 6A and Supplemental Figure 15). We then classified MTC astrocytes and microglia as reactive or homeostatic, classified OPCs as *OLIG1*^+^ maturing, *NRXN1*^+^ homeostatic, or *NCOA1*^+^ stressed/reactive^28^, and classified oligodendrocytes as *OPALIN*^+^, *RBFOX1*^+^, or low-myelinating *PLP1^-^* (Figure 6A). MTC cell populations differed substantially from similar cell populations in the LSC; in astrocytes, changes to gene expression and chromatin accessibility are specific for both tissue and function (Supplemental Figure 16). A small proportion of endothelial nuclei were classified as pericytes. Among these glial subpopulations, we observed a significant increase in the number, but not the proportion, of reactive astrocytes and *OPALIN*^+^ oligodendrocytes in ALS donors. (Supplemental Figure 17A-B).

**Figure 6.**
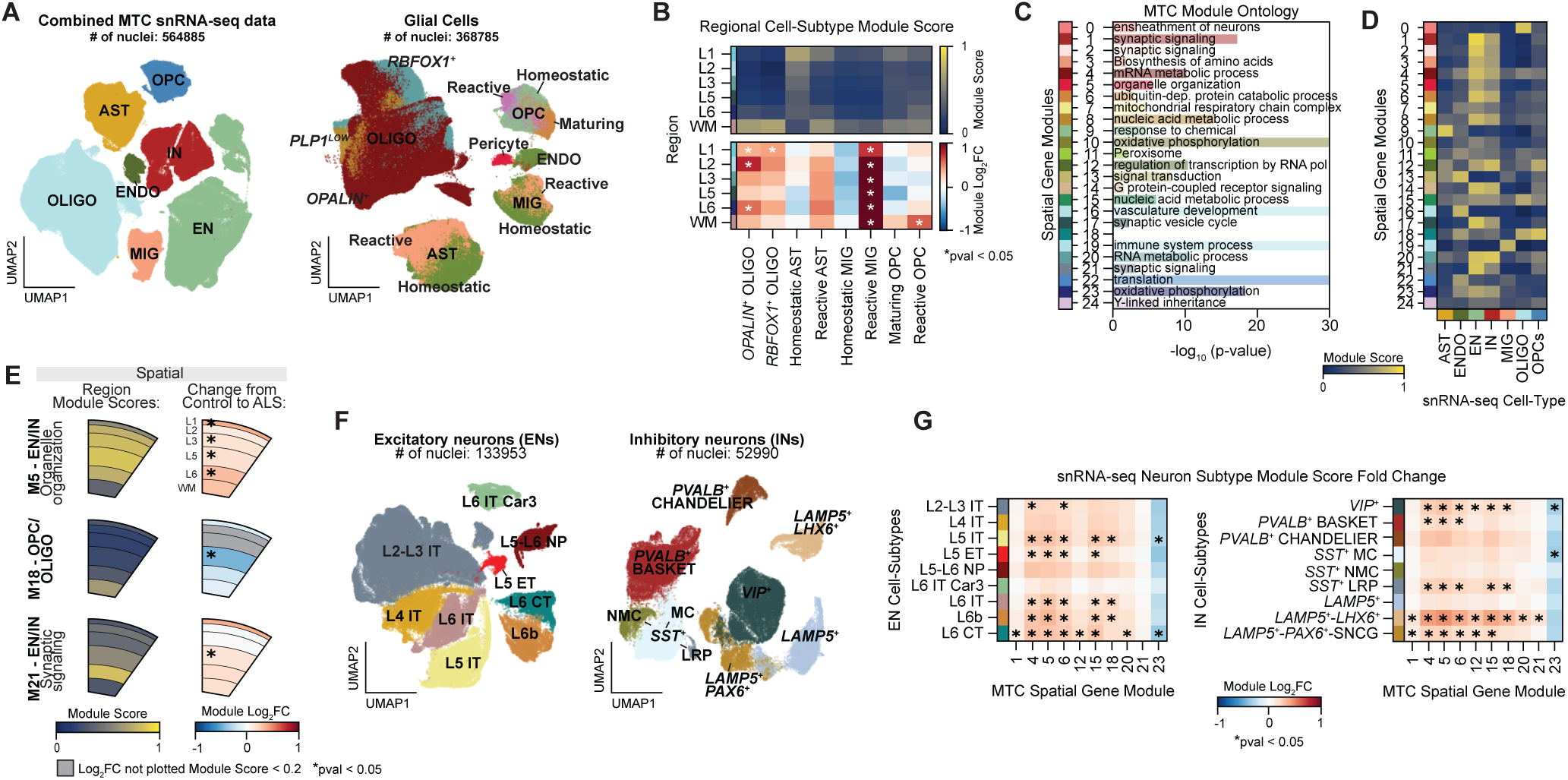
Synaptic signaling is increased in L5 of MTC in ALS. **A)** UMAP plot of cell types and non-neuronal cell-subtypes from integrated MTC snRNA-seq data. Each data point represents one nucleus, color-coded by cell type or subtype annotation. **B)** Spatial expression of glial subtype module scores across all MTC Visium arrays (top). Log_2_ fold change of glial subtype module scores in ALS relative to controls (bottom). Asterisks indicate significant differences relative to control donors (Welch’s t-test, p < 0.05, FDR-BY). **C)** Most significant gene ontology terms associated with each MTC spatial gene module. Length of underlying bar plot indicates -log_10_(*p*-value). Empty lines indicate no enriched terms for that spatial gene module. **D)** Average spatial gene module score for each cell type from integrated MTC snRNA-seq data. **E)** The average spatial gene module scores per MTC region (left). Log_2_ fold change of spatial gene modules in ALS donors relative to controls (right). Regions with module scores less than 0.2 are masked in log₂ fold change panels. Asterisks indicate significant differences relative to control donors (Welch’s t-test, p < 0.05, FDR-BY). **F)** UMAP plot of EN and IN subtypes from integrated snRNA-seq data. Each data point represents one nucleus, color-coded by neuronal subtype. **G)** Log_2_ fold change of selected spatial gene modules scores for each MTC snRNA-seq neuronal subtype, comparing ALS and control donors. Asterisks indicate significant differences from controls (Welch’s t-test, p < 0.05, FDR-BH).

We used these classified MTC snRNA-seq nuclei to identify glial subtype-specific gene sets, and scored MTC spatial transcriptomes with these subtype modules for analysis (Supplemental Figure 17C). In ALS donors, there is a significant increase in reactive microglia-associated gene expression throughout the MTC, and a significant increase in genes associated with stressed/reactive OPCs in the white matter. In contrast to the decrease seen in the LSC, genes associated with *RBFOX1*^+^ and *OPALIN*^+^ oligodendrocytes were significantly increased in specific layers of the MTC in ALS (Figure 6B).

We applied the Splotch model and the spatial gene module and submodule method used for LSC to the MTC, identifying 25 spatial co-expression MTC gene modules (M_MTC_), with M_MTC_23 and M_MTC_24 corresponding to mitochondrial- and Y-chromosome gene expression (Supplemental Figures 18-20). GO analysis shows modules are linked to synaptic signaling (M_MTC_1, M_MTC_2, and M_MTC_21), myelination (M_MTC_0), and various metabolic processes (M_MTC_4, M_MTC_8, M_MTC_10, M_MTC_15, and M_MTC_20) (Figure 6C). Using the integrated MTC snRNA-seq dataset, we assessed whether spatial gene modules were enriched by specific cell types, identifying spatial modules associated with oligodendrocytes (M_MTC_0), endothelial cells (M_MTC_16), astrocytes (M_MTC_9), microglia (M_MTC_19), and neurons (Figure 6D).

Some MTC spatial gene modules were broadly expressed across all spatial layers, and some showed spatially restricted patterns (Supplemental Figure 18C). For example, neuronal-enriched M_MTC_5 (organelle organization) was prominent in grey matter, while M_MTC_18 is specific for white matter and M_MTC_21 is enriched in L6 (Figure 6E). Some spatial modules were significantly increased in ALS donors across all cortical layers, such as M_MTC_5, whereas others showed layer-specific changes (Figure 6E). In contrast to LSC, synaptic signaling-associated M_MTC_21 was significantly increased in L5. Importantly, only M_MTC_17 correlated with donor post-mortem interval across the MTC, suggesting that these region-specific changes are driven by disease rather than sample quality (Supplemental Figure 21).

We leveraged our integrated snRNA-seq data to determine whether specific neuronal subtypes contributed to changes in MTC spatial gene module expression in ALS. From the snRNA-seq data, we identified 133,953 excitatory and 52,990 inhibitory nuclei, which were classified into subtypes using key marker genes (Figure 6F, Supplemental Figure 22). ENs were classified into nine subtypes, including L2/3, L4, L5, L6, and L6 Car3 intratelencephalic (IT) neurons, L5 *VAT1L*^+^ extratelencephalic (ET) neurons^29,30^, L5/L6 near-projecting (NP) neurons, L6 corticothalamic (CT) projection neurons, and L6b neurons^28^. INs were also classified into nine subtypes, including *VIP*^+^, *PVALB*^+^ (basket and chandelier), three *LAMP5*^+^ subtypes, and three *SST*^+^ interneuron subtypes. *SST*^+^ clusters were further annotated as Martinotti (MC), non-Martinotti (NMC), and long-range projection (LRP) interneurons^28,31^. We did not observe differences in the number or proportion of EN subtypes between ALS and control donors, but did find many differences in IN subtypes; we observed a significant increase in the number of *VIP*^+^, *PVALB*^+^ Basket, and *LAMP5*^+^ INs in ALS, while both the number and proportion of *SST*^+^ MC INs increased in ALS (Supplemental Figure 22C-F). We also detected a significant decrease in the proportion in *PVALB*^+^ chandelier INs in ALS (Supplemental Figure 22F). In addition, among the MTC spatial gene modules that were significantly altered across one or more cortical layers in ALS, we found that synaptic signaling–associated M_MTC_21 was significantly increased in *LAMP5*⁺–*LHX6*⁺ interneurons, suggesting that this subpopulation drives the L5–specific increase (Figure 6J). Importantly, we confirmed that snRNA-seq module scores in both ENs and INs were driven by disease and not age (Supplemental Figure 23).

### MTC and LSC have distinct molecular signatures in ALS

Our cohort includes 18 donors (4 controls, 14 ALS) with spatial profiling of both the MTC and LSC, enabling comparisons within individual donors. This allows us to test whether white matter degeneration in the LSC is associated with gene expression changes in the MTC, by using the average LSC ΔOD measurements of the medial lateral, ventral lateral, and ventral medial white matter regions as a predictor in a DESeq2 model applied to our MTC spatial data. We identified 206 genes, including *SOD1*, that showed increased expression and 63 genes that showed decreased expression in association with decreases in ΔOD (Figure 7A). The upregulated MTC genes are linked to upper MN dysfunction, macroautophagy, protein maturation, and synapse organization while the downregulated genes are associated with transmembrane transport (Figure 7B). In MTC snRNA-seq data, most of the upregulated genes are primarily expressed by neurons; in the MTC spatial data, these genes are predominantly expressed in the grey matter and are significantly increased throughout the cortical layers in ALS donors (Figure 7C). While these genes are also expressed in the grey matter of the LSC, they are decreased in expression in ALS donors, opposite to the upregulation seen in the MTC (Figure 7C).

**Figure 7.**
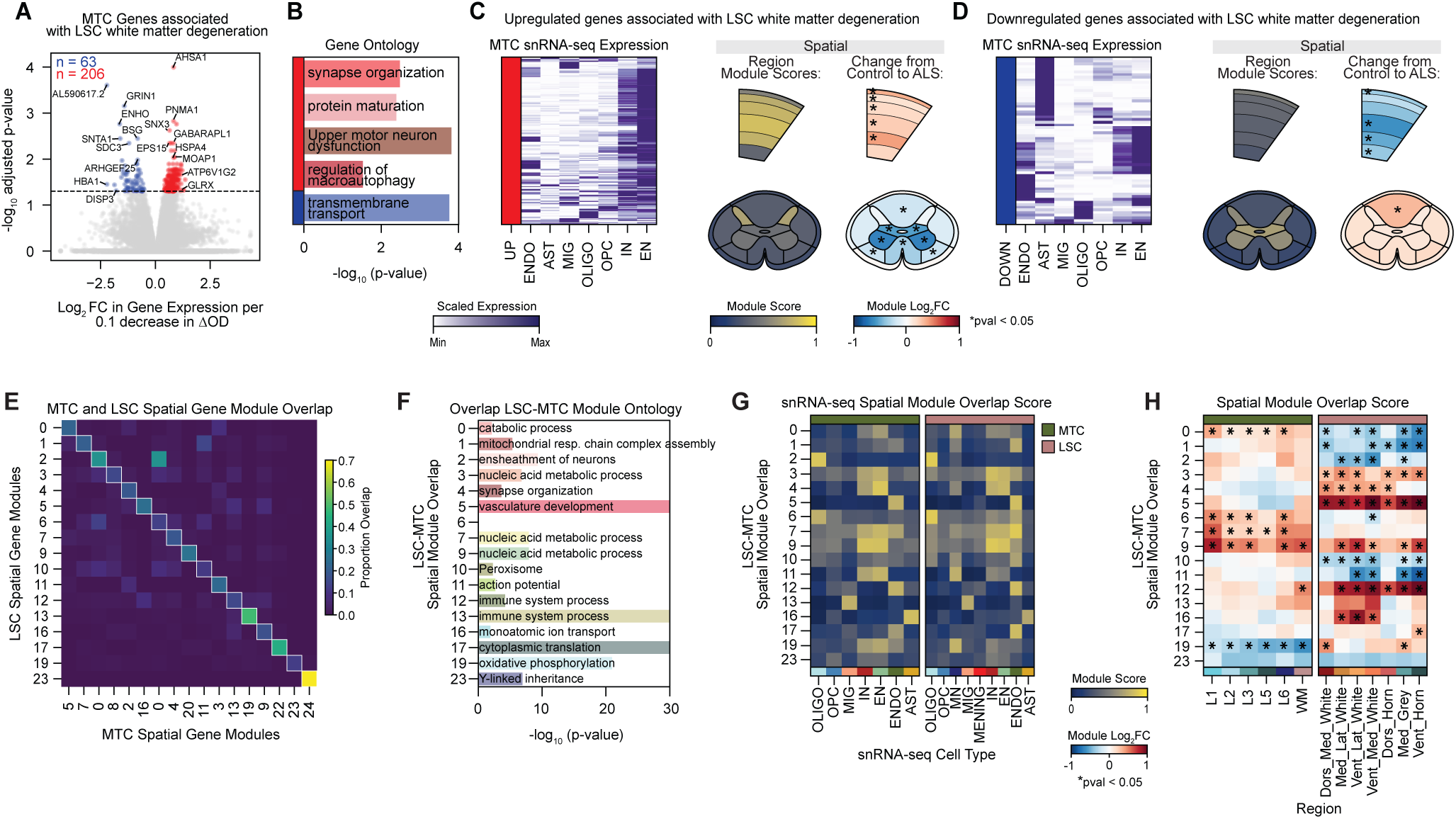
MTC and LSC have distinct molecular signatures in ALS. **A)** Volcano plot showing genes from MTC spatial data associated with decreases in LSC ΔOD. The x-axis represents the change in MTC gene expression per 0.1 decrease in ΔOD. Genes in red are predicted to increase in expression with the loss of ΔOD, while genes in blue decrease with the loss of ΔOD. (DESeq2, multiple hypothesis test corrected, Wald test, p < 0.05) **B)** Selected gene ontology terms associated with upregulated (red) and downregulated (blue) genes, with bar length representing -log_10_(*p*-value). **C)** Min-max scaled expression of upregulated genes across MTC snRNA-seq cell types (left). Average module score for upregulated genes across MTC and LSC regions, including all arrays (not limited to donors with paired MTC and LSC ST data) (right). Log_2_ fold change of upregulated genes in ALS donors relative to controls (far right). Asterisks indicate significant differences relative to control donors (Welch’s t-test, p < 0.05, FDR-BY). **D)** Same as (C), but for downregulated genes. **E)** Each square represents the proportion of genes shared between each LSC and MTC spatial module. For each row, a white box highlights the module pair with the greatest gene overlap, defined by a threshold of at least 0.1. For example, LSC M2 corresponds to MTC M0 (332 shared genes). Module pairs are referred to using LSC nomenclature hereafter. **F)** Most significant gene ontology terms for genes shared between paired LSC and MTC spatial modules. Length of underlying bar plot indicates -log_10_(*p*-value). Empty lines indicate no enriched terms for that spatial gene module. **G)** Average module score of genes shared between paired LSC and MTC spatial modules, calculated for each snRNA-seq cell type in MTC (left) and LSC (right). **H)** Log_2_ fold change in module scores for genes shared between paired LSC and MTC spatial modules in ALS donors relative to controls, shown for MTC (left) and LSC (right). Analysis was not limited to donors with paired MTC and LSC ST data. Asterisks indicate significant differences relative to control donors (Welch’s t-test, p < 0.05, FDR-BY).

In contrast, the 63 genes whose expression decreased as ΔOD decreased were expressed by astrocytes, endothelial cells, and neurons (Figure 7D). These downregulated genes were expressed diffusely across the MTC, significantly decreasing the most in L5 in ALS donors. In the LSC, these genes were expressed in the grey matter and showed little change in ALS (Figure 7D). While these findings are correlative, and they do not establish a mechanistic link, we interpret them as an association between white matter degeneration in the LSC and neuronal and glial dysfunction in the MTC. The opposing expression patterns observed between the LSC and MTC suggests that molecular changes in ALS are region-specific.

To compare changes across regions, we calculated the proportion of genes shared between each LSC and MTC spatial module and identified the module pairs with the greatest overlap (Figure 7D). For example, LSC M2 exhibited the greatest gene overlap with M_MTC_0, sharing 332 genes. For simplicity, overlapping modules are hereafter referred to by their LSC module number. We investigated the functional significance of these shared genes using GO enrichment analysis. The resulting GO terms largely recapitulated the associations of the individual LSC and MTC modules – e.g., both LSC M2 and MTC M_MTC_0 are associated with similar biological processes, including ensheathment of neurons (Figure 7E). To assess whether these shared spatial gene modules are driven by similar cell types, we calculated module scores for the shared genes in our LSC and MTC snRNA-seq data. We found that these gene modules are largely expressed in the same cell types across both regions. For example, M2 is enriched in oligodendrocytes, M13 in microglia, and M16 in astrocytes (Figure 7F). However, spatial analysis revealed distinct regional expression patterns of these shared modules in LSC and MTC in ALS (Figure 7G). Despite being expressed in similar cell types and involved in comparable functional pathways, these genes exhibit markedly distinct changes in expression in ALS donors. These findings suggest that ALS-linked functional changes are different between the spinal cord and motor cortex.

## DISCUSSION

Our study characterizes the cellular composition and cytoarchitecture of the LSC and MTC in both healthy and ALS disease states by combining single-nucleus and spatial transcriptomic profiles to construct spatially-resolved gene expression maps. We find regional specificity in altered functions in ALS, as exemplified by a gradient loss of synaptic signaling (M11), with no change in the dorsal horn, a modest decrease in the medial grey, and a pronounced decrease in the ventral horn. This pattern indicates that sensory neurons in the dorsal horn are relatively spared from synaptic signaling alterations, whereas MNs of the ventral horn are particularly vulnerable. This affirms prior work showing synaptic density is preserved in the dorsal regions in the LSC of ALS donors, but significantly decreased in ventral regions^32^. Synaptic signaling deficits in the ventral horn likely reflect a combination of lower MN degeneration and the loss of upper MN input.

We find that loss of synaptic signaling-associated genes (M11) in the ventral horn is correlated with the loss of axonal ensheathment (M2) genes in adjacent white matter. However, it remains unclear whether myelination impairments are a cause of synaptic deficits, a consequence of MN degeneration and the resulting loss of axonal substrate to myelinate, or reflect a combination of both processes. Notably, loss of synaptic signaling did not correlate with changes in the expression of astrocyte-enriched genes associated with circulatory system process (M16) in the nearby white matter. This lack of association may be due to the specific functional profile captured by M16. Alternatively, synaptic signaling and this astrocyte function may have been associated at earlier stages of disease progression, but this association may no longer be detectable at end-stage.

Our cohort was purposefully designed to compare ALS pathology in the LSC between limb-onset and bulbar-onset donors. We restricted our site of symptom onset analyses to the LSC, as its columnar anatomical organization allows for comparable regions across donors. We show that MN-associated gene expression in the ventral horn is reduced in bulbar-onset donors, and recapitulate the finding^5^ that the number of MNs in the ventral horn is greatly reduced in limb-onset donors relative to bulbar-onset donors. In contrast, vascular- and immune-linked modules (M5 and M12) are increased in ALS donors in general, and are significantly increased in limb-onset donors relative to bulbar-onset donors. We hypothesize that this reflects an angiogenic response to MN atrophy in the ventral horn, which is more pronounced in limb-onset donors, and an immune response associated with the disruption of the blood spinal cord barrier. Increased microvascular density has been observed in the LSC grey matter of ALS donors, suggesting a compensatory angiogenic response to MN degeneration or vascular deficits^31–33^. Additionally, endothelial and pericyte dysfunction in ALS increases blood spinal cord barrier permeability, allowing entry of normally excluded components such as immunoglobulin G and lymphocytes, contributing to a local immune response^14,33–36^.

Astrocyte- and oligodendrocyte-enriched modules in the LSC are increased in ALS donors, but are similar between bulbar- and limb-onset donors. As decreased gray matter neuronal gene expression (M11) correlated with decreased axonal ensheathment gene expression in adjacent white matter (M2), we expected more loss of axonal ensheathment expression in limb-onset donors. However, we do not see this, and found no difference in the loss of lower MN axons (as measured by ΔOD in ventral lateral white matter) between limb- and bulbar-onset donors. This suggests that deficits in oligodendrocyte-mediated myelination may occur earlier and more uniformly across donors, possibly preceding regionally distinct MN loss and dysfunction, and SOD1^G93A^ mouse models of ALS have shown that oligodendrocyte degeneration and reduced expression of myelin-related genes precede disease onset^37,38^.

In contrast to the LSC, the MTC exhibits a L5-specific increase in synaptic signaling (M_MTC_21) that is driven by *LAMP5*⁺–*LHX6*⁺ INs. This is notable given the cortical hyperexcitability observed in ALS donors and the vulnerability of MNs to excitotoxicity^39–42^. Although *LAMP5*⁺–*LHX6*⁺ INs remain incompletely characterized in humans, they are abundant in primate cortices, including the human MTC, and are primarily localized to deep cortical layers^43^. In primates, these cells most closely resemble mouse hippocampal *LAMP5*⁺–*LHX6*⁺ INs, particularly neurogliaform cells^43–45^. In human cortical slices, neurogliaform cells evoke slow, long-lasting inhibition in postsynaptic cells^46^. It is possible that increased expression of synaptic signaling-related genes in L5 *LAMP5*⁺–*LHX6*⁺ INs reflects a compensatory process to modulate hyperexcitation.

Comparison of the molecular and cellular changes in the LSC and MTC shows that ALS-associated changes in these modules differ markedly between the LSC and MTC. The astrocyte-linked spatial module (M16) increases in ALS donors in the white matter of the LSC, but does not change in the MTC. In support of these region-specific changes, a previous study^47^ found that inflammatory glial signatures occurred more frequently in the spinal cord than in the MTC of ALS donors with matched tissues (71% vs 10% of cases, respectively). Notably, the neuronal-linked spatial modules (M0) increases across cortical layers in the MTC but decreases in grey matter regions of the LSC and contains genes broadly associated with catabolic processes, including protein homeostasis and macroautophagy. These findings converge with the matched LSC ΔOD and MTC analysis, where neuronal-enriched genes linked to protein homeostasis and macroautophagy increase in the MTC in association with white matter degeneration in the LSC. Previous studies have also reported upregulation of protein homeostasis-associated genes in excitatory neurons of the MTC in ALS donors^29,48^.

In summary, ALS-associated changes differ markedly between the LSC and MTC, suggesting region-specific functional changes in disease. These differences, in part, may reflect the distinct tissue environments: the MTC is primarily grey matter with heterogenous neuronal populations, whereas the LSC is predominantly white matter enriched in glia. Consequently, cell types may be differentially affected during disease progression, resulting in the distinct end-stage molecular signatures we observe.

## METHODS

### Tissue sourcing and donor information

Human post-mortem lumbar spinal cord and motor cortex tissue were acquired through the ALS Consortium and TargetALS initiatives. Samples were collected in accordance with institutional review board (IRB) protocols from member sites and were transferred to NYGC following all applicable regulations for processing, sequencing, and analysis. Metadata for donors in this study are in Supplemental Table 1.

### Nuclei isolation from frozen post-mortem tissue

Nuclei were isolated from fresh frozen post-mortem motor cortex and spinal cord samples using a protocol modified from previously reported methods^28,49,50^. All steps were performed on ice unless stated otherwise. Tissue blocks were stored at -80°C prior to processing and were placed on dry ice for 20-30 minutes to equilibrate before being transferred to a cryostat at -16°C. Tissue blocks equilibrated in cryostat for 15-20 minutes before sectioning. To profile nuclei from the same representative plane used in our Visium experiments, approximately 5-10 50 µm sections of tissue were collected into pre-chilled 1.5 mL Eppendorf tubes. Each sample received 700 µL of homogenization buffer (320 mM sucrose, 5 mM CaCl_2_ [Millipore Sigma; #21115], 3 mM magnesium acetate [Thermo Fisher; AAJ60041AE], 10 mM Tris-HCl pH 7.8 [prepared from Thermo Fisher’s #AM9850G and #AM9855G], 0.1 mM EDTA pH 8 [Fisher; 15-575-020], 0.1% IGEPAL CA-630 [Millipore Sigma, I8896-50ML], 1 mM DTT [Millipore Sigma, 646563-10X], 0.4 U/µL RNAse inhibitor [Millipore Sigma, 03335399001], H_2_O) and incubated on ice for 5 minutes. The samples were transferred to 1 mL Dounce tissue grinders (DWK Life Sciences, 357538) and homogenized using ∼10-15 strokes with the loose pestle followed by ∼10-15 strokes with the tight pestle. After an additional 5 minute incubation on ice, the homogenized samples were filtered through a 70 µm strainer (Bel-Art, H13680-0070) into 2 mL Eppendorf tubes. The filtered samples were mixed with an equal volume of working solution (50% OptiPrep [STEMCELL Technologies, #07820], 5 mM CaCl_2_, 3 mM magnesium acetate, 10 mM Tris-HCl pH 7.8, 0.1 mM EDTA pH 8, 1 mM DTT). In a 15 mL Falcon tube, 750 µL of 30% OptiPrep solution (30% OptiPrep, 134 mM sucrose, 5 mM CaCl_2_, 3 mM magnesium acetate, 10 mM Tris-HCl pH 7.8, 0.1 mM EDTA pH 8, 0.04% IGEPAL CA-630, 1 mM DTT, 0.17 U/µL RNAse inhibitor) was layered directly over 300 µL 40% OptiPrep solution (40% OptiPrep, 96 mM sucrose, 5 mM CaCl_2_, 3 mM magnesium acetate, 10 mM Tris-HCl pH 7.8, 0.1 mM EDTA pH 8, 0.03% IGEPAL CA-630, 1 mM DTT, 0.12 U/µL RNAse inhibitor). The filtered and mixed sample was gently pipetted on top of this gradient and centrifuged at 3,000 RCF for 20 minutes at 4°C in a swinging bucket rotor with the brake off to avoid disrupting the gradient. Post-centrifugation, ∼150 µL of nuclei were collected at the 30%/40% interphase and transferred to a fresh 1.5 mL Eppendorf tube. Approximately 500 µL of wash buffer (1% BSA, 1 U/µL RNAse inhibitor, PBS) was added to each sample and incubated on ice for 5 minutes. Following the incubation, each sample was gently pipette-mixed to resuspend the nuclei and centrifuged at 500 RCF for 5 minutes at 4°C in a fixed-angle rotor. The supernatant was carefully removed from each sample, and nuclei were resuspended in ∼10-20 µL of diluted 10x Genomics Nuclei Buffer. To count the nuclei, 1 µL of the nuclei resuspension was mixed with 4 µL of PBS and 5 µL of 125 nM SYTOX (Thermo Fisher; #S7020) and incubated for 5 minutes. The number of GFP^+^ nuclei/µL was determined using the Countess II FL instrument (Thermo Fisher, #AMQAF1000). Each nuclei suspension was further diluted to 3,230-8,060 nuclei/µL, allowing for a targeted nuclei recovery of 10,000 nuclei per sample.

### 10x Genomics Single-nucleus RNA library preparation and sequencing

Single nucleus RNA-seq and snATAC-seq libraries were prepared using the Chromium Next GEM Single Cell Multiome ATAC + Gene Expression Kit (10x Genomics, #PN-1000283) following the Chromium Next GEM Single Cell Multiome ATAC + Gene Expression User Guide (CG000338 Rev F). Diluted nuclei suspensions from individual donors were loaded onto a 10x Genomics Chromium X instrument to generate Gel Beads-in-Emulsion. Quality control of cDNAs and libraries was performed using a Qubit Flex Fluorometer (Thermo Fisher, Q33327) and an Agilent 2100 Bioanalyzer (Agilent High Sensitivity DNA Kit, 5067-4626). Final library quantification prior to pooling was conducted with the KAPA Library Quantification Kit for Illumina Platforms (Kapa Biosystems, 07960255001). Single nucleus ATAC-seq libraries were pooled equimolarly and sequenced across four lanes of a NovaSeq X 25B flow cell, using the following run configurations: Read 1, 50 cycles; Read 2, 49 cycles; i5, 24 cycles; and i7 Index, 8 cycles each. Single nucleus RNA-seq libraries were separated by tissue type and pooled equimolarly. Single nucleus RNA-seq libraries from motor cortex and lumbar spinal cord were each sequenced across two lanes of a NovaSeq X 25B flow cell, using the following run configurations: Read 1, 28 cycles; Read 2, 90 cycles; i5 and i7 Index, 10 cycles each.

### 10x Genomics Multiome Data Processing

Single nucleus RNA-seq libraries were aligned using the GRCh38 reference genome (hg38; GENCODE v32; Cellranger human reference 2020-A), and a count matrix was generated using Cellranger-ARC v2.0.2. Cellranger QC metrics for LSC and MTC can be found in Supplemental Tables 2 and 3, respectively. LSC and MTC nuclei were included if they had between 500 and 25,000 gene expression transcripts (UMIs) and a mitochondrially-encoded transcript fraction less than 5% (LSC) and 20% (MTC). The LSC mitochondrially-encoded threshold was set lower because LSC samples had a large number of high MT-fraction ‘nuclei’ which were transcriptionally similar to motor neurons but lacked nuclear transcript markers (e.g. *MALAT1*) and are likely to be non-nuclei debris. We did not observe comparable debris in the MTC. A total of 139,103 LSC nuclei from 20 libraries and 266,234 MTC nuclei from 24 libraries passed this initial filtering.

To account for sequencing depth across nuclei, transcript counts for each nucleus were normalized to a total of 10,000. This was achieved by dividing all transcript counts for each nucleus by its size factor, to reach a total of 10,000 counts per nucleus. Normalized counts were log(x+1) transformed, and each gene was scaled by dividing by the interquartile range (RobustScaling). Genes with a variance greater than 1 after scaling were further scaled so that their variance was equal to 1, resulting in log standardized snRNA-seq counts.

Log standardized counts were projected by Principal Component Analysis (PCA) into 25 components. Nuclei were then embedded into a 25-neighbor k-Nearest Neighbors (k-NN) graph using these 25 components, and this graph was used for Uniform Manifold Approximation and Projection (UMAP) (Minimum distance of 0.5; initialized from spectral graph embedding).

### Cell type and subtype annotations

For LSC snRNA-seq data, log-normalized counts were used for PCA (25 components), k-NN graph construction (25 neighbors), UMAP, and Leiden clustering (resolution = 0.5). Each cluster was classified as neuronal or non-neuronal based on expression of canonical marker genes. Neuronal nuclei were then re-clustered (PCA: 25 components; k-NN: 15 neighbors; Leiden resolution = 0.15) and classified as excitatory, inhibitory, motor neuron, or doublets based on marker expression. Non-neuronal nuclei were similarly re-clustered (same PCA and k-NN parameters; Leiden resolution = 0.5) and annotated as doublets, astrocyte, endothelial, meninge, microglia, OPC, or oligodendrocyte. Finally, each non-neuronal cell type was further subclustered (PCA: 25 components; k-NN: 15 neighbors; Leiden resolution = 0.5 or 1.5) to define subtype-level populations or identify doublets. MTC snRNA-seq data was similarly processed and annotated. Gene markers can be found in Supplemental Table 6 and in Petrescu et al^28^.

### Integrating MTC snRNA-seq from Pineda et al. (2025)

MTC snRNA-seq data from Pineda et al.^30^, was standardized as above, and log-normalized counts were used for PCA (25 components), k-NN graph construction (25 neighbors), UMAP, and Leiden clustering (resolution = 0.5). Broad cell types were annotated as above, and each broad cell type was then separately reclustered and annotated for neuronal subtype or glial functional class based on expression of key marker transcripts. The annotated transcript count data from this work and Pineda et al was concatenated, and the concatenated count data was standardized by depth normalization and log transformation. No batch correction was performed. Standardized expression data was projected by PCA (25 components), embedded into a k-NN graph, and projected by UMAP for plotting.

### Cell proportions per sample

For each snRNA-seq library, the number of nuclei in each cell subtype was divided by the total number of nuclei for that cell type (e.g., 20 GM astrocytes and 80 WM astrocytes correspond to proportions of 0.2 and 0.8, respectively). Welch’s t-test was used to assess differences in proportions between groups. To determine the relationship between cell subtypes within each donor, the correlation between these ratios was calculated. Wald test was used to assess correlations and multiple comparisons were corrected using Benjamini-Hochberg.

### snRNA-seq Differential Expression Gene Analysis

Libraries with at least 20 nuclei for each cell type and at least 5 nuclei per cell subtype, and genes with average expression levels below 0.1 within each cell type or subtype were excluded. Nuclei were then aggregated at the donor level by summing counts for each nuclei. Differential expression gene analysis was performed using pyDESeq2 on the raw summed integer counts^51,52^. Only non-zero counts were used for standardizing. Mitochondrially encoded genes and *MALAT1* were excluded from size factor calculations. Statistical significance was defined as an adjusted *p*-value < 0.05 under a null hypothesis of log_2_FC ≤ ±0.263.

### snATAC-seq Differential Peak Accessibility Analysis

Peak counts from cellranger-arc were aggregated at the donor level by summing counts for each peak, for each cell type and subtype. Pseudobulked observations from fewer than 5 nuclei were removed, and the resulting count matrix was used for differential peak accessibility using pyDESeq2, standardizing with non-zero counts (‘poscounts’). Statistical significance was defined as an adjusted *p*-value < 0.05 under a null hypothesis of log_2_FC = 0.

### snATAC-seq Motif Enrichment Analysis

Sequences of differentially accessible peaks were extracted from the HG38 reference genome with bedtools, and these sequences were searched for enriched motifs with Simple Enrichment Analysis from the MEME Suite, against the JASPAR motif database (1661 TFs). Enriched motifs were defined as having a *q*-value for false discovery rate < 0.05.

### Tissue processing, RNA capture, and library preparation for spatial transcriptomics

Fresh frozen post-mortem motor cortex and lumbar spinal cord tissue blocks embedded in Tissue Plus OCT Compound (Fisher Healthcare, #4585) were stored at -80°C prior to processing and were placed in a cryostat at -16°C to equilibrate before sectioning. Ten micron tissue sections were placed onto pre-chilled Visium Spatial Gene Expression Slides (10x Genomics, #1000185). Four sections, each approximately 40 µm apart, per subject were collected across two Visium slides. When necessary, technical replicates from subjects with arrays that did not pass downstream QC metrics were repeated to maintain balanced motor cortex and spinal cord cohorts.

Spatially resolved gene expression data was generated following the 10x Genomics’ Visium Spatial Gene Expression Reagent User Guide (CG000239, Rev F and G). Briefly, tissue sections were methanol-fixed and stained with hematoxylin and eosin (H&E). Brightfield RGB histological images were obtained using an EC Plan-Neofluar 10x/0.3 M27 objective on a Zeiss Axio Observer Z1 equipped with a Zeiss Axiocam 506 mono camera. Motor cortex and spinal cord tissue sections were permeabilized for 12 minutes, allowing for the release and binding of poly-adenylated mRNA onto adjacent barcoded capture probes. The number of cDNA amplification cycles for each Visium array was determined by qPCR, as instructed by 10x Genomics. Amplified cDNA was quantified using the Agilent 2100 Bioanalyzer (Agilent High Sensitivity DNA Kit, 5067-4626).

Visium libraries were generated using Illumina-compatible PCR primers with Dual Index Kit TT Set A (10x Genomics, PN-1000215). Quality control of libraries was performed using a Qubit Flex Fluorometer (Thermo Fisher, Q33327) and an Agilent 2100 Bioanalyzer (Agilent High Sensitivity DNA Kit, 5067-4626). Final library quantification prior to pooling was conducted with the KAPA Library Quantification Kit for Illumina Platforms (Kapa Biosystems, 07960255001). Visium libraries from 48-52 arrays were each diluted to the same concentration, ranging from 5-10 nM, and pooled equimolarly for sequencing. Pooled Visium libraries were sequenced on either NovaSeq-X 10B or 25B, using the following run configurations: Read 1 and Read 2, 100 cycles each; i5 and i7 index reads, 10 cycles each. We processed a total of 137 LSC arrays from 35 donors and 183 MTC arrays from 50 donors.

### 10x Genomics Visium Data Processing and Annotation

Raw FASTQ files from sequenced Visium libraries were aligned to the same GRCh38 reference genome (hg38; GENCODE v32; Cellranger human reference 2020-A) with spaceranger v.3.0.0. Spaceranger metrics for LSC and MTC can be found in Supplemental Tables 4 and 5, respectively.

Using the 10x Genomics’ Loupe Browser, each spatial spot was manually annotated to one of eleven anatomical regions of the LSC based on hematoxylin and eosin (H&E) staining, cellular composition, and anatomical landmarks. The gray matter was distinguished by its butterfly-like shape and light pink hue, and contained: 1) the central canal, located at the center of the spinal cord and characterized by its small, densely packed nuclei; 2) the ventral horn, characterized by the presence of very large lower MNs; 3) the dorsal horn, consisted of a heterogeneous mixture of nuclei sizes; 4) the medial grey, the region between the dorsal and ventral horns and surrounding the central canal.

The white matter of the spinal cord was distinguished by its purple hue and contained: 5) the dorsal medial white region, situated between the dorsal horns and where the dorsal column-medial lemniscus pathway is found; 6) the medial lateral white, situated between the dorsal and ventral horns and adjacent to the medial grey, where the lateral corticospinal tract runs; 7) the ventral medial white, situated between the ventral horns; 8) the ventral lateral white, which contains the axons of lower motor neurons; and (9-11) the dorsal, lateral, and ventral edges, which surround the outer borders of the corresponding white matter regions. Any spot containing tissue artifacts, such as folds or tears, were excluded from annotations.

Spatial spots for MTC arrays were assigned one of six anatomical regions: 1) the white matter, characterized by its distinct purple hue and the presence of small, compact glial nuclei organized in a linear fashion; 2) layer 1, identified by its light pink hue and sparsely populated nuclei; 3) layer 2, characterized by a thin layer of densely packed small nuclei; 4) layer 3, marked by its smaller pyramidal neurons; 5) layer 5, marked by the presence of very large pyramidal neurons, the upper motor neurons; 6) layer 6, consisting of a heterogenous mixture of pyramidal neurons. Any spot containing tissue artifacts, such as folds or tears, were excluded from annotations.

To maintain the difference in count depth between anatomical regions, spots were standardized for count depth by dividing all transcript counts by a size factor, to reach the median count depth of its annotated anatomical region. Size factors were clipped at a minimum of 0.25 to prevent overinflating spots with low counts. Standardized spatial counts were then log(x+1) transformed and scaled by dividing by the interquartile range (RobustScaler). Any genes with a variance greater than 1 after scaling were further scaled so that their variance was equal to 1, resulting in log standardized spatial counts.

PCA was performed on the standardized spatial counts using a truncated singular value decomposition with 25 components. Spatial spots were embedded into a 25-neighbor k-NN graph constructed from these principal components and then projected into 2 dimensions with UMAP (Minimum distance of 0.5; initialized from spectral graph embedding).

### Splotch Modeling of Spatial Gene Expression across Sites of Symptom Onset

Raw sequencing data for LSC and MTC were separately filtered to remove mitochondrially encoded genes, lncRNAs, pseudogenes, and genes expressed in less than 1% of spots. Spatial spots with less than 100 total UMIs for the remaining genes were excluded from Splotch modeling. Spots without an assigned annotation were discarded, as were any spots without at least one immediate neighbor in the Visium grid to prevent discontinuities in the spatial autocorrelation component of the Splotch model.

Splotch modeling and analysis was performed as previously described^26,28,53^. In brief, Splotch utilizes a zero-inflated Poisson likelihood function (where the zero-inflation accounts for technical dropouts) to model the expression (λ) of each gene in each spot. This expression level is then modeled using a generalized linear model (GLM) with the following three components: the characteristic expression of the spot’s annotated anatomical region (β), autocorrelation with the spot’s spatial neighbors (ψ) and spot-level variation (ε). Furthermore, the characteristic expression rate (β) is hierarchically formulated to account for the experimental design, with donor-level effects (level 3) conditioned on ALS presentation (level 2) and further on ALS diagnosis (level 1), enabling the investigation of the effect of sample covariates on differential gene expression. By performing posterior inference on the model using the observed spatial expression data, we identified disease-level trends in spatial gene expression (β) that best explained our observations.

### Parameter Inference

Splotch has been implemented in the probabilistic programming language Stan, and is available at https://github.com/adaly/cSplotch. For all analyses, Bayesian inference was performed over the parameters using Stan’s adaptive Hamiltonian Monte-Carlo (HMC) sampler with default parameters. Four independent chains were sampled, each with 175 warm-up iterations and 175 sampling iterations (700 total), and convergence was monitored using the R-hat statistic.

### Splotch differential expression analysis

To assess differential expression between two sample conditions “i” and “j” – which without loss of generality can span different tissue types, diagnostic groups (ALS or control), sites of onset, or donor IDs – we examine posterior distributions over Splotch parameters β_*i*_ and β_j_. From these posterior estimates, we can calculate two quantities: (1) the mean log2-fold change in expression, and (2) the Bayes factor quantifying the strength with which we reject the null hypothesis of identical expression (*P*(β_*i*_-β_j_=0)). The Bayes factor (BF) was calculated using the Savage-Dickey density ratio of the likelihood of the null hypothesis under the prior vs. the posterior model^54^. Larger values indicate that the data support rejection of the null hypothesis, with BF>5 indicating substantial support by convention.

### Identifying Spatial Gene Modules

Spatial gene modules were identified as previously described^28^. Briefly, spatial gene modules were generated using the spot-level gene expression measurements (λ) from Splotch, which produced a matrix of 11,990 genes x 579,373 spots in LSC and 13,727 genes x 649,436 spots in MTC. The spot-level gene measurements were capped at the 99^th^ percentile to minimize the influence of outliers and scaled by dividing by the gene interquartile range. Genes with high variability (variance > 1) were scaled again to have their variance equal to 1. Pearson correlation coefficients were computed for every gene pair across all spots, forming 11,990 x 11,990 and 13,727 x 13,727 correlation matrices for LSC and MTC, respectively. The correlation distance (1 – Pearson correlation coefficient) between each gene-gene pair was computed and used as the distance metric for a gene-gene k-NN graph embedding (k=10). Genes were projected into 2 dimensions using UMAP and grouped into modules using Leiden clustering (resolution = 2). The scself package (single-cell self-supervised; v0.4.8) in Python was used to help identify spatial gene modules. Genes for each LSC and MTC spatial gene module are in Supplemental Tables 8 and 12, respectively.

### Module Scores

Each gene’s expression (the standardized transcript count for spatial and snRNA-seq data) was scaled to a min-max range of 0-1, where 0 and 1 were set to the 1^st^ and 99^th^ percentile, respectively. Any value below the 1^st^ percentile or above the 99^th^ percentile was set to 0 or 1, respectively. For spatial data, raw integer counts were aggregated by averaging across each annotated region for each array and standardized for count depth by dividing aggregated transcript counts by a size factor, to reach the median count depth of all arrays. Size factors were clipped at a minimum of 0.25 to prevent overinflating spots with low counts. Standardized spatial counts were then log(x+1) transformed and scaled by dividing by the interquartile range (RobustScaler). Any genes with a variance greater than 1 after scaling were further scaled so that their variance was equal to 1, resulting in log standardized spatial counts. Module scores were then calculated for each anatomical region per array from the aggregated count data for statistical testing. For snRNA-seq, a module score was calculated for each nuclei from log standardized counts.

For spatial data, the module score for each anatomical region per array was used for statistical testing. Welch’s t-test was used for statistical tests between module scores in different groups, controlling for false discovery with Benjamini-Yekutieli (FDR-BY). For plotting, the module scores for each anatomical region were averaged across all the arrays per donor and scaled to a 0-1 range.

For snRNA-seq data, module scores were aggregated for each donor tissue (LSC or MTC) sample by averaging the score for each nuclei assigned a cell type. Welch’s t-test was used for statistical tests between module scores in different groups, controlling for false discovery with Benjamini-Hochberg (FDR-BH). The aggregated module scores were scaled to a 0-1 for plotting.

### Assigning Genes to Cell Type Submodules

For each spatial gene module, the snRNA-seq correlation distance (1 – Pearson correlation coefficient) between each gene-gene pair was used as the distance metric for a gene-gene k-NN graph embedding (k=10). Genes were projected into 2 dimensions using UMAP and grouped into submodules with Leiden clustering (resolution = 0.75). Submodule scores were calculated using the same method as module scores for snRNA-seq nuclei and spatial regions. Genes for each LSC and MTC spatial gene submodule are in Supplemental Tables 8 and 12, respectively.

### Gene Ontology Enrichment Analysis

Gene ontology enrichment was performed as a multiple hypothesis test corrected Fisher’s exact test using g:Profiler^55^. Gene lists tested consisted of either differentially expressed genes against a background of genes with non-zero expression, or module genes against a background of all genes that were modeled for spatial gene modules. Gene ontology enrichment for LSC and MTC spatial gene modules are in Supplemental Tables 9 and 13, respectively. Gene ontology enrichment for LSC and MTC spatial submodules are in Supplemental Tables 10 and 14, respectively.

### Ventral Horn Cell Type Masks and Motor Neuron Quantification

Using the H&E stained spinal cord images (*n* = 137) obtained during the Visium workflow, motor neuron and non-motor neuron masks were generated for cells in the ventral horn and manually corrected using napari^56^. Any array in which the ventral horn contained tissue artifacts, such as folds or tears, were excluded from the masks. In total, masks were generated for 122 LSC Visium arrays. For each array, the number of motor neuron masks was divided by the total number of spatial spots annotated as ventral horn and multiplied by 100, resulting in the number of motor neurons per 100 ventral horn spots.

### Optical Density of ROIs

The H&E stained spinal cord images (*n* = 137) obtained during the Visium workflow were manually segmented into annotated anatomical regions. To model gene expression and compare staining across regions of interest (ROIs) from the same image, an optical density (OD) was calculated for each Visium spot on every array. OD is defined as follows:

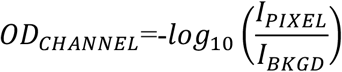

*OD_CHANNEL_* is the optical density in the image red, green, or blue channels, I*_PIXEL_* is the Visium spot pixel intensity and I*_BKGD_* is the estimated background intensity. For images without background, background intensity (I*_BKGD_*) was approximated by finding the brightest pixel (highest RGB value) in each image. Scalar OD for each Visium spot was computed as the Euclidean norm of the three OD color channels:

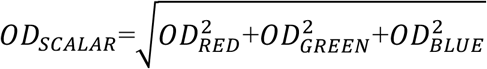

OD_SCALAR_ was used per-spot for modeling or averaged across all pixels within an annotated anatomical region to get the reported OD for the ROI on the array.

### Inferring Corticospinal Tract Loss

Optical density was quantified for each Visium spot in the dorsal medial white (DMW), medial lateral white (MLW), ventral lateral white (VLW), and ventral medial white (VMW) regions from each H&E image. For each image, spot-level OD_SCALAR_ measurements were min-max scaled 0-1. Scaled OD values in DMW were subtracted from the OD values in the MLW, VLW, and VMW (ΔOD_ROI_ = OD_ROI_ – OD_DMW_) and the resulting differences were averaged per region for each donor to adjust for regional changes in staining intensity.

### Using ΔOD as Predictor for Differential Expression Gene Analysis

Differential expression gene analysis was performed using pyDESeq2^51,52^ on the raw integer counts from MTC spatial data. The dataset included 18 donors (4 controls; 14 ALS) with paired MTC and LSC spatial data. MTC spatial counts were aggregated by summing counts for all spots per region to the donor. ΔOD measurements for medial lateral white, ventral lateral white, and ventral medial white matter regions were averaged per donor and used as the predictor in pyDESeq2. Age and sex were also included as covariates in the model. Only non-zero counts were used for standardizing. Mitochondrially encoded genes and *MALAT1* were excluded from size factor calculations. Statistical significance was defined as an adjusted *p*-value < 0.05 under a null hypothesis that log_2_ fold change is zero. These results can be found in Supplemental Table 15.

### Finding overlap in LSC and MTC Spatial Gene Modules

LSC and MTC spatial datasets were restricted to genes present in both datasets and assigned to spatial gene modules. Pairwise Jaccard index values were calculated to quantify the similarity between each LSC and MTC spatial module pair, defined as:

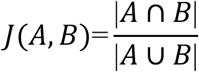

For each LSC module, the MTC module with the highest Jaccard similarity (threshold > 0.1) was selected. Genes shared between LSC and MTC spatial gene modules can be found in Supplemental Table 16.

### General Procedures

Unless specified, data were analyzed using the scanpy^57^ framework in Python, and plots were generated with Matplotlib^58^ and seaborn^59^. Figures were assembled in Adobe Illustrator.

## ACKNOWLEDGEMENTS

We are grateful to the donors whose contributions made this research possible. We thank the Target ALS and ALS Consortium postmortem core for their tissue procurement. This study was supported by awards from the National Institutes of Health (R01NS118183, R01NS116350, R01NS127186, and R01NS118570) and Target ALS (FD-2023-GEN-S1). ChatGPT (OpenAI) was used to assist with language editing of the manuscript text. The New York Genome Center and the Scientific Computing Core at the Flatiron Institute, a division of the Simons Foundation, provided computational resources for this study. We thank Maria Hauge Pedersen for their efforts in project coordination.

## AUTHOR CONTRIBUTIONS

Conceptualization: NB, HP, CS; Cohort design: NB, KK, JP; Resources: HP, CS; Data generation (ST and snRNA-seq experiments): NB, OC, OK, KK, JP, ML, SK; Software: AD, JE, CAJ; Data processing: NB, CAJ, AD, JE, BG; Data analysis: NB, AD, JE, CAJ; Data visualization: NB, AD, CAJ; Writing: NB, CAJ, HP; Editing: NB, BF, AD, OK, MX, JE, CAJ, HP; Supervision: CAJ, HP; Project administration: HP; Funding acquisition: HP.

## DATA AVAILABILITY

Paired snRNA-seq and snATAC-seq and spatial transcriptomic sequencing data have been deposited in NCBI GEO (GSE330130 and GSE330224, respectively).

