## Supplemental Figures for "A multi-omics characterization reveals distinct molecular signatures in the human motor cortex and lumbar spinal cord in ALS"

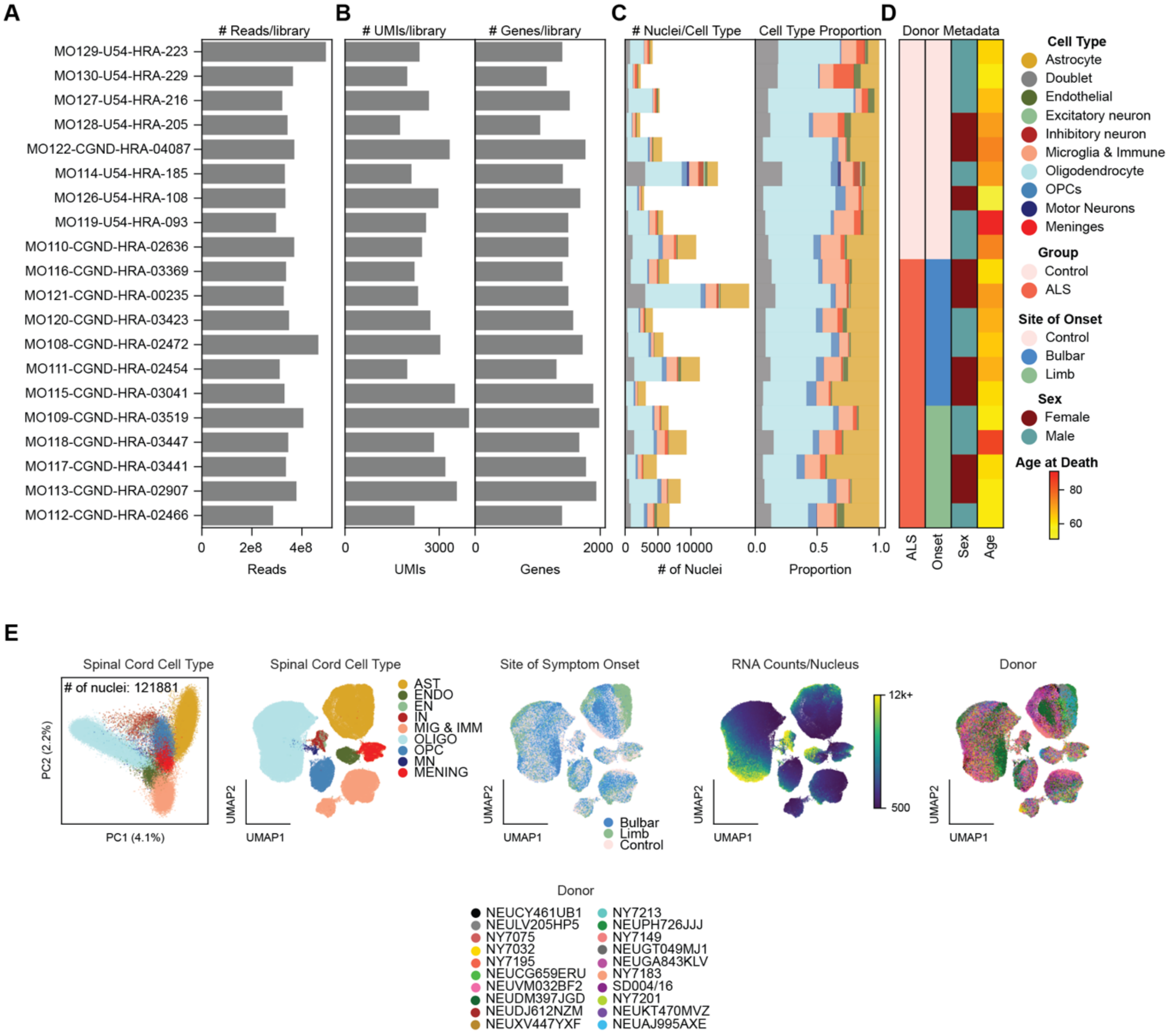
A
B
C
D
### Reads/library
### UMIs/library
### Genes/library
### Nuclei/Cell Type
Cell Type Proportion
Donor Metadata
MO129-U54-HRA-223
MO130-U54-HRA-229
MO127-U54-HRA-216
MO128-U54-HRA-205
MO122-CGND-HRA-04087
MO114-U54-HRA-185
MO126-U54-HRA-108
MO119-U54-HRA-093
MO110-CGND-HRA-02636
MO116-CGND-HRA-03369
MO121-CGND-HRA-00235
MO120-CGND-HRA-03423
MO108-CGND-HRA-02472
MO111-CGND-HRA-02454
MO115-CGND-HRA-03041
MO109-CGND-HRA-03519
MO118-CGND-HRA-03447
MO117-CGND-HRA-03441
MO113-CGND-HRA-02907
MO112-CGND-HRA-02466
0 2e8 4e8
Reads
0 3000
UMIs
0 2000
Genes
0 5000 10000
### of Nuclei
0.0 0.5 1.0
Proportion
ALS
Onset
Sex
Age
Cell Type
Astrocyte
Doublet
Endothelial
Excitatory neuron
Inhibitory neuron
Microglia & Immune
Oligodendrocyte
OPCs
Motor Neurons
Meninges
Group
Control
ALS
Site of Onset
Control
Bulbar
Limb
Sex
Female
Male
Age at Death
80
60
E
Spinal Cord Cell Type
### of nuclei: 121881
PC2 (2.2%)
PC1 (4.1%)
Spinal Cord Cell Type
AST
ENDO
EN
IN
MIG & IMM
OLIGO
OPC
MN
MENING
UMAP2
UMAP1
Site of Symptom Onset
Bulbar
Limb
Control
RNA Counts/Nucleus
12k+
500
Donor
Donor
NEUCY461UB1
NEULV205HP5
NY7075
NY7032
NY7195
NEUCG659ERU
NEUVM032BF2
NEUDM397JGD
NEUDJ612NZM
NEUXV447YXF
NY7213
NEUPH726JJJ
NY7149
NEUGT049MJ1
NEUGA843KLV
NY7183
SD004/16
NY7201
NEUKT470MVZ
NEUAJ995AXE

**Supplemental Figure 1: LSC snRNA-seq Quality Control.**

**A)** Number of paired-end reads per library.

**B)** Median number of UMI counts and genes per nucleus before doublet removal.

**C)** Cell type annotations by transcriptomic-inference, including doublets. Total number of nuclei and distribution of cell types per library (left). Proportion of cell types from total nuclei per library (right).

**D)** Cohort metadata.

**E)** PCA and UMAP plots of snRNA-seq data. Each dot represents an individual nucleus and is colored by cell type, site of symptom onset, UMI counts per nucleus, or donor ID.

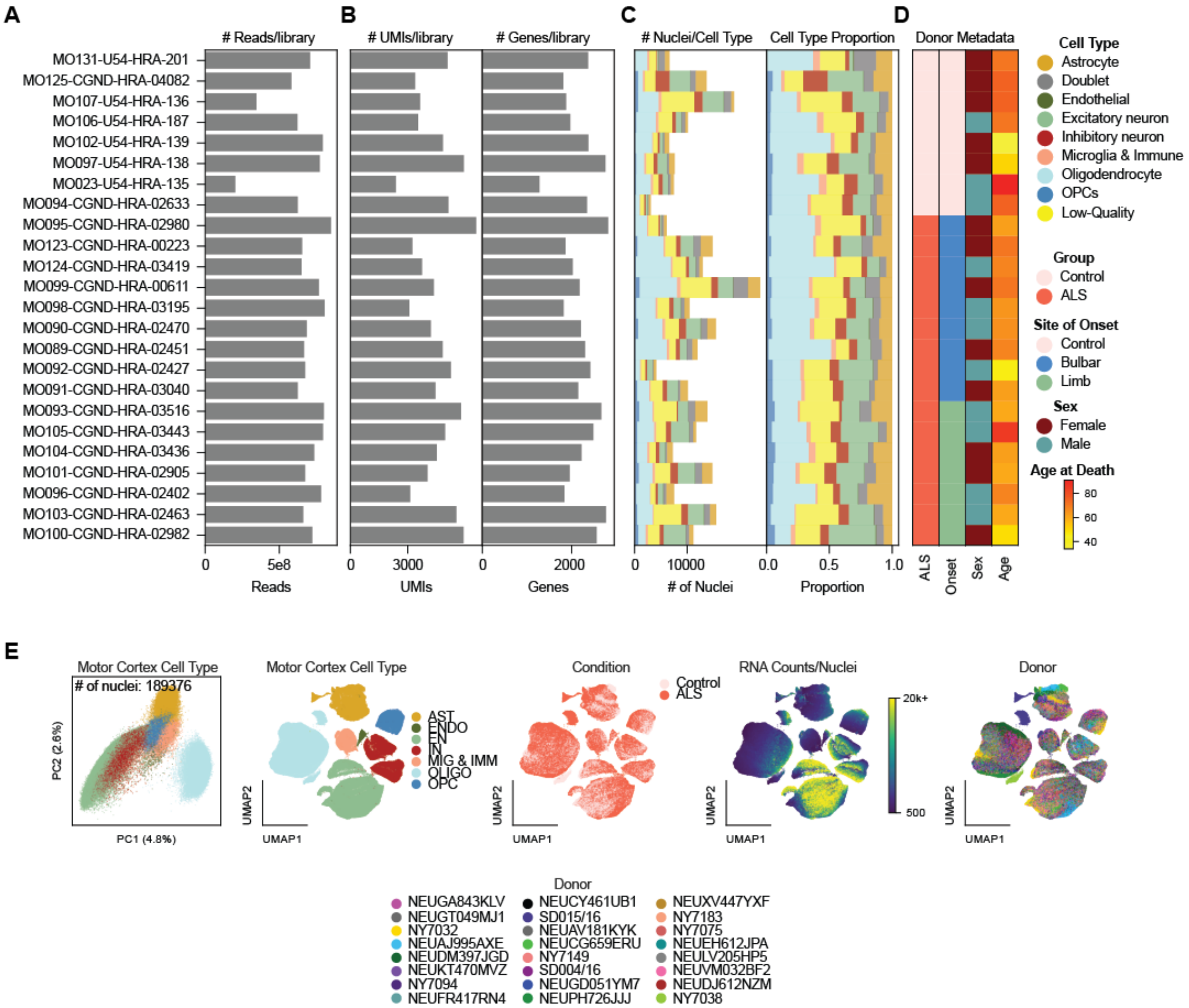
A
### Reads/library
B
### UMIs/library
### Genes/library
C
### Nuclei/Cell Type
Cell Type Proportion
D
Donor Metadata
MO131-U54-HRA-201
MO125-CGND-HRA-04082
MO107-U54-HRA-136
MO106-U54-HRA-187
MO102-U54-HRA-139
MO097-U54-HRA-138
MO023-U54-HRA-135
MO094-CGND-HRA-02633
MO095-CGND-HRA-02980
MO123-CGND-HRA-00223
MO124-CGND-HRA-03419
MO099-CGND-HRA-00611
MO098-CGND-HRA-03195
MO090-CGND-HRA-02470
MO089-CGND-HRA-02451
MO092-CGND-HRA-02427
MO091-CGND-HRA-03040
MO093-CGND-HRA-03516
MO105-CGND-HRA-03443
MO104-CGND-HRA-03436
MO101-CGND-HRA-02905
MO096-CGND-HRA-02402
MO103-CGND-HRA-02463
MO100-CGND-HRA-02982
0
5e8
Reads
0
3000
UMIs
0
2000
Genes
0
10000
### of Nuclei
0.0
0.5
1.0
Proportion
ALS
Onset
Sex
Age
Cell Type
Astrocyte
Doublet
Endothelial
Excitatory neuron
Inhibitory neuron
Microglia & Immune
Oligodendrocyte
OPCs
Low-Quality
Group
Control
ALS
Site of Onset
Control
Bulbar
Limb
Sex
Female
Male
Age at Death
80
60
40
E
Motor Cortex Cell Type
### of nuclei: 189376
PC2 (2.6%)
PC1 (4.8%)
Motor Cortex Cell Type
AST
ENDO
EN
IN
MIG & IMM
OLIGO
OPC
UMAP2
UMAP1
Condition
Control
ALS
UMAP2
UMAP1
RNA Counts/Nuclei
20k+
500
UMAP2
UMAP1
Donor
UMAP2
UMAP1
Donor
NEUGA843KLV
NEUGT049MJ1
NY7032
NEUAJ995AXE
NEUDM397JGD
NEUKT470MVZ
NY7094
NEUFR417RN4
NEUCY461UB1
SD015/16
NEUAV181KYK
NEUCG659ERU
NY7149
SD004/16
NEUGD051YM7
NEUPH726JJJ
NEUXV447YXF
NY7183
NY7075
NEUEH612JPA
NEULV205HP5
NEUVM032BF2
NEUDJ612NZM
NY7038

**Supplemental Figure 2: MTC snRNA-seq Quality Control.**

**A)** Number of paired-end reads per library.

**B)** Median number of UMI counts and genes per nucleus before doublet removal.

**C)** Cell type annotations by transcriptomic-inference, including doublets. Total number of nuclei and distribution of cell types per library (left). Proportion of cell types from total nuclei per library (right).

**D)** Cohort metadata.

**E)** PCA and UMAP plots of snRNA-seq data. Each dot represents an individual nucleus and is colored by cell type, condition, UMI counts per nucleus, or donor ID.

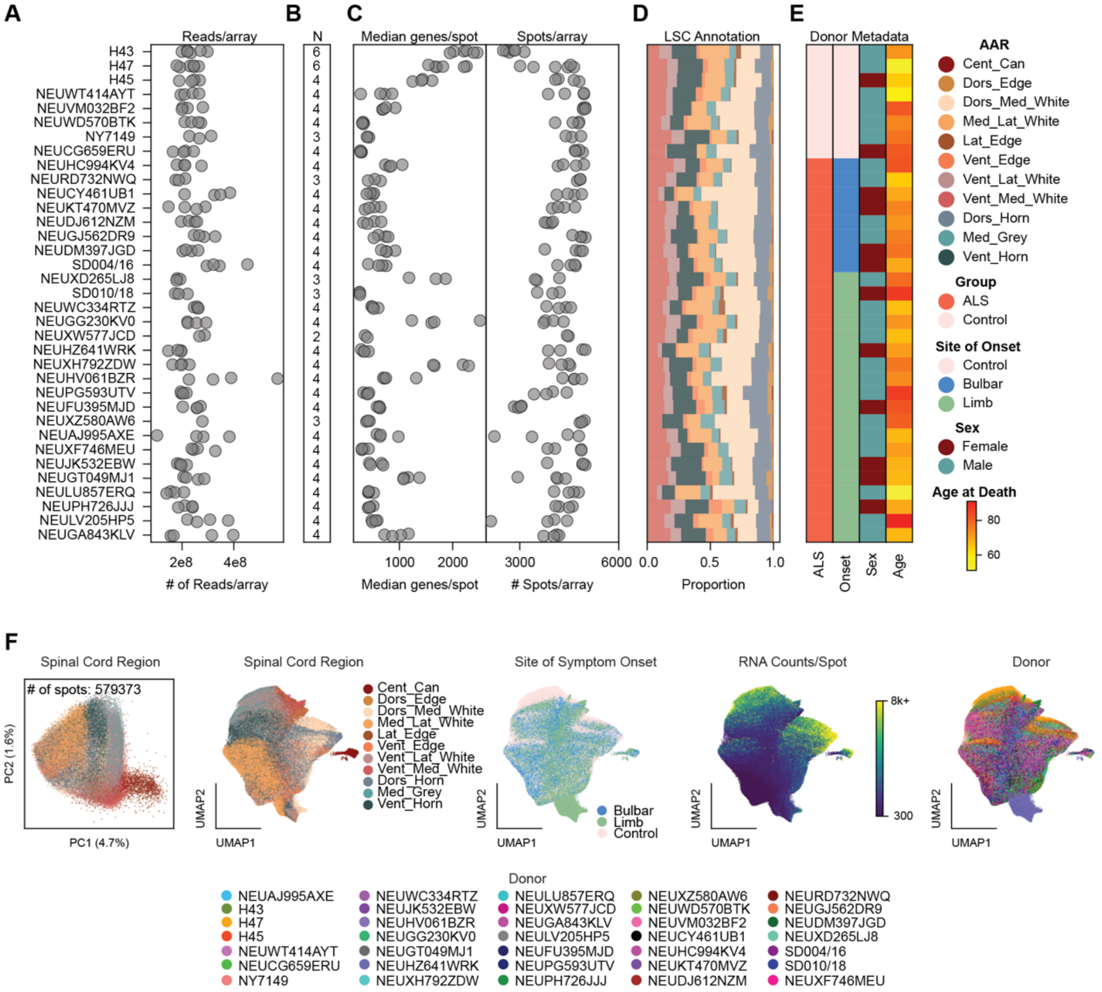

A
Reads/array
H43
H47
H45
NEUWT414AYT
NEUVM032BF2
NEUWD570BTK
NY7149
NEUCG659ERU
NEUHC994KV4
NEURD732NWQ
NEUCY461UB1
NEUKT470MVZ
NEUDJ612NZM
NEUGJ562DR9
NEUDM397JGD
SD004/16
NEUXD265LJ8
SD010/18
NEUWC334RTZ
NEUGG230KV0
NEUXW577JCD
NEUHZ641WRK
NEUXH792ZDW
NEUHV061BZR
NEUPG593UTV
NEUFU395MJD
NEUXZ580AW6
NEUAJ995AXE
NEUXF746MEU
NEUJK532EBW
NEUGT049MJ1
NEULU857ERQ
NEUPH726JJJ
NEULV205HP5
NEUGA843KLV
2e8
4e8
### of Reads/array
B
N
6 6 4 4 4 4 3 4 4 3 4 4 4 4 4 4 3 3 4 4 2 4 4 4 4 4 3 4 4 4 4 4 4 4 4
C
Median genes/spot
Spots/array
1000
2000
3000
6000
Median genes/spot
### Spots/array
D
LSC Annotation
0.0
0.5
1.0
Proportion
E
Donor Metadata
ALS
Onset
Sex
Age
AAR
Cent_Can
Dors_Edge
Dors_Med_White
Med_Lat_White
Lat_Edge
Vent_Edge
Vent_Lat_White
Vent_Med_White
Dors_Horn
Med_Grey
Vent_Horn
Group
ALS
Control
Site of Onset
Control
Bulbar
Limb
Sex
Female
Male
Age at Death
80
60
F
Spiral Cord Region
### of spots: 579373
PC2 (1.6%)
PC1 (4.7%)
Spinal Cord Region
Cent_Can
Dors_Edge
Dors_Med_White
Med_Lat_White
Lat_Edge
Vent_Edge
Vent_Lat_White
Vent_Med_White
Dors_Horn
Med_Grey
Vent_Horn
UMAP2
UMAP1
Site of Symptom Onset
Bulbar
Limb
Control
RNA Counts/Spot
8k+
300
Donor
Donor
NEUAJ995AXE
H43
H47
H45
NEUWT414AYT
NEUCG659ERU
NY7149
NEUWC334RTZ
NEUJK532EBW
NEUHV061BZR
NEUGG230KV0
NEUGT049MJ1
NEUHZ641WRK
NEUXH792ZDW
NEULU857ERQ
NEUXW577JCD
NEUGA843KLV
NEULV205HP5
NEUFU395MJD
NEUPG593UTV
NEUPH726JJJ
NEUXZ580AW6
NEUWD570BTK
NEUVM032BF2
NEUCY461UB1
NEUHC994KV4
NEUKT470MVZ
NEUDJ612NZM
NEURD732NWQ
NEUGJ562DR9
NEUDM397JGD
NEUXD265LJ8
SD004/16
SD010/18
NEUXF746MEU

**Supplemental Figure 3: LSC Spatial Transcriptomics Quality Control.**

**A)** Number of paired-end reads per Visium array.

**B)** Number of Visium arrays per donor.

**C)** Median number of genes per donor (left) and number of spots per Visium array (right)**.**

**D)** The average proportion of spots annotated per LSC anatomical region.

**E)** Cohort metadata.

**F)** PCA and UMAP plots of spatial data. Each dot represents an individual spatial spot colored by LSC annotated anatomical region (AAR), site of symptom onset, UMI counts per spot, or donor ID.

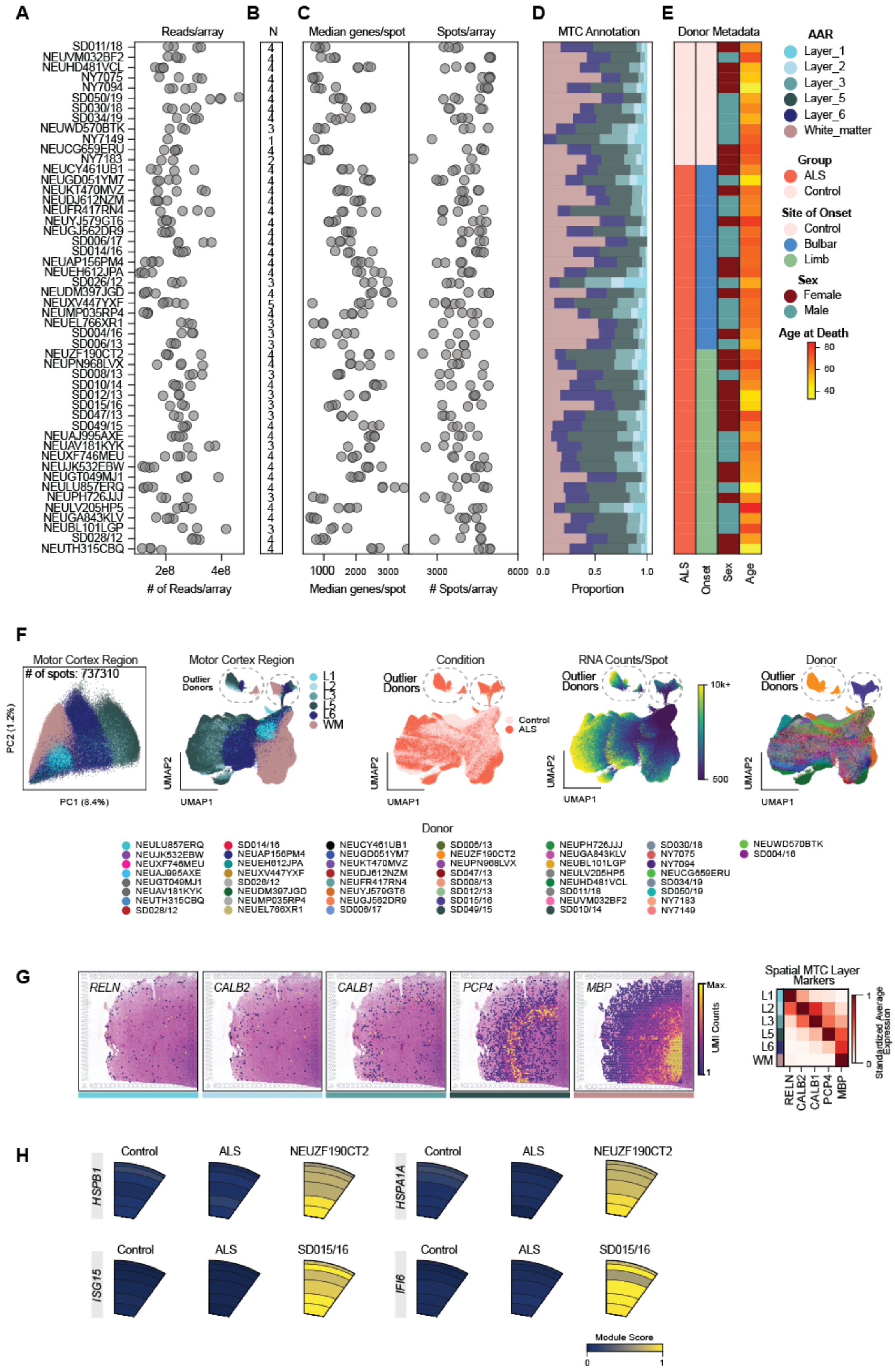
A
Reads/array
### of Reads/array
2e8
4e8
B
N
C
Median genes/spot
Spots/array
Median genes/spot
### Spots/array
1000
2000
3000
3000
6000
D
MTC Annotation
Proportion
0.0
0.5
1.0
E
Donor Metadata
ALS
Onset
Sex
Age
AAR
Layer_1
Layer_2
Layer_3
Layer_5
Layer_6
White_matter
Group
ALS
Control
Site of Onset
Control
Bulbar
Limb
Sex
Female
Male
Age at Death
80
60
40
SD011/18
NEUVM032BF2
NEUHD481VCL
NY7075
NY7094
SD050/19
SD030/18
SD034/19
NEUWD570BTK
NY7149
NEUCG659ERU
NY7183
NEUCY461UB1
NEUGD051YM7
NEUKT470MVZ
NEUDJ612NZM
NEUFR417RN4
NEUYJ579GT6
NEUGJ562DR9
SD006/17
SD014/16
NEUAP156PM4
NEUEH612JPA
SD026/12
NEUDM397JGD
NEUXV447YXF
NEUMP035RP4
NEUEL766XR1
SD004/16
SD006/13
NEUZF190CT2
NEUPN968LVX
SD008/13
SD010/14
SD012/13
SD015/16
SD047/13
SD049/15
NEUAJ995AXE
NEUAV181KYK
NEUXF746MEU
NEUJK532EBW
NEUGT049MJ1
NEULU857ERQ
NEUPH726JJJ
NEULV205HP5
NEUGA843KLV
NEUBL101LGP
SD028/12
NEUTH315CBQ
F
Motor Cortex Region
### of spots: 737310
PC2 (1.2%)
PC1 (8.4%)
Motor Cortex Region
Outlier Donors
L1
L2
L3
L5
L6
WM
UMAP2
UMAP1
Condition
Outlier Donors
Control
ALS
RNA Counts/Spot
Outlier Donors
10k+
500
Donor
Outlier Donors
Donor
NEULU857ERQ
NEUJK532EBW
NEUXF746MEU
NEUAJ995AXE
NEUGT049MJ1
NEUAV181KYK
NEUTH315CBQ
SD028/12
SD014/16
NEUAP156PM4
NEUEH612JPA
SD026/12
NEUXV447YXF
NEUDM397JGD
NEUMP035RP4
NEUEL766XR1
NEUCY461UB1
NEUGD051YM7
NEUKT470MVZ
NEUDJ612NZM
NEUFR417RN4
NEUYJ579GT6
NEUGJ562DR9
SD006/17
SD006/13
NEUZF190CT2
NEUPN968LVX
SD047/13
SD008/13
SD012/13
SD015/16
SD049/15
NEUPH726JJJ
NEUGA843KLV
NEUBL101LGP
NEULV205HP5
NEUHD481VCL
SD011/18
NEUVM032BF2
SD010/14
SD030/18
NY7075
NY7094
NEUCG659ERU
SD034/19
SD050/19
NY7183
NY7149
NEUWD570BTK
SD004/16
G
RELN
CALB2
CALB1
PCP4
MBP
Max.
UMI Counts
1
Spatial MTC Layer Markers
L1
L2
L3
L5
L6
WM
RELN
CALB2
CALB1
PCP4
MBP
Standardized Average Expression
1
0
H
HSPB1
Control
ALS
NEUZF190CT2
HSPA1A
Control
ALS
NEUZF190CT2
ISG15
Control
ALS
SD015/16
IFI6
Control
ALS
SD015/16
Module Score
0
1

**Supplemental Figure 4: MTC Spatial Transcriptomics Quality Control.**

**A)** Number of paired-end reads per Visium array.

**B)** Number of Visium arrays per donor.

**C)** Median number of genes per donor (left) and number of spots per Visium array (right)**.**

**D)** The average proportion of spots annotated per LSC anatomical region.

**E)** Cohort metadata.

**F)** PCA and UMAP plots of spatial data. Each dot represents an individual spatial spot colored by MTC annotated anatomical region (AAR), condition, UMI counts per spot, or donor ID.

**G)** i) Representative spot plots showing *RELN*, *CALB2/1*, *PCP4*, and *MBP* transcript expression overlaid on corresponding H&E-stained tissue section. Maximum expression is normalized per array for each transcript. ii) The average expression of each marker gene in each MTC region across all Visium arrays.

**H)** Expression of heat shock-related genes and immune response genes in two outlier donors.

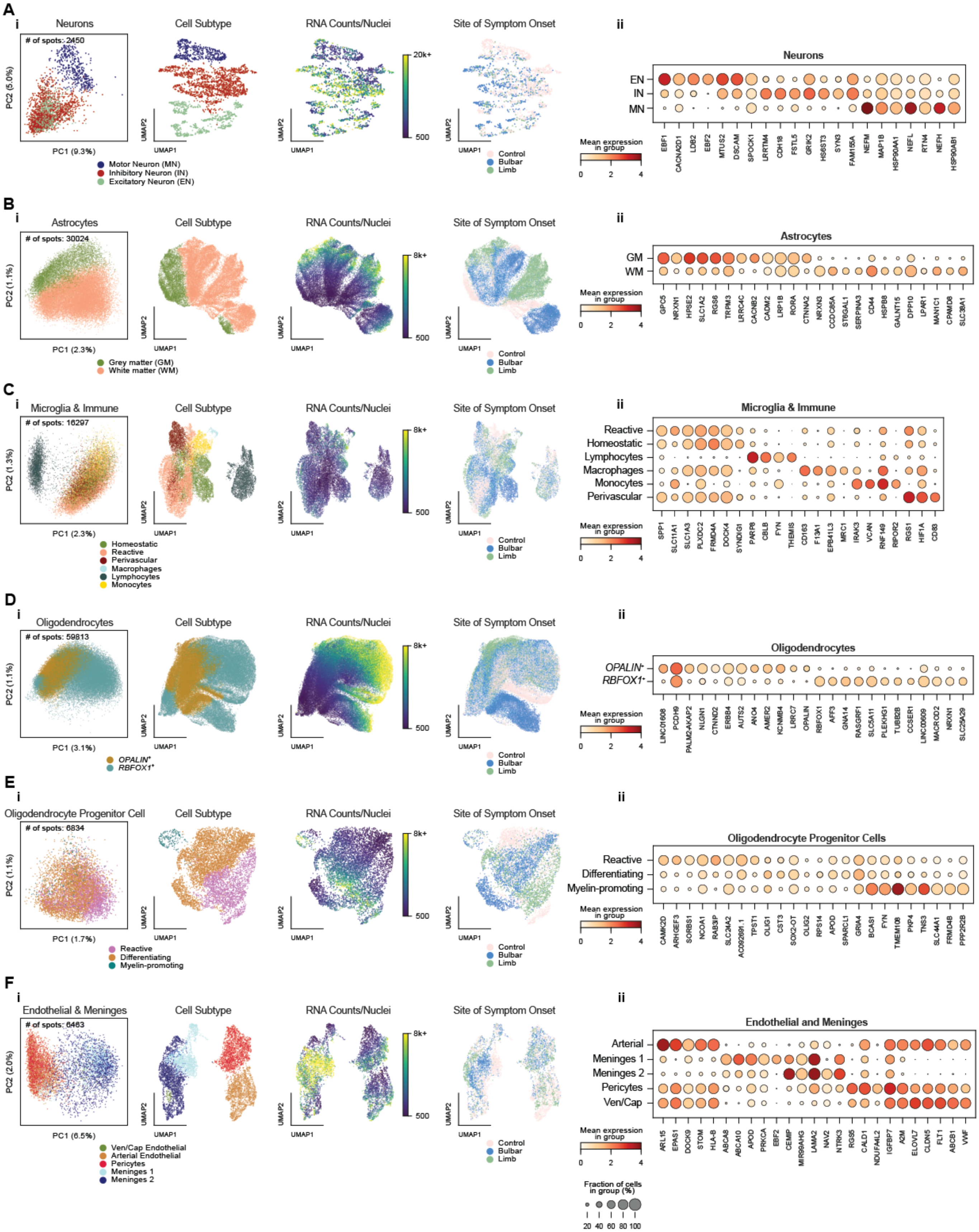

A
i
Neurons
### of spots: 2450
PC2 (5.0%)
PC1 (9.3%)
Motor Neuron (MN)
Inhibitory Neuron (IN)
Excitatory Neuron (EN)
Cell Subtype
RNA Counts/Nuclei
20k+
500
Site of Symptom Onset
Control
Bulbar
Limb
UMAP1
UMAP2
ii
Neurons
EN
IN
MN
Mean expression in group
0 2 4
EBF1 CACNA2D1 LDB2 EBF2 MTUS2 DSCAM SPOCK1 LRRTM4 CDH18 FSTL5 GRIK2 HS6ST3 SYN3 FAM155A NEFM MAP1B HSP90AA1 NEFL RTN4 NEFH HSP90AB1
B
i
Astrocytes
### of spots: 30024
PC2 (1.1%)
PC1 (2.3%)
Grey matter (GM)
White matter (WM)
Cell Subtype
RNA Counts/Nuclei
8k+
500
Site of Symptom Onset
Control
Bulbar
Limb
ii
Astrocytes
GM
WM
Mean expression in group
0 2 4
GPC5 NRXN1 HPSE2 SLC1A2 RGS6 TRPM3 LRRC4C CACNB2 CADM2 LRP1B RORA CTNNA2 NRXN3 CCDC85A ST6GAL1 SERPINA3 CD44 HSPB8 GALNT15 DPP10 LPAR1 MAN1C1 CPAMD8 SLC38A1
C
i
Microglia & Immune
### of spots: 16297
PC2 (1.3%)
PC1 (2.3%)
Homeostatic
Reactive
Perivascular
Macrophages
Lymphocytes
Monocytes
Cell Subtype
RNA Counts/Nuclei
8k+
500
Site of Symptom Onset
Control
Bulbar
Limb
ii
Microglia & Immune
Reactive
Homeostatic
Lymphocytes
Macrophages
Monocytes
Perivascular
Mean expression in group
0 2 4
SPP1 SLC11A1 SLC1A3 PLXDC2 FRMD4A DOCK4 SYNDIG1 PARP8 CBLB FYN THEMIS CD163 F13A1 EPB41L3 MRC1 IRAK3 VCAN RNF149 RBPOR2 RGS1 HIF1A CD83
D
i
Oligodendrocytes
### of spots: 59813
PC2 (1.1%)
PC1 (3.1%)
OPALIN+
RBFOX1+
Cell Subtype
RNA Counts/Nuclei
8k+
500
Site of Symptom Onset
Control
Bulbar
Limb
ii
Oligodendrocytes
OPALIN+
RBFOX1+
Mean expression in group
0 2 4
LINC01608 PCDH9 PALM2-AKAP2 NLGN1 CTNND2 ERBB4 AUTS2 ANO4 AMER2 KCNMB4 LRRC7 OPALIN RBFOX1 AFF3 GNA14 RASGRF1 SLC5A11 PLEKHG1 TUBB2B CCSER1 LINC00609 MACROD2 NRXN1 SLC25A29
E
i
Oligodendrocyte Progenitor Cell
### of spots: 6834
PC2 (1.1%)
PC1 (1.7%)
Reactive
Differentiating
Myelin-promoting
Cell Subtype
RNA Counts/Nuclei
8k+
500
Site of Symptom Onset
Control
Bulbar
Limb
ii
Oligodendrocyte Progenitor Cells
Reactive
Differentiating
Myelin-promoting
Mean expression in group
0 2 4
CAMK2D ARHGEF3 SORBS1 NCOA1 RAB3IP SLC24A2 AC092691.1 TPST1 OLIG1 CST3 SOX2-OT OLIG2 RPS14 APOD SPARCL1 GRIA4 BCAS1 FYN TMEM108 PKP4 TNS3 SLC44A1 FRMD4B PPP2R2B
F
i
Endothelial & Meninges
### of spots: 6463
PC2 (2.0%)
PC1 (6.5%)
Ven/Cap Endothelial
Arterial Endothelial
Pericytes
Meninges 1
Meninges 2
Cell Subtype
RNA Counts/Nuclei
8k+
500
Site of Symptom Onset
Control
Bulbar
Limb
ii
Endothelial and Meninges
Arterial
Meninges 1
Meninges 2
Pericytes
Ven/Cap
Mean expression in group
0 2 4
ARL15 EPAS1 DOCK9 STOM HLA-B ABCA8 ABCA10 APOD PRKCA EBF2 CEMIP MIR99AHG LAMA2 NAV2 NTRK3 RGS5 CALD1 NDUFA4L2 IGFBP7 A2M ELOVL7 CLDN5 FLT1 ABCB1 VWF
Fraction of cells in group (%)
20 40 60 80 100

**Supplemental Figure 5: LSC snRNA-seq cell subtype annotations.**

**A)** i) PCA and UMAP plot of neurons. Each data point represents one nucleus, color-coded by assigned cell subtype, RNA counts per nucleus, or site of symptom onset. ii) Dot plot showing expression of the most differentially expressed neuron-associated transcripts for each subtype (identified using Scanpy's ranking method). Color indicates mean gene expression, and dot size denotes the proportion of cells with non-zero counts, from 0-100%.

**B – F)** Same as A, shown for different cell types.

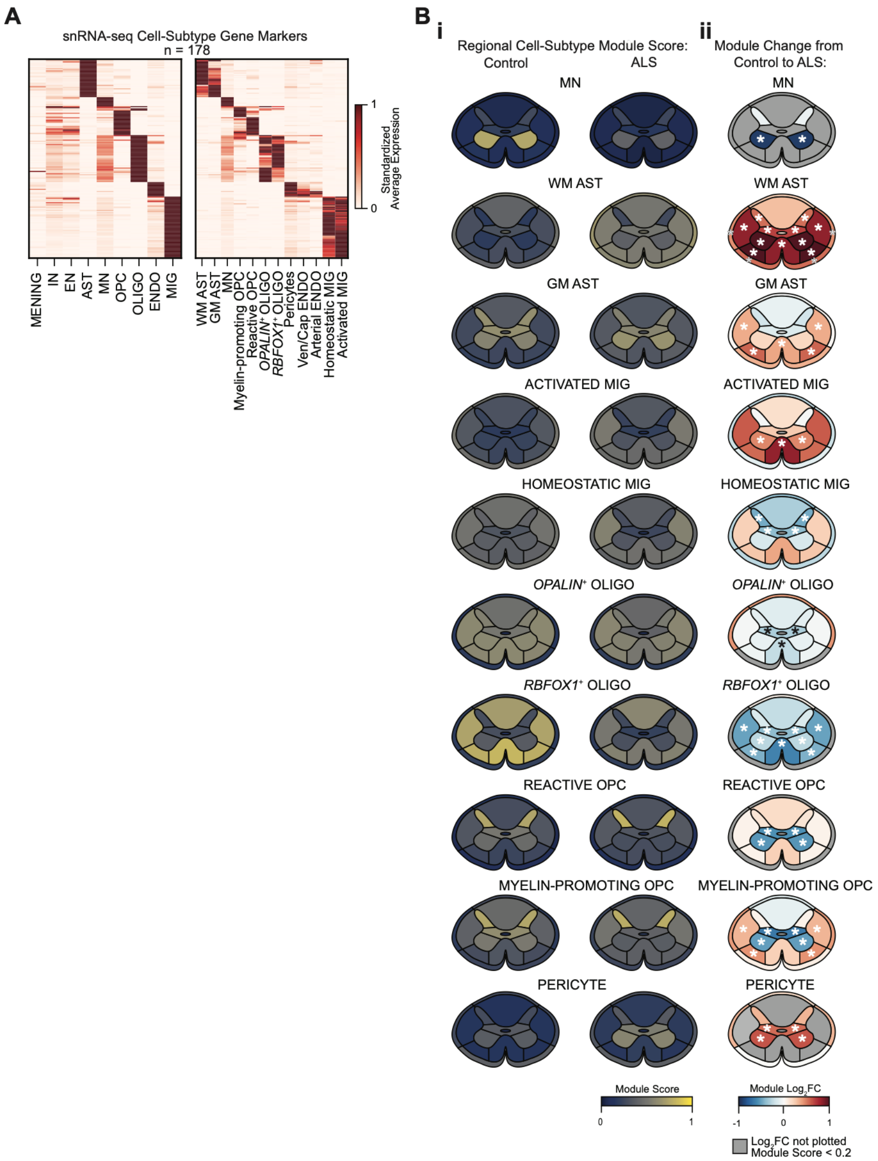
A
snRNA-seq Cell-Subtype Gene Markers
n = 178
Standardized Average Expression
1
0
MENING
IN
EN
AST
MN
OPC
OLIGO
ENDO
MIG
WM AST
GM AST
MN
Myelin-promoting OPC
Reactive OPC
OPALIN+ OLIGO
RBFOX1+ OLIGO
Pericytes
Ven/Cap ENDO
Arterial ENDO
Homeostatic MIG
Activated MIG
B
i
Regional Cell-Subtype Module Score:
Control
ALS
ii
Module Change from Control to ALS:
MN
WM AST
GM AST
ACTIVATED MIG
HOMEOSTATIC MIG
OPALIN+ OLIGO
RBFOX1+ OLIGO
REACTIVE OPC
MYELIN-PROMOTING OPC
PERICYTE
Module Score
0
1
Module Log2FC
-1
0
1
Log2FC not plotted
Module Score < 0.2

**Supplemental Figure 6: Spatial expression of LSC snRNA-seq cell subtypes.**

**A)** Min-maxed scaled expression of marker genes across all LSC cell types (left) and cell subtypes (right). Marker genes are differentially expressed within each cell subtype (i.e. genes differentially expressed in homeostatic MG relative to reactive MIG) and are used to infer the spatial distribution of cell subtypes.

**B)** (i) Spatial expression of cell subtype module scores across all control (left) and ALS (right) arrays. (ii) $Log_2$ fold change of cell subtype module scores in ALS relative to controls. Regions with module scores less than 0.2 are masked in $log_2$ fold change panels. Asterisks indicate significant differences relative to control donors (Welch's t-test, $p < 0.05$, FDR-BY).

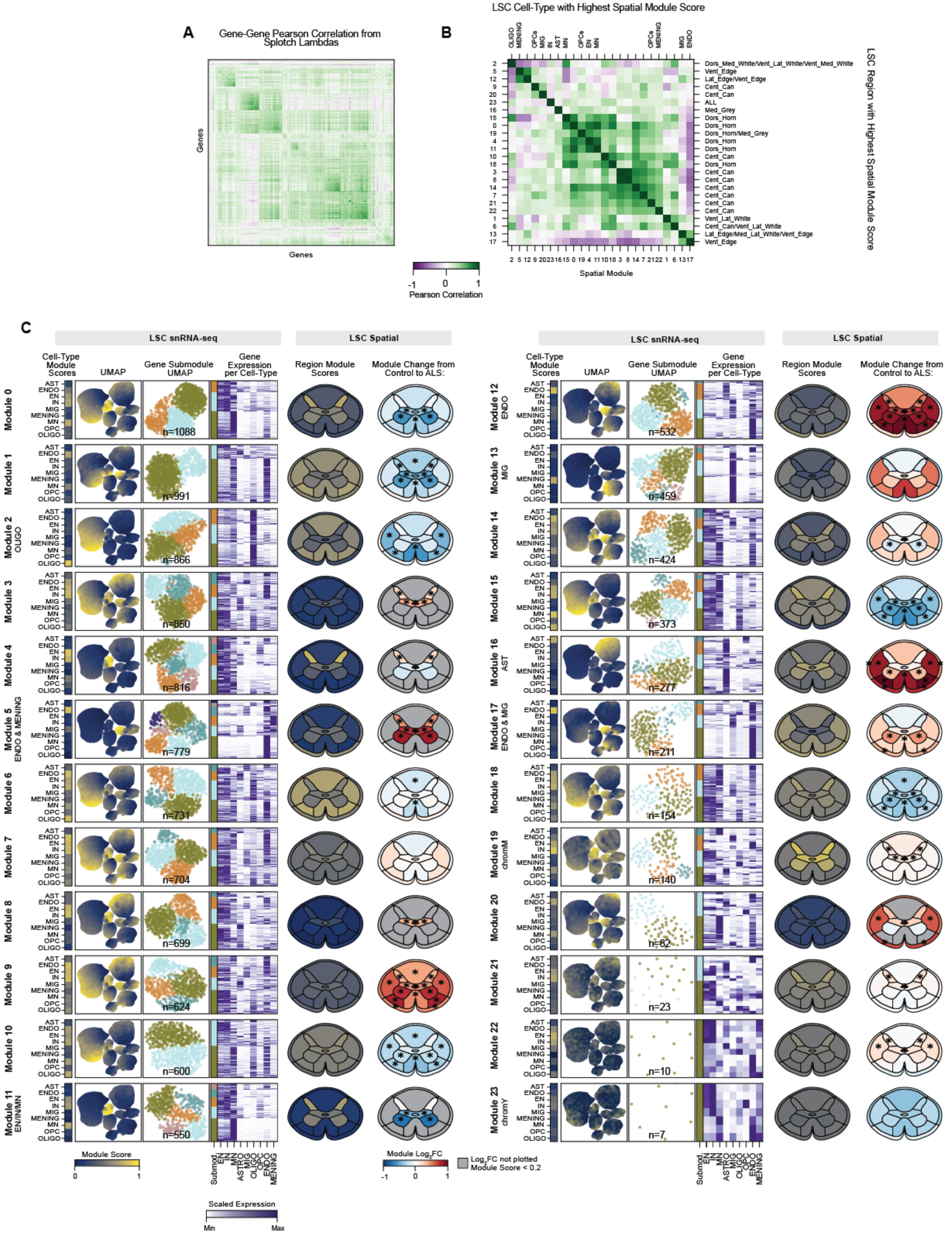

A
Gene-Gene Pearson Correlation from Splotch Lambdas
Genes
Genes
-1
0
1
Pearson Correlation
B
LSC Cell-Type with Highest Spatial Module Score
LSC Region with Highest Spatial Module Score
Spatial Module
Dors_Med_White/Vent_Lat_White/Vent_Med_White
Vent_Edge
Lat_Edge/Vent_Edge
Cent_Can
ALL
Med_Grey
Dors_Horn
Dors_Horn/Med_Grey
Vent_Lat_White
Cent_Can/Vent_Lat_White
Lat_Edge/Med_Lat_White/Vent_Edge
C
LSC snRNA-seq
LSC Spatial
Cell-Type Module Scores
UMAP
Gene Submodule UMAP
Gene Expression per Cell-Type
Region Module Scores
Module Change from Control to ALS:
Module 0
Module 1
Module 2 OLIGO
Module 3
Module 4
Module 5 ENDO & MENING
Module 6
Module 7
Module 8
Module 9
Module 10
Module 11 EN/IN/MN
Module 12 ENDO
Module 13 MIG
Module 14
Module 15
Module 16 AST
Module 17 ENDO & MIG
Module 18
Module 19 chromM
Module 20
Module 21
Module 22
Module 23 chromY
n=1088
n=991
n=866
n=850
n=816
n=779
n=731
n=704
n=699
n=624
n=600
n=550
n=532
n=459
n=424
n=373
n=277
n=211
n=154
n=140
n=82
n=23
n=10
n=7
AST
ENDO
EN
IN
MIG
MENING
MN
OPC
OLIGO
Module Score
0
1
Module Log2FC
-1
0
1
Log2FC not plotted Module Score < 0.2
Scaled Expression
Min
Max

**Supplemental Figure 7: LSC Spatial gene modules show cell type and region specificity and are altered in ALS.**

**A)** Pearson correlation between Splotch estimates of spot-level spatial expression (lambdas).

**B)** Pearson correlation between spatial module scores across all LSC ST arrays. The top x-axis is labelled with the snRNA-seq cell type that has the highest relative expression for each module. The right y-axis is labelled with the spinal region that has the highest relative expression of the module.

**C)** Overview of each LSC spatial gene module. Left to right: 1) Average spatial module score for each snRNA-seq cell type; 2) snRNA-seq UMAP plot, with each dot corresponding to an individual nucleus colored by its calculated spatial module score; 3) gene UMAP plot of all genes in spatial module from a *k*-NN graph (10 neighbors, snRNA-seq gene correlation distance), with each dot corresponding to an individual gene colored by assigned submodule; 4) average standardized gene expression for each snRNA-seq cell type, with submodule assignment annotated; 5) from ST data, the average spatial module scores per region of spinal cord; 6) $\log_2$ fold change of module scores in ALS donors relative to controls. Regions with module scores less than 0.2 are masked in $\log_2$ fold change panels. Asterisks indicate significant differences relative to control donors (Welch's t-test, $p < 0.05$, FDR-BY).

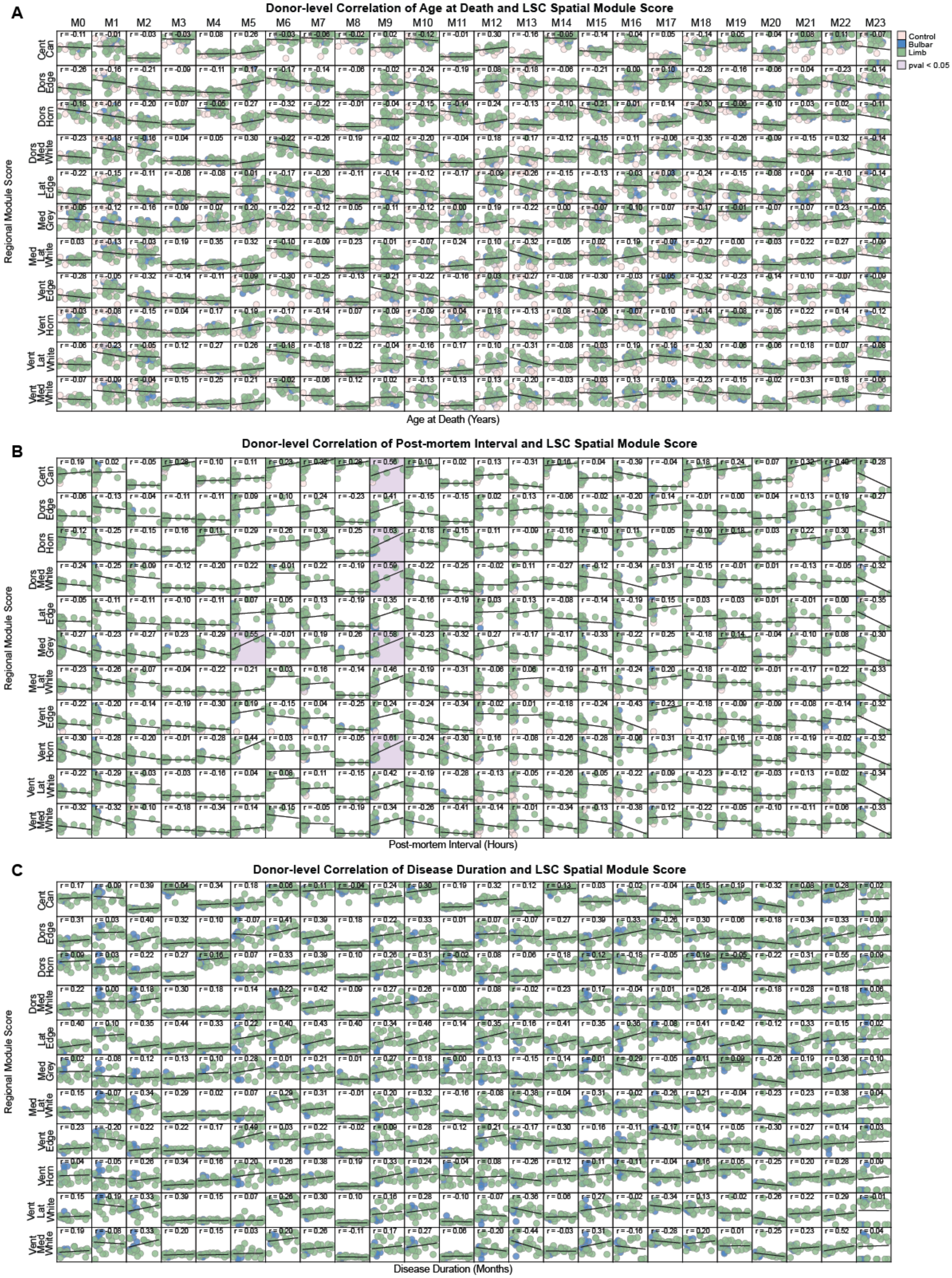
A
Donor-level Correlation of Age at Death and LSC Spatial Module Score
M0 M1 M2 M3 M4 M5 M6 M7 M8 M9 M10 M11 M12 M13 M14 M15 M16 M17 M18 M19 M20 M21 M22 M23
Control
Bulbar
Limb
pval < 0.05
Regional Module Score
Cent Can
Dors Edge
Dors Horn
Dors Med White
Lat Edge
Med Grey
Med Lat White
Vent Edge
Vent Horn
Vent Lat White
Vent Med White
Age at Death (Years)
B
Donor-level Correlation of Post-mortem Interval and LSC Spatial Module Score
Regional Module Score
Post-mortem Interval (Hours)
C
Donor-level Correlation of Disease Duration and LSC Spatial Module Score
Regional Module Score
Disease Duration (Months)

**Supplemental Figure 8: LSC Spatial gene module 9 is affected by post-mortem interval.**

The averaged spatial modules scores per AAR for each donor sample in the LSC ST data, plotted on the y-axis, compared to **A)** donor age at death, **B)** post-mortem interval (PMI), and **C)** ALS disease duration plotted on the x-axis. Each dot represents an individual donor. Plots include a least squares regression line and Pearson correlation coefficient, and are colored violet when correlations are significantly different from 0 (Wald test, $p < 0.05$, FDR-BH).

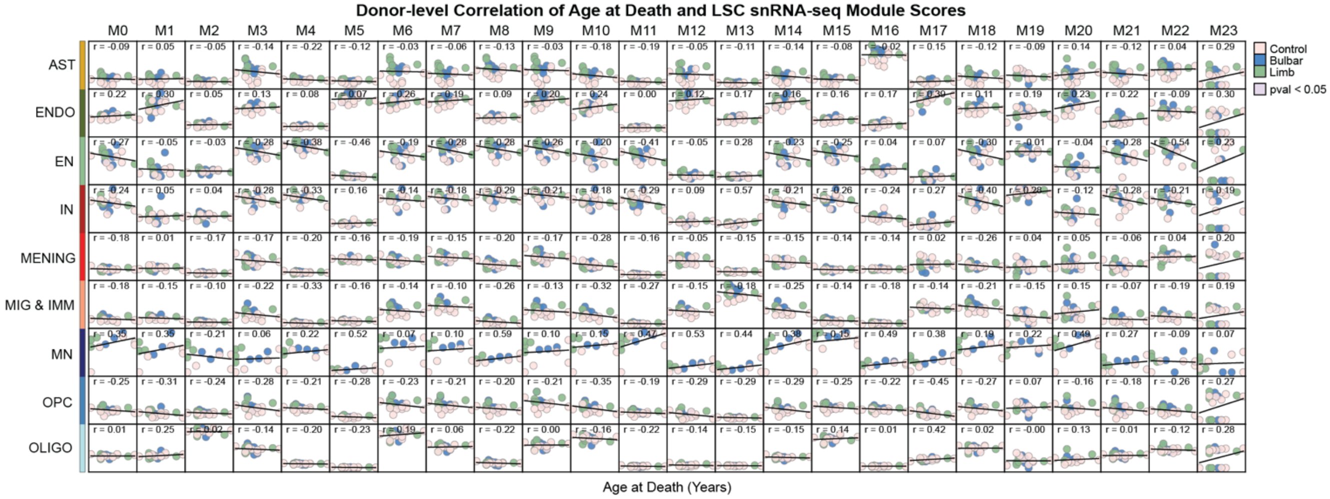

Donor-level Correlation of Age at Death and LSC snRNA-seq Module Scores
M0 M1 M2 M3 M4 M5 M6 M7 M8 M9 M10 M11 M12 M13 M14 M15 M16 M17 M18 M19 M20 M21 M22 M23
AST
ENDO
EN
IN
MENING
MIG & IMM
MN
OPC
OLIGO
Control
Bulbar
Limb
pval < 0.05
Age at Death (Years)

**Supplemental Figure 9: No significant correlations detected between donor-level metadata and LSC snRNA-seq spatial gene module scores.**

Age at death plotted against snRNA-seq module scores per LSC cell type. Each dot represents an individual donor. Plots include a least squares regression line and Pearson correlation coefficient, and are colored violet when correlations are significantly different from 0 (Wald test, $p < 0.05$, FDR-BH).

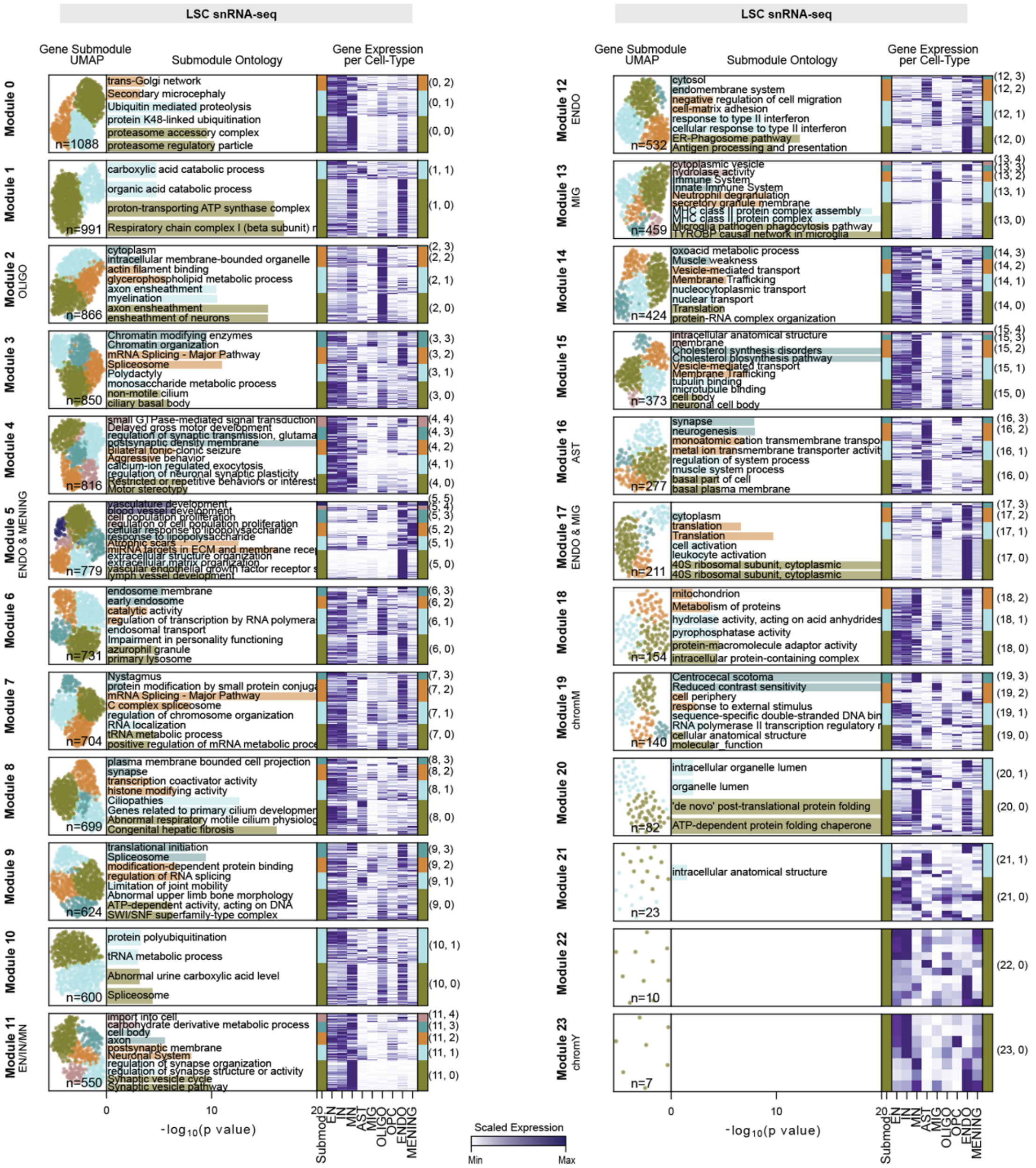

LSC snRNA-seq
Gene Submodule UMAP
Submodule Ontology
Gene Expression per Cell-Type
Module 0
trans-Golgi network
Secondary microcephaly
Ubiquitin mediated proteolysis
protein K48-linked ubiquitination
proteasome accessory complex
proteasome regulatory particle
n=1088
(0, 2) (0, 1) (0, 0)
Module 1
carboxylic acid catabolic process
organic acid catabolic process
proton-transporting ATP synthase complex
Respiratory chain complex I (beta subunit) r
n=991
(1, 1) (1, 0)
Module 2 OLIGO
cytoplasm
intracellular membrane-bounded organelle
actin filament binding
glycerophospholipid metabolic process
axon ensheathment
myelination
axon ensheathment
ensheathment of neurons
n=866
(2, 3) (2, 2) (2, 1) (2, 0)
Module 3
Chromatin modifying enzymes
Chromatin organization
mRNA Splicing - Major Pathway
Spliceosome
Polydactyly
monosaccharide metabolic process
non-motile cilium
ciliary basal body
n=850
(3, 3) (3, 2) (3, 1) (3, 0)
Module 4
small GTPase-mediated signal transduction
Delayed gross motor development
regulation of synaptic transmission, glutama
postsynaptic density membrane
Bilateral tonic-clonic seizure
Aggressive behavior
calcium-ion regulated exocytosis
regulation of neuronal synaptic plasticity
Restricted or repetitive behaviors or interest
Motor stereotypy
n=816
(4, 4) (4, 3) (4, 2) (4, 1) (4, 0)
Module 5 ENDO & MENING
vasculature development
blood vessel development
cell population proliferation
regulation of cell population proliferation
cellular response to lipopolysaccharide
response to lipopolysaccharide
Atrophic scars
miRNA targets in ECM and membrane recep
extracellular structure organization
extracellular matrix organization
vascular endothelial growth factor receptor s
lymph vessel development
n=779
(5, 5) (5, 4) (5, 3) (5, 2) (5, 1) (5, 0)
Module 6
endosome membrane
early endosome
catalytic activity
regulation of transcription by RNA polymera
endosomal transport
Impairment in personality functioning
azurophil granule
primary lysosome
n=731
(6, 3) (6, 2) (6, 1) (6, 0)
Module 7
Nystagmus
protein modification by small protein conjuga
mRNA Splicing - Major Pathway
C complex spliceosome
regulation of chromosome organization
RNA localization
tRNA metabolic process
positive regulation of mRNA metabolic proce
n=704
(7, 3) (7, 2) (7, 1) (7, 0)
Module 8
plasma membrane bounded cell projection
synapse
transcription coactivator activity
histone modifying activity
Ciliopathies
Genes related to primary cilium developmen
Abnormal respiratory motile cilium physiolog
Congenital hepatic fibrosis
n=699
(8, 3) (8, 2) (8, 1) (8, 0)
Module 9
translational initiation
Spliceosome
modification-dependent protein binding
regulation of RNA splicing
Limitation of joint mobility
Abnormal upper limb bone morphology
ATP-dependent activity, acting on DNA
SWI/SNF superfamily-type complex
n=624
(9, 3) (9, 2) (9, 1) (9, 0)
Module 10
protein polyubiquitination
tRNA metabolic process
Abnormal urine carboxylic acid level
Spliceosome
n=600
(10, 1) (10, 0)
Module 11 EN/IN/MN
import into cell
carbohydrate derivative metabolic process
cell body
axon
postsynaptic membrane
Neuronal System
regulation of synapse organization
regulation of synapse structure or activity
Synaptic vesicle cycle
Synaptic vesicle pathway
n=550
(11, 4) (11, 3) (11, 2) (11, 1) (11, 0)
0 10 20
-log10(p value)
Submod EN IN MN AST MIG OLIGO OPC ENDO MENING
Module 12 ENDO
cytosol
endomembrane system
negative regulation of cell migration
cell-matrix adhesion
response to type II interferon
cellular response to type II interferon
ER-Phagosome pathway
Antigen processing and presentation
n=532
(12, 3) (12, 2) (12, 1) (12, 0)
Module 13 MIG
cytoplasmic vesicle
hydrolase activity
Immune System
Innate Immune System
Neutrophil degranulation
secretory granule membrane
MHC class II protein complex assembly
MHC class II protein complex
Microglia pathogen phagocytosis pathway
TYROBP causal network in microglia
n=459
(13, 4) (13, 3) (13, 2) (13, 1) (13, 0)
Module 14
oxoacid metabolic process
Muscle weakness
Vesicle-mediated transport
Membrane Trafficking
nucleocytoplasmic transport
nuclear transport
Translation
protein-RNA complex organization
n=424
(14, 3) (14, 2) (14, 1) (14, 0)
Module 15
intracellular anatomical structure
membrane
Cholesterol synthesis disorders
Cholesterol biosynthesis pathway
Vesicle-mediated transport
Membrane Trafficking
tubulin binding
microtubule binding
cell body
neuronal cell body
n=373
(15, 4) (15, 3) (15, 2) (15, 1) (15, 0)
Module 16 AST
synapse
neurogenesis
monoatomic cation transmembrane transpo
metal ion transmembrane transporter activit
regulation of system process
muscle system process
basal part of cell
basal plasma membrane
n=277
(16, 3) (16, 2) (16, 1) (16, 0)
Module 17 ENDO & MIG
cytoplasm
translation
Translation
cell activation
leukocyte activation
40S ribosomal subunit, cytoplasmic
40S ribosomal subunit, cytoplasmic
n=211
(17, 3) (17, 2) (17, 1) (17, 0)
Module 18
mitochondrion
Metabolism of proteins
hydrolase activity, acting on acid anhydrides
pyrophosphatase activity
protein-macromolecule adaptor activity
intracellular protein-containing complex
n=154
(18, 2) (18, 1) (18, 0)
Module 19 chromM
Centrocecal scotoma
Reduced contrast sensitivity
cell periphery
response to external stimulus
sequence-specific double-stranded DNA bin
RNA polymerase II transcription regulatory r
cellular anatomical structure
molecular_function
n=140
(19, 3) (19, 2) (19, 1) (19, 0)
Module 20
intracellular organelle lumen
organelle lumen
'de novo' post-translational protein folding
ATP-dependent protein folding chaperone
n=82
(20, 1) (20, 0)
Module 21
intracellular anatomical structure
n=23
(21, 1) (21, 0)
Module 22
n=10
(22, 0)
Module 23 chromY
n=7
(23, 0)
Scaled Expression
Min Max

**Supplemental Figure 10: Using snRNA-seq correlations to identify cell type submodules from LSC spatial gene modules.**

Overview of spatial submodules. Left to right: 1) gene UMAP plot of all genes in spatial module from a *k*-NN graph (10 neighbors, snRNA-seq gene correlation distance), with each dot corresponding to an individual gene colored by assigned submodule; 2) GO terms associated with each submodule, with length of underlying bar plot indicating $-\log_{10}(p\text{-value})$; 3) average standardized gene expression for each LSC snRNA-seq cell type, with submodule assignment annotated.

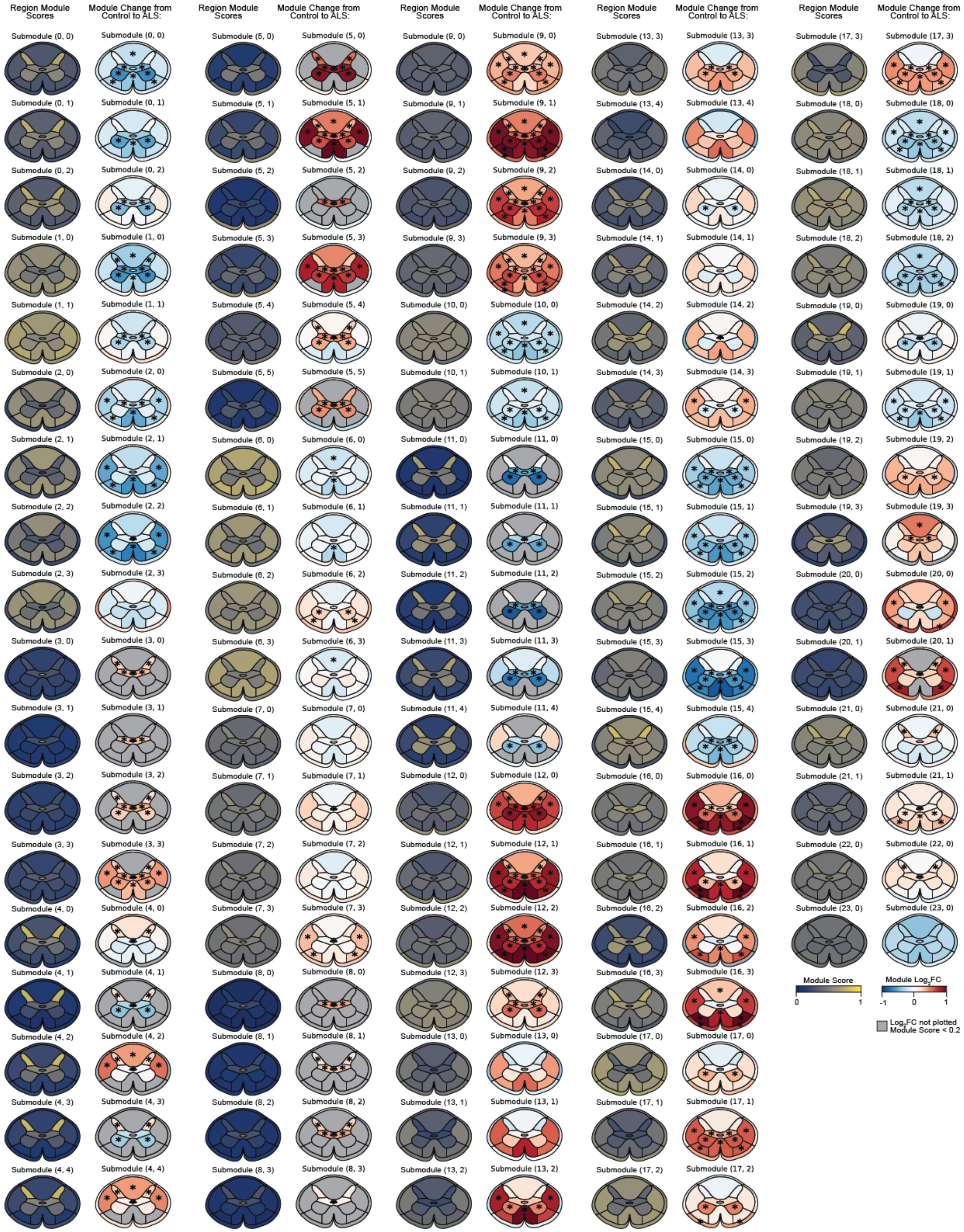
Region Module Scores
Module Change from Control to ALS:
Submodule (0, 0)
Submodule (0, 1)
Submodule (0, 2)
Submodule (1, 0)
Submodule (1, 1)
Submodule (2, 0)
Submodule (2, 1)
Submodule (2, 2)
Submodule (2, 3)
Submodule (3, 0)
Submodule (3, 1)
Submodule (3, 2)
Submodule (3, 3)
Submodule (4, 0)
Submodule (4, 1)
Submodule (4, 2)
Submodule (4, 3)
Submodule (4, 4)
Submodule (5, 0)
Submodule (5, 1)
Submodule (5, 2)
Submodule (5, 3)
Submodule (5, 4)
Submodule (5, 5)
Submodule (6, 0)
Submodule (6, 1)
Submodule (6, 2)
Submodule (6, 3)
Submodule (7, 0)
Submodule (7, 1)
Submodule (7, 2)
Submodule (7, 3)
Submodule (8, 0)
Submodule (8, 1)
Submodule (8, 2)
Submodule (8, 3)
Submodule (9, 0)
Submodule (9, 1)
Submodule (9, 2)
Submodule (9, 3)
Submodule (10, 0)
Submodule (10, 1)
Submodule (11, 0)
Submodule (11, 1)
Submodule (11, 2)
Submodule (11, 3)
Submodule (11, 4)
Submodule (12, 0)
Submodule (12, 1)
Submodule (12, 2)
Submodule (12, 3)
Submodule (13, 0)
Submodule (13, 1)
Submodule (13, 2)
Submodule (13, 3)
Submodule (13, 4)
Submodule (14, 0)
Submodule (14, 1)
Submodule (14, 2)
Submodule (14, 3)
Submodule (15, 0)
Submodule (15, 1)
Submodule (15, 2)
Submodule (15, 3)
Submodule (15, 4)
Submodule (16, 0)
Submodule (16, 1)
Submodule (16, 2)
Submodule (16, 3)
Submodule (17, 0)
Submodule (17, 1)
Submodule (17, 2)
Submodule (17, 3)
Submodule (18, 0)
Submodule (18, 1)
Submodule (18, 2)
Submodule (19, 0)
Submodule (19, 1)
Submodule (19, 2)
Submodule (19, 3)
Submodule (20, 0)
Submodule (20, 1)
Submodule (21, 0)
Submodule (21, 1)
Submodule (22, 0)
Submodule (23, 0)
Module Score
0
1
Module Log2FC
-1
0
1
Log2FC not plotted
Module Score < 0.2

**Supplemental Figure 11: LSC Spatial submodules show region specificity and are altered in ALS.**

The average spatial submodule scores per spinal cord region (left). $Log_2$ fold change of submodule scores in ALS donors relative to controls (right). Regions with submodule scores less than 0.2 are masked in $log_2$ fold change panels. Asterisks indicate significant differences relative to control donors (Welch's t-test, $p < 0.05$, FDR-BY).

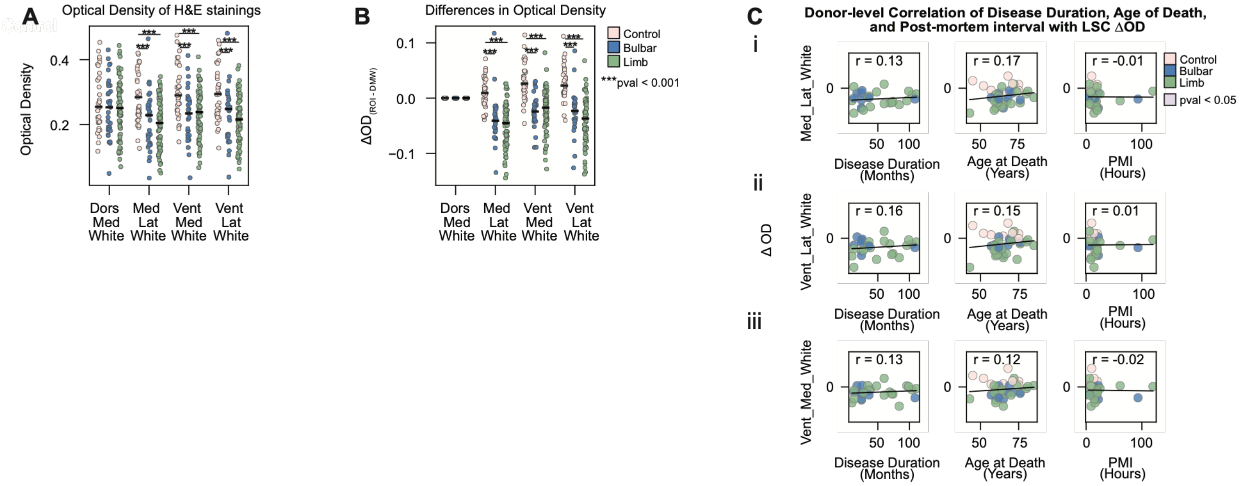

A
Optical Density of H&E stainings
Optical Density
0.4
0.2
***
Dors Med White
Med Lat White
Vent Med White
Vent Lat White
B
Differences in Optical Density
ΔOD(ROI - DMW)
0.1
0.0
-0.1
Control
Bulbar
Limb
***pval < 0.001
C
Donor-level Correlation of Disease Duration, Age of Death, and Post-mortem interval with LSC ΔOD
i
ii
iii
Δ OD
Med_Lat_White
Vent_Lat_White
Vent_Med_White
r = 0.13
r = 0.17
r = -0.01
r = 0.16
r = 0.15
r = 0.01
r = 0.13
r = 0.12
r = -0.02
pval < 0.05
Disease Duration (Months)
Age at Death (Years)
PMI (Hours)
50
100
75
0

**Supplemental Figure 12: Optical density is not influenced by disease duration, age at death, or post-mortem interval.**

**A)** Average scaled optical density (OD) measurement per specified LSC anatomical region. Each dot represents an individual array, colored by site of symptom onset. The black line indicates the median OD for each group. (Welch's t-test, $p < 0.05$, FDR-BY).

**B)** Average scaled OD relative to DMW, indicating the magnitude of OD loss for that region of interest (ROI). Each dot represents an individual array, colored by site of symptom onset. The black line indicates the median OD for each group. (Welch's t-test, $p < 0.05$, FDR-BY).

**C)** Correlation of disease duration, age at death, or PMI with ΔOD for the i) medial lateral white, ii) ventral lateral white, and iii) ventral medial white matter regions. Each dot represents an individual donor, colored by site of symptom onset. Plots include a least squares regression line and Pearson correlation coefficient, and are colored violet when correlations are significantly different from 0 (Wald test, $p < 0.05$, FDR-BH).

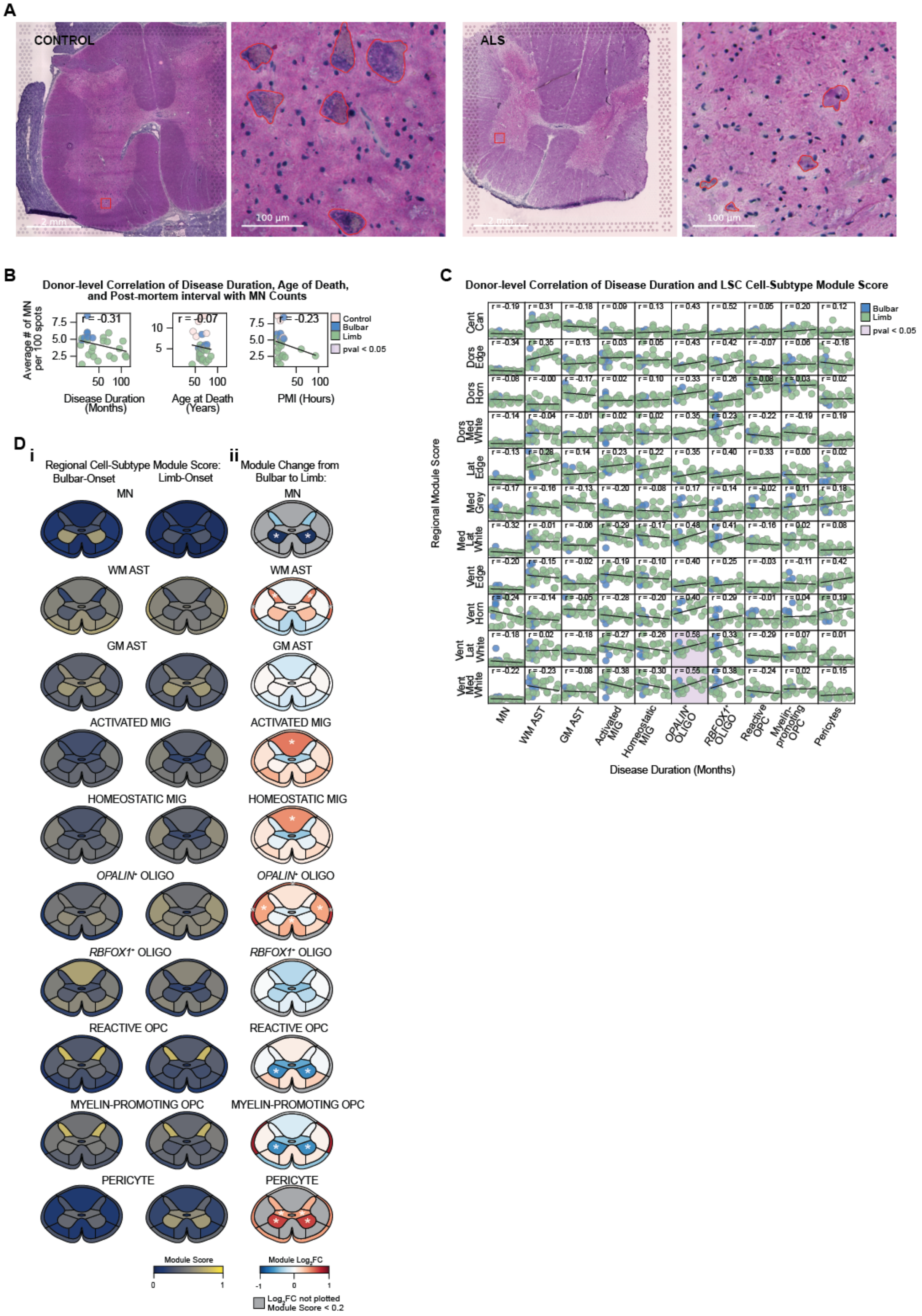
A
CONTROL
ALS
2 mm
100 µm
B
Donor-level Correlation of Disease Duration, Age of Death, and Post-mortem interval with MN Counts
Average # of MN per 100 spots
r = -0.31
r = -0.07
r = -0.23
Control
Bulbar
Limb
pval < 0.05
Disease Duration (Months)
Age at Death (Years)
PMI (Hours)
C
Donor-level Correlation of Disease Duration and LSC Cell-Subtype Module Score
Bulbar
Limb
pval < 0.05
Regional Module Score
Cent Can
Dors Edge
Dors Horn
Dors Med White
Lat Edge
Med Grey
Med Lat White
Vent Edge
Vent Horn
Vent Lat White
Vent Med White
MN
WM AST
GM AST
Activated MIG
Homeostatic MIG
OPALIN+ OLIGO
RBFOX1+ OLIGO
Reactive OPC
Myelin-promoting OPC
Pericytes
Disease Duration (Months)
D i
Regional Cell-Subtype Module Score:
Bulbar-Onset
Limb-Onset
ii
Module Change from Bulbar to Limb:
MN
WM AST
GM AST
ACTIVATED MIG
HOMEOSTATIC MIG
OPALIN+ OLIGO
RBFOX1+ OLIGO
REACTIVE OPC
MYELIN-PROMOTING OPC
PERICYTE
Module Score
0
1
Module Log2FC
-1
0
1
Log2FC not plotted
Module Score < 0.2

**Supplemental Figure 13: Glial composition is largely unaffected by site of symptom onset.**

**A)** Representative Visium H&E images with a red box outlining a 1000 by 1000 pixel (px) region (left), which is magnified on the right to show motor neurons outlined in red.

**B)** Donor-level correlation between the average ventral horn motor neuron count and disease duration, age at death, and post-mortem interval. Each dot represents an individual donor and is color-coded according to site of symptom onset. Plots are annotated with least squares regression line, Pearson correlation coefficient, and shaded in violet if correlation is significantly different from zero (Wald test, $p < 0.05$, FDR-BH).

**C)** Donor-level correlation between the cell subtype gene expression module score per LSC AAR and disease duration. Each dot represents an individual donor and is color-coded according to site of symptom onset. Plots are annotated with least squares regression line, Pearson correlation coefficient, and shaded in violet if correlation is significantly different from zero (Wald test, $p < 0.05$, FDR-BH).

**D)** (i) Spatial expression of cell subtype module scores across all bulbar-onset (left) and limb-onset (right) arrays. (ii) $Log_2$ fold change of cell subtype module scores in limb relative to bulbar-onset. Regions with module scores less than 0.2 are masked in $log_2$ fold change panels. Asterisks indicate significant differences relative to control donors (Welch's t-test, $p < 0.05$, FDR-BY).

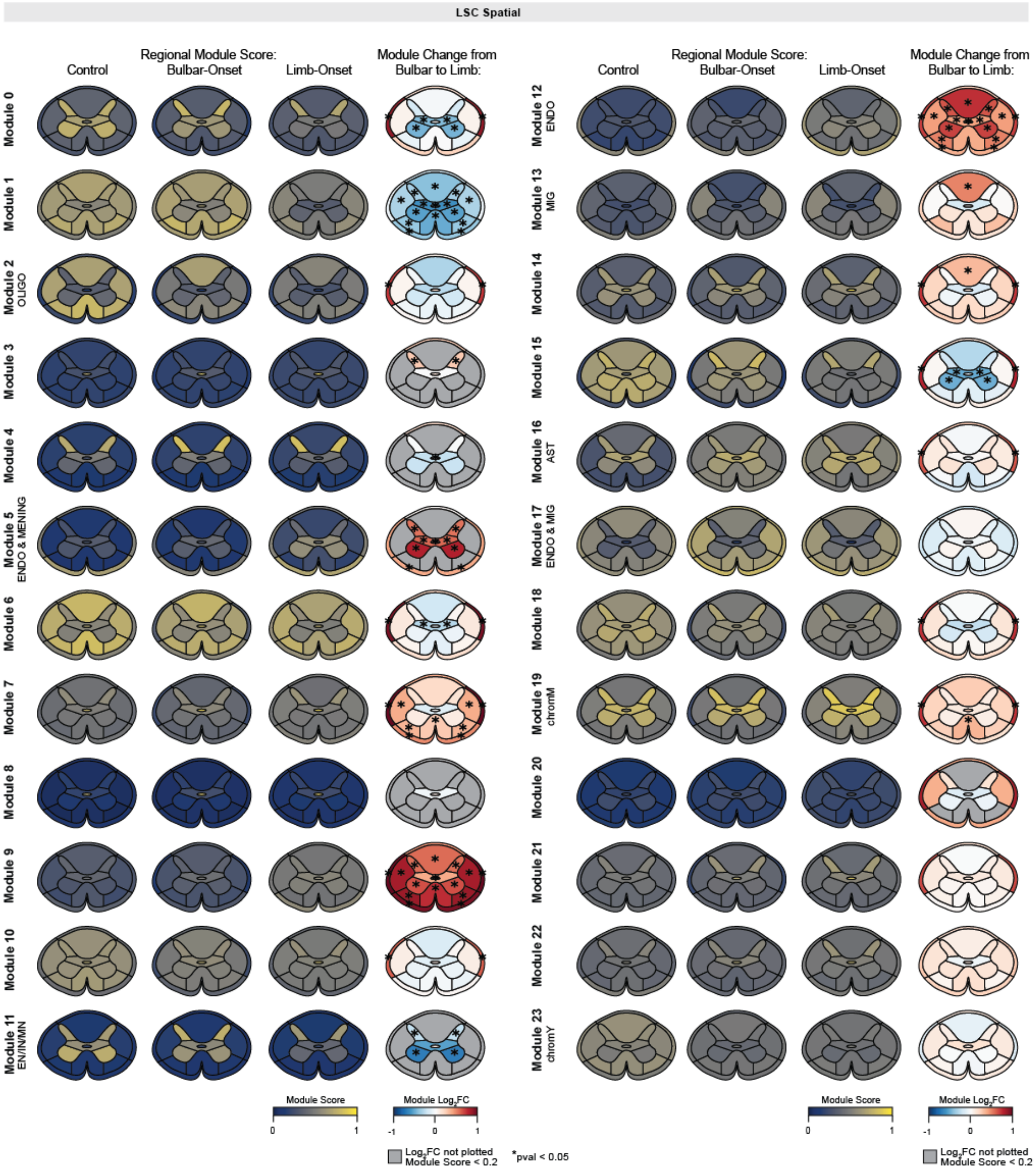

LSC Spatial
Control
Regional Module Score:
Bulbar-Onset
Limb-Onset
Module Change from
Bulbar to Limb:
Module 0
Module 1
Module 2
OLIGO
Module 3
Module 4
Module 5
ENDO & MENING
Module 6
Module 7
Module 8
Module 9
Module 10
Module 11
EN/IN/MN
Module 12
ENDO
Module 13
MIG
Module 14
Module 15
Module 16
AST
Module 17
ENDO & MIG
Module 18
Module 19
chromM
Module 20
Module 21
Module 22
Module 23
chromY
Module Score
0
1
Module Log2FC
-1
0
1
Log2FC not plotted
Module Score < 0.2
*pval < 0.05

**Supplemental Figure 14: Expression of LSC spatial gene modules stratified by site of symptom onset.**

Left to right: (i) The average spatial gene module scores per spinal cord region across all control (left) and ALS (right) arrays. (ii) $\text{Log}_2$ fold change of spatial gene module scores in ALS relative to controls. (iii) The average spatial gene module scores per spinal cord region across all bulbar-onset (left) and limb-onset (right) arrays. (iv) $\text{Log}_2$ fold change of spatial gene module scores in limb relative to bulbar-onset. Regions with module scores less than 0.2 are masked in $\log_2$ fold change panels. Asterisks indicate significant differences relative to control donors (Welch's t-test, $p < 0.05$, FDR-BY).

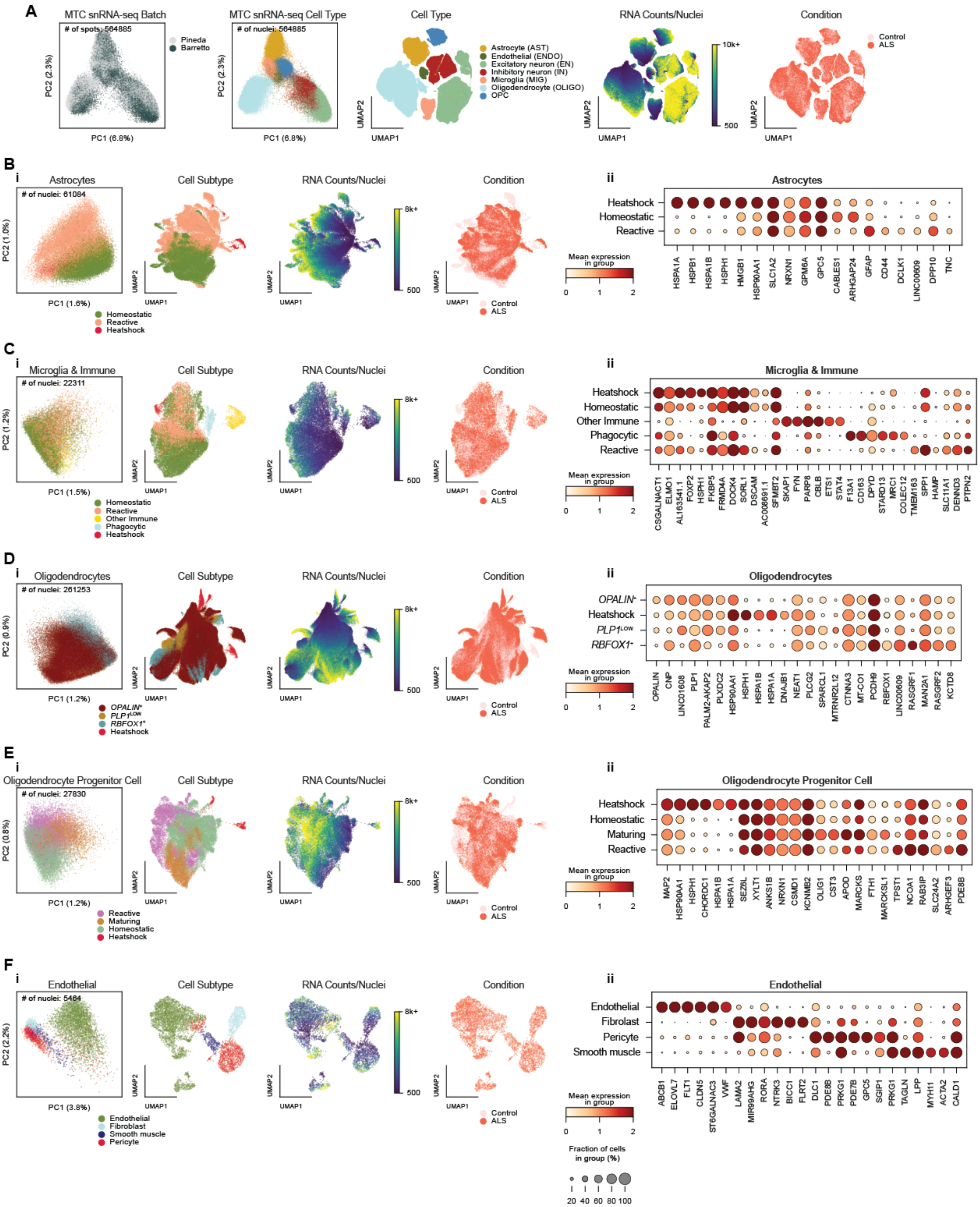
A
MTC snRNA-seq Batch
### of spots: 564885
Pineda
Barretto
PC2 (2.3%)
PC1 (6.8%)
MTC snRNA-seq Cell Type
### of nuclei: 564885
Cell Type
Astrocyte (AST)
Endothelial (ENDO)
Excitatory neuron (EN)
Inhibitory neuron (IN)
Microglia (MIG)
Oligodendrocyte (OLIGO)
OPC
UMAP2
UMAP1
RNA Counts/Nuclei
10k+
500
Condition
Control
ALS
B
i
Astrocytes
### of nuclei: 61084
PC2 (1.0%)
PC1 (1.6%)
Homeostatic
Reactive
Heatshock
Cell Subtype
RNA Counts/Nuclei
8k+
500
Condition
Control
ALS
ii
Astrocytes
Heatshock
Homeostatic
Reactive
Mean expression in group
0 1 2
HSPA1A HSPB1 HSPA1B HSPH1 HMGB1 HSP90AA1 SLC1A2 NRXN1 GPM6A GPC5 CABLES1 ARHGAP24 GFAP CD44 DCLK1 LINC00609 DPP10 TNC
C
i
Microglia & Immune
### of nuclei: 22311
PC2 (1.2%)
PC1 (1.5%)
Homeostatic
Reactive
Other Immune
Phagocytic
Heatshock
Cell Subtype
RNA Counts/Nuclei
Condition
ii
Microglia & Immune
Heatshock
Homeostatic
Other Immune
Phagocytic
Reactive
CSGALNACT1 ELMO1 AL163541.1 FOXP2 HSPH1 FKBP5 FRMD4A DOCK4 SORL1 DSCAM AC008691.1 SFMBT2 SKAP1 FYN PARP8 CBLB ETS1 STAT4 F13A1 CD163 DPYD STARD13 MRC1 COLEC12 TMEM163 SPP1 HAMP SLC11A1 DENND3 PTPN2
D
i
Oligodendrocytes
### of nuclei: 261253
PC2 (0.9%)
PC1 (1.2%)
OPALIN+
PLP1LOW
RBFOX1+
Heatshock
ii
Oligodendrocytes
OPALIN+
Heatshock
PLP1LOW
RBFOX1+
OPALIN CNP LINC01608 PLP1 PALM2-AKAP2 PLXDC2 HSP90AA1 HSPH1 HSPA1B HSPA1A DNAJB1 NEAT1 PLCG2 SPARCL1 MTRNR2L12 CTNNA3 MT-CO1 PCDH9 RBFOX1 LINC00609 RASGRF1 MAN2A1 RASGRF2 KCTD8
E
i
Oligodendrocyte Progenitor Cell
### of nuclei: 27830
PC2 (0.8%)
PC1 (1.2%)
Reactive
Maturing
Homeostatic
Heatshock
ii
Oligodendrocyte Progenitor Cell
Heatshock
Homeostatic
Maturing
Reactive
MAP2 HSP90AA1 HSPH1 CHORDC1 HSPA1B HSPA1A SEZ6L XYLT1 ANKS1B NRXN1 CSMD1 KCNMB2 OLIG1 CST3 APOD MARCKS FTH1 MARCKSL1 TPST1 NCOA1 RAB3IP SLC24A2 ARHGEF3 PDE8B
F
i
Endothelial
### of nuclei: 5464
PC2 (2.2%)
PC1 (3.8%)
Endothelial
Fibroblast
Smooth muscle
Pericyte
ii
Endothelial
Endothelial
Fibroblast
Pericyte
Smooth muscle
ABCB1 ELOVL7 FLT1 CLDN5 ST6GALNAC3 VWF LAMA2 MIR99AHG RORA NTRK3 BICC1 FLRT2 DLC1 PDE8B PRKG1 PDE7B GPC5 SGIP1 PRKG1 TAGLN LPP MYH11 ACTA2 CALD1
Fraction of cells in group (%)
20 40 60 80 100

**Supplemental Figure 15: Integration of MTC snRNA-seq data with Pineda et al.**

**A)** PCA and UMAP plots of integrated snRNA-seq data, with each dot corresponding to an individual nucleus colored by batch (Barretto or Pineda et al.), cell type, UMI counts per nucleus, and condition.

**B)** i) PCA and UMAP plot of astrocytes. Each data point represents one nucleus, color-coded by assigned cell subtype, RNA counts per nucleus, or condition. ii) Dot plot showing expression of the most differentially expressed neuron-associated transcripts for each subtype (identified using Scanpy's ranking method). Color indicates mean gene expression, and dot size denotes the proportion of cells with non-zero counts, from 0-100%.

**C – F)** Same as A, shown for different cell types.

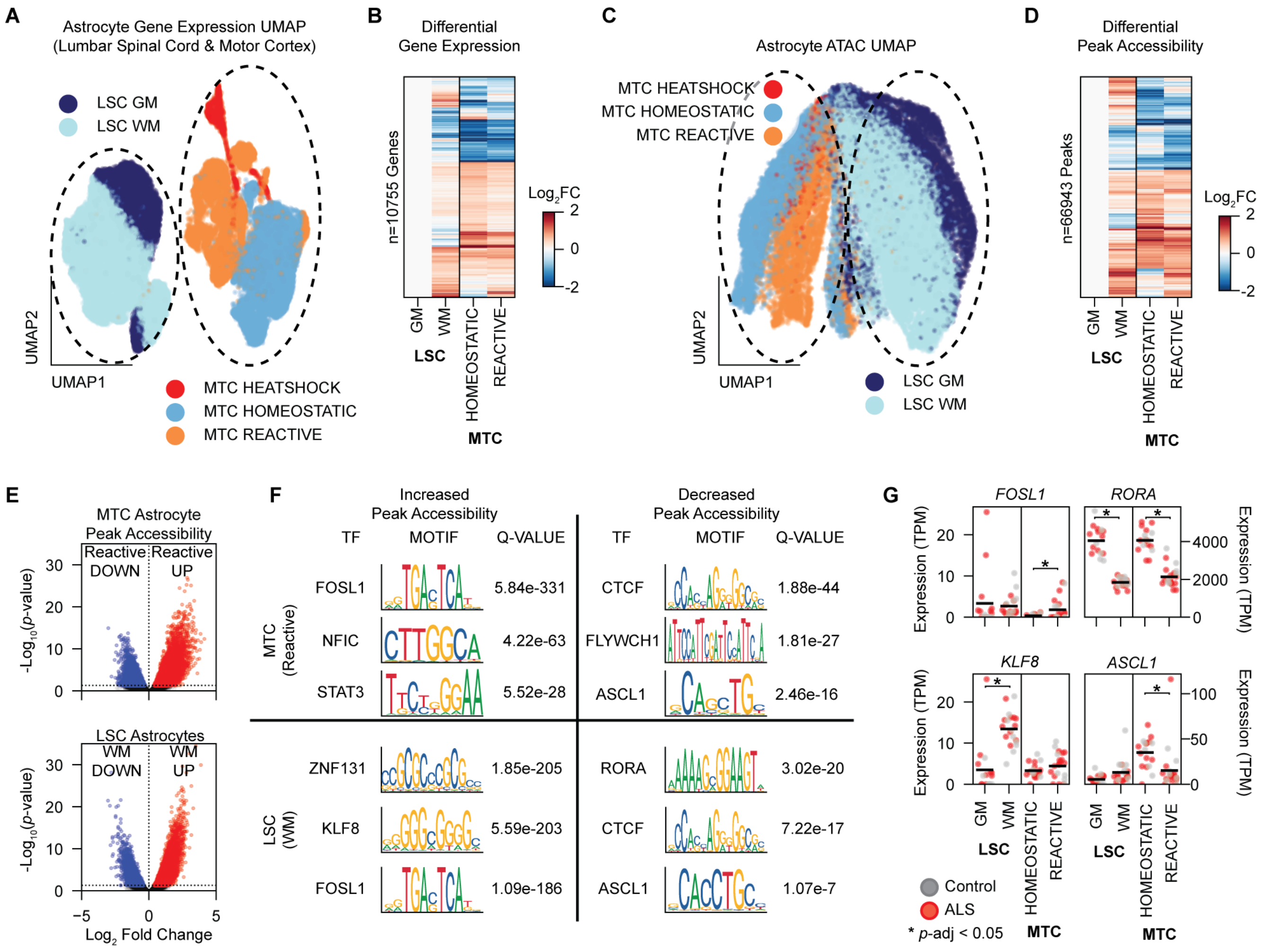

A
Astrocyte Gene Expression UMAP
(Lumbar Spinal Cord & Motor Cortex)
LSC GM
LSC WM
UMAP2
UMAP1
MTC HEATSHOCK
MTC HOMEOSTATIC
MTC REACTIVE
B
Differential
Gene Expression
n=10755 Genes
Log2FC
2
0
-2
GM
WM
LSC
HOMEOSTATIC
REACTIVE
MTC
C
Astrocyte ATAC UMAP
MTC HEATSHOCK
MTC HOMEOSTATIC
MTC REACTIVE
UMAP2
UMAP1
LSC GM
LSC WM
D
Differential
Peak Accessibility
n=66943 Peaks
Log2FC
2
0
-2
GM
WM
LSC
HOMEOSTATIC
REACTIVE
MTC
E
MTC Astrocyte
Peak Accessibility
Reactive DOWN
Reactive UP
-Log10(p-value)
LSC Astrocytes
WM DOWN
WM UP
-Log10(p-value)
Log2 Fold Change
F
Increased
Peak Accessibility
TF
MOTIF
Q-VALUE
Decreased
Peak Accessibility
TF
MOTIF
Q-VALUE
MTC
(Reactive)
FOSL1 5.84e-331
NFIC 4.22e-63
STAT3 5.52e-28
CTCF 1.88e-44
FLYWCH1 1.81e-27
ASCL1 2.46e-16
LSC
(WM)
ZNF131 1.85e-205
KLF8 5.59e-203
FOSL1 1.09e-186
RORA 3.02e-20
CTCF 7.22e-17
ASCL1 1.07e-7
G
FOSL1
RORA
KLF8
ASCL1
Expression (TPM)
Expression (TPM)
GM
WM
LSC
HOMEOSTATIC
REACTIVE
MTC
Control
ALS
* p-adj < 0.05

**Supplemental Figure 16: Comparison of Astrocytes between MTC and LSC.**

**A)** UMAP plot of astrocyte snRNA-seq gene expression, colored by tissue of origin and annotated astrocyte function.

**B)** $Log_2$ fold change of differentially expressed (pseudobulked DESeq2) genes, compared to LSC GM astrocytes.

**C)** UMAP plot of astrocyte snATAC-seq accessible chromatin, colored as in **A**.

**D)** $Log_2$ fold change of differentially accessible (pseudobulked DESeq2) ATAC peaks, compared to LSC GM astrocytes.

**E)** Volcano plot of differentially expressed genes, comparing reactive astrocytes to homeostatic astrocytes in the MTC (top) and comparing WM to GM astrocytes in the LSC (bottom).

**F)** Selected motifs enriched in peaks that are more accessible (left) or less accessible (right) in Reactive or WM astrocytes.

**G)** Expression of selected transcription factors in different astrocyte functional groups, pseudobulked and colored by control vs ALS. Asterisks indicate significant difference (pseudobulked DESeq2; Wald test, adjusted $p$-value < 0.05).

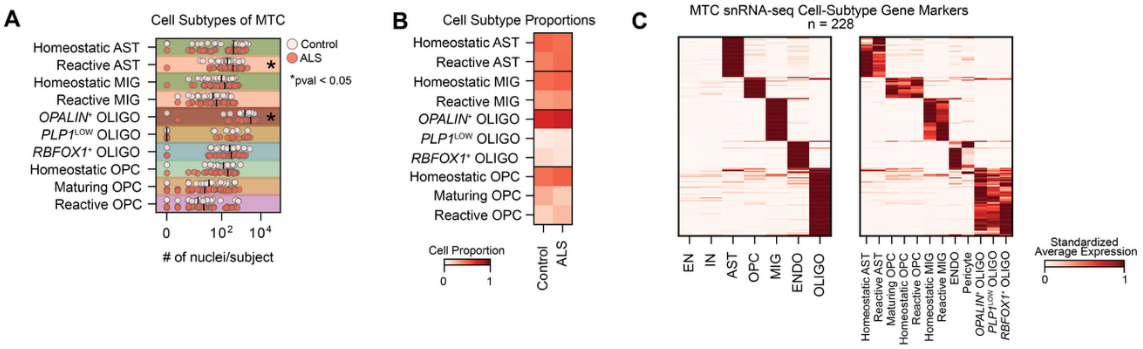
A
Cell Subtypes of MTC
Control
ALS
*pval < 0.05
Homeostatic AST
Reactive AST
Homeostatic MIG
Reactive MIG
OPALIN+ OLIGO
PLP1LOW OLIGO
RBFOX1+ OLIGO
Homeostatic OPC
Maturing OPC
Reactive OPC
0
10²
10⁴
### of nuclei/subject
B
Cell Subtype Proportions
Homeostatic AST
Reactive AST
Homeostatic MIG
Reactive MIG
OPALIN+ OLIGO
PLP1LOW OLIGO
RBFOX1+ OLIGO
Homeostatic OPC
Maturing OPC
Reactive OPC
Cell Proportion
0
1
Control
ALS
C
MTC snRNA-seq Cell-Subtype Gene Markers
n = 228
EN
IN
AST
OPC
MIG
ENDO
OLIGO
Homeostatic AST
Reactive AST
Maturing OPC
Homeostatic OPC
Reactive OPC
Homeostatic MIG
Reactive MIG
ENDO
Pericyte
OPALIN+ OLIGO
PLP1LOW OLIGO
RBFOX1+ OLIGO
Standardized
Average Expression
0
1

**Supplemental Figure 17: MTC snRNA-seq cell subtypes.**

**A)** Number of nuclei annotated per cell subtype across all snRNA-seq libraries, color-coded by disease status. Black bars mark median nuclei counts; asterisks denote significant differences between ALS and control donors (Welch's t-test, $p < 0.05$, FDR-BH).

**B)** Averaged proportions of cell subtypes for controls and ALS (Welch's t-test, $p < 0.05$, FDR-BH).

**C)** Min-maxed scaled expression of marker genes across MTC cell subtypes.

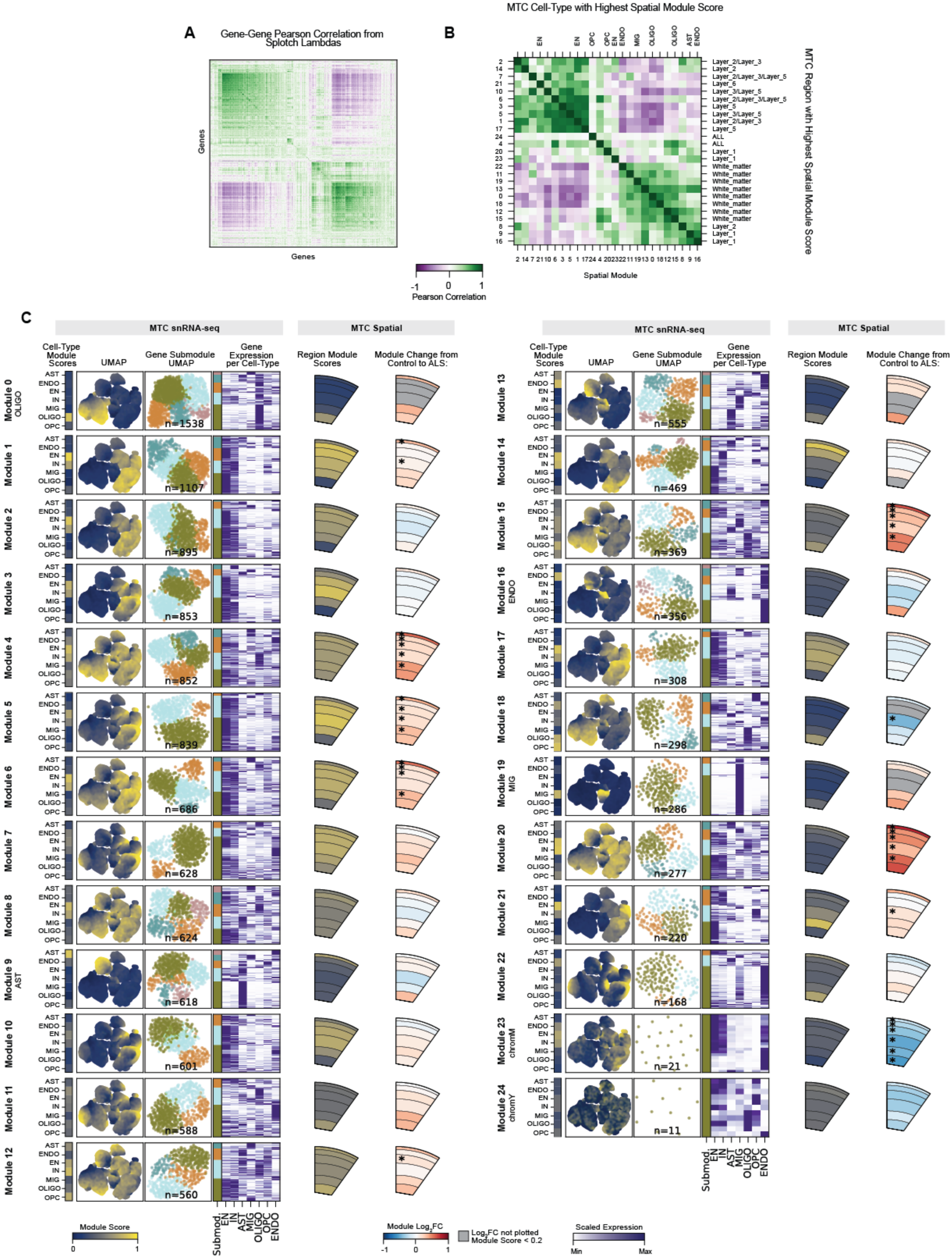
A
Gene-Gene Pearson Correlation from Splotch Lambdas
Genes
Genes
-1 0 1
Pearson Correlation
B
MTC Cell-Type with Highest Spatial Module Score
MTC Region with Highest Spatial Module Score
Spatial Module
C
MTC snRNA-seq
MTC Spatial
Cell-Type Module Scores
UMAP
Gene Submodule UMAP
Gene Expression per Cell-Type
Region Module Scores
Module Change from Control to ALS:
Module Score
Module Log2FC
Log2FC not plotted Module Score < 0.2
Scaled Expression
Min
Max

**Supplemental Figure 18: MTC Spatial gene modules show cell type and region specificity, and are altered in ALS.**

**A)** Pearson correlation between Splotch estimates of spot-level spatial expression (lambdas).

**B)** Pearson correlation between spatial module scores across all MTC ST arrays. The top x-axis is labelled with the snRNA-seq cell type that has the highest relative expression for each module. The right y-axis is labelled with the spinal region that has the highest relative expression of the module.

**C)** Overview of each MTC spatial gene module. Left to right: 1) Average spatial module score for each snRNA-seq cell type; 2) snRNA-seq UMAP plot, with each dot corresponding to an individual nucleus colored by its calculated spatial module score; 3) gene UMAP plot of all genes in spatial module from a *k*-NN graph (10 neighbors, snRNA-seq gene correlation distance), with each dot corresponding to an individual gene colored by assigned submodule; 4) average standardized gene expression for each snRNA-seq cell type, with submodule assignment annotated; 5) from ST data, the average spatial module scores per region of spinal cord; 6) $\log_2$ fold change of module scores in ALS donors relative to controls. Regions with module scores less than 0.2 are masked in $\log_2$ fold change panels. Asterisks indicate significant differences relative to control donors (Welch's t-test, $p < 0.05$, FDR-BY).

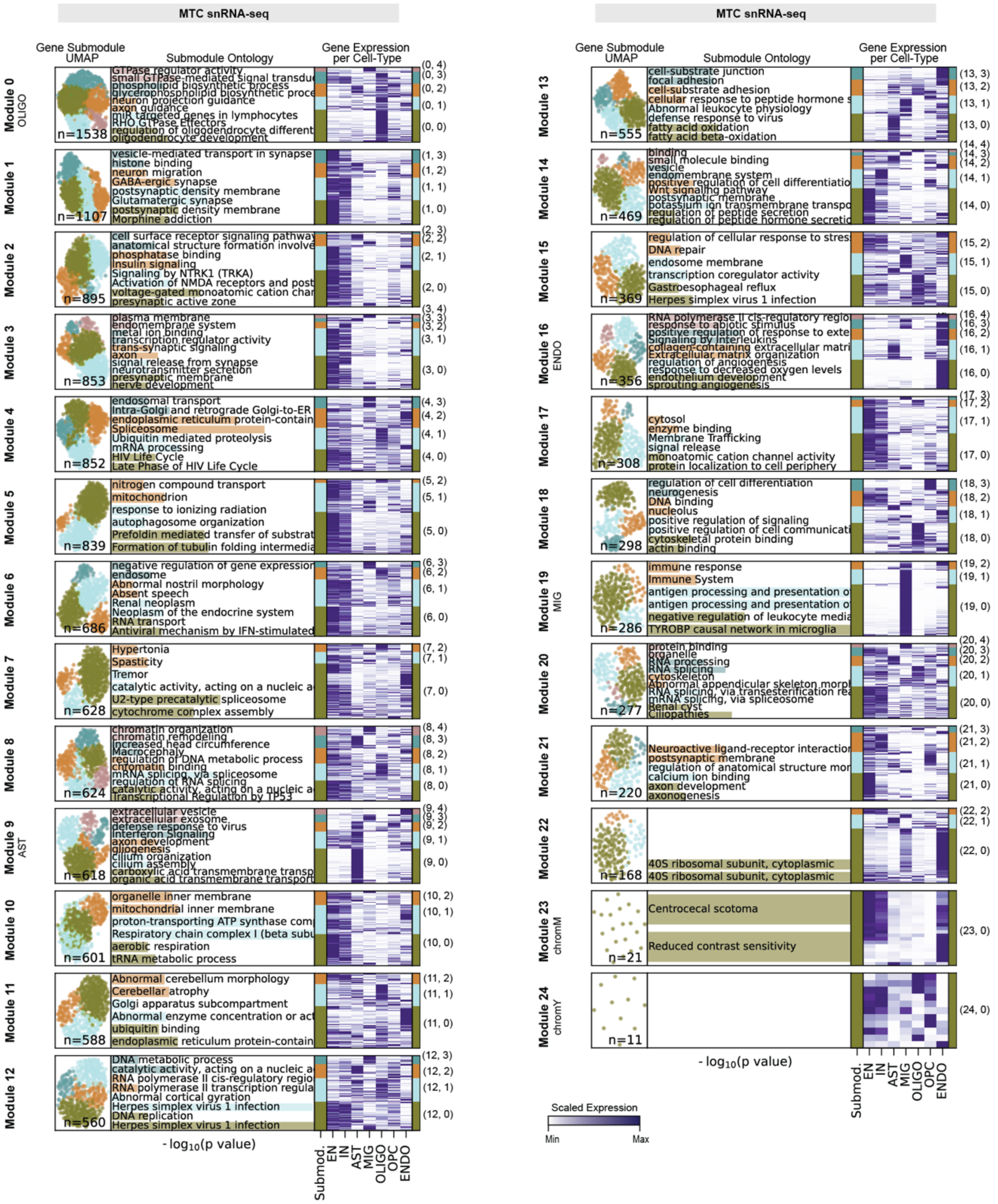
MTC snRNA-seq
Gene Submodule UMAP
Submodule Ontology
Gene Expression per Cell-Type
Module 0 OLIGO
n=1538
Module 1
n=1107
Module 2
n=895
Module 3
n=853
Module 4
n=852
Module 5
n=839
Module 6
n=686
Module 7
n=628
Module 8
n=624
Module 9 AST
n=618
Module 10
n=601
Module 11
n=588
Module 12
n=560
- log10(p value)
Submod.
EN
IN
AST
MIG
OLIGO
OPC
ENDO
MTC snRNA-seq
Module 13
n=555
Module 14
n=469
Module 15
n=369
Module 16 ENDO
n=356
Module 17
n=308
Module 18
n=298
Module 19 MIG
n=286
Module 20
n=277
Module 21
n=220
Module 22
n=168
Module 23 chromM
n=21
Centrocecal scotoma
Reduced contrast sensitivity
Module 24 chromY
n=11
Scaled Expression
Min
Max

**Supplemental Figure 19: Using snRNA-seq correlations to identify cell type submodules from MTC spatial gene modules.**

Overview of spatial submodules. Left to right: 1) gene UMAP plot of all genes in spatial module from a *k*-NN graph (10 neighbors, snRNA-seq gene correlation distance), with each dot corresponding to an individual gene colored by assigned submodule; 2) GO terms associated with each submodule, with length of underlying bar plot indicating $-\log_{10}(p\text{-value})$; 3) average standardized gene expression for each MTC snRNA-seq cell type, with submodule assignment annotated.

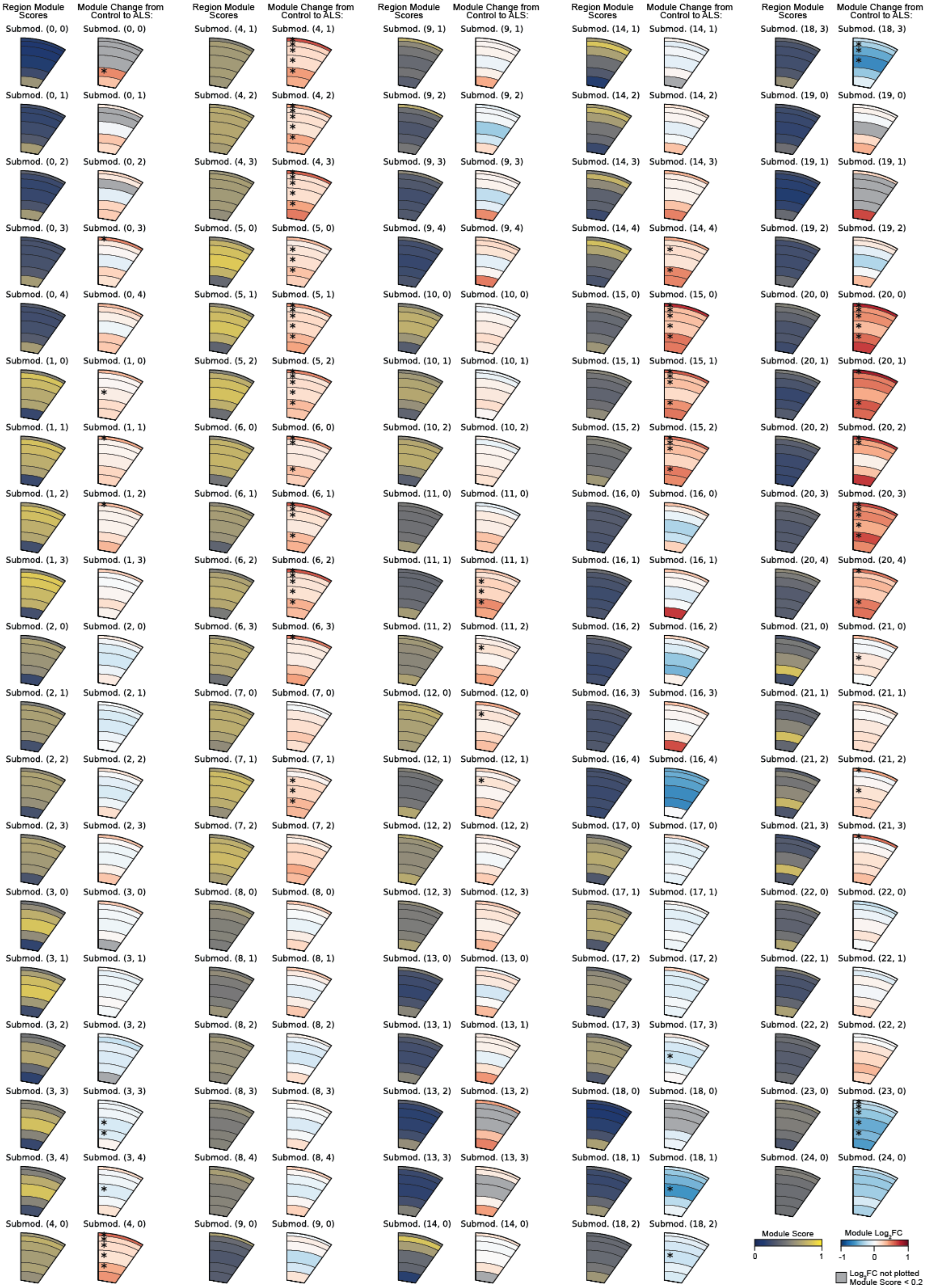

Region Module Scores
Module Change from Control to ALS:
Submod. (0, 0) Submod. (0, 0)
Submod. (0, 1) Submod. (0, 1)
Submod. (0, 2) Submod. (0, 2)
Submod. (0, 3) Submod. (0, 3)
Submod. (0, 4) Submod. (0, 4)
Submod. (1, 0) Submod. (1, 0)
Submod. (1, 1) Submod. (1, 1)
Submod. (1, 2) Submod. (1, 2)
Submod. (1, 3) Submod. (1, 3)
Submod. (2, 0) Submod. (2, 0)
Submod. (2, 1) Submod. (2, 1)
Submod. (2, 2) Submod. (2, 2)
Submod. (2, 3) Submod. (2, 3)
Submod. (3, 0) Submod. (3, 0)
Submod. (3, 1) Submod. (3, 1)
Submod. (3, 2) Submod. (3, 2)
Submod. (3, 3) Submod. (3, 3)
Submod. (3, 4) Submod. (3, 4)
Submod. (4, 0) Submod. (4, 0)
Region Module Scores
Module Change from Control to ALS:
Submod. (4, 1) Submod. (4, 1)
Submod. (4, 2) Submod. (4, 2)
Submod. (4, 3) Submod. (4, 3)
Submod. (5, 0) Submod. (5, 0)
Submod. (5, 1) Submod. (5, 1)
Submod. (5, 2) Submod. (5, 2)
Submod. (6, 0) Submod. (6, 0)
Submod. (6, 1) Submod. (6, 1)
Submod. (6, 2) Submod. (6, 2)
Submod. (6, 3) Submod. (6, 3)
Submod. (7, 0) Submod. (7, 0)
Submod. (7, 1) Submod. (7, 1)
Submod. (7, 2) Submod. (7, 2)
Submod. (8, 0) Submod. (8, 0)
Submod. (8, 1) Submod. (8, 1)
Submod. (8, 2) Submod. (8, 2)
Submod. (8, 3) Submod. (8, 3)
Submod. (8, 4) Submod. (8, 4)
Submod. (9, 0) Submod. (9, 0)
Region Module Scores
Module Change from Control to ALS:
Submod. (9, 1) Submod. (9, 1)
Submod. (9, 2) Submod. (9, 2)
Submod. (9, 3) Submod. (9, 3)
Submod. (9, 4) Submod. (9, 4)
Submod. (10, 0) Submod. (10, 0)
Submod. (10, 1) Submod. (10, 1)
Submod. (10, 2) Submod. (10, 2)
Submod. (11, 0) Submod. (11, 0)
Submod. (11, 1) Submod. (11, 1)
Submod. (11, 2) Submod. (11, 2)
Submod. (12, 0) Submod. (12, 0)
Submod. (12, 1) Submod. (12, 1)
Submod. (12, 2) Submod. (12, 2)
Submod. (12, 3) Submod. (12, 3)
Submod. (13, 0) Submod. (13, 0)
Submod. (13, 1) Submod. (13, 1)
Submod. (13, 2) Submod. (13, 2)
Submod. (13, 3) Submod. (13, 3)
Submod. (14, 0) Submod. (14, 0)
Region Module Scores
Module Change from Control to ALS:
Submod. (14, 1) Submod. (14, 1)
Submod. (14, 2) Submod. (14, 2)
Submod. (14, 3) Submod. (14, 3)
Submod. (14, 4) Submod. (14, 4)
Submod. (15, 0) Submod. (15, 0)
Submod. (15, 1) Submod. (15, 1)
Submod. (15, 2) Submod. (15, 2)
Submod. (16, 0) Submod. (16, 0)
Submod. (16, 1) Submod. (16, 1)
Submod. (16, 2) Submod. (16, 2)
Submod. (16, 3) Submod. (16, 3)
Submod. (16, 4) Submod. (16, 4)
Submod. (17, 0) Submod. (17, 0)
Submod. (17, 1) Submod. (17, 1)
Submod. (17, 2) Submod. (17, 2)
Submod. (17, 3) Submod. (17, 3)
Submod. (18, 0) Submod. (18, 0)
Submod. (18, 1) Submod. (18, 1)
Submod. (18, 2) Submod. (18, 2)
Region Module Scores
Module Change from Control to ALS:
Submod. (18, 3) Submod. (18, 3)
Submod. (19, 0) Submod. (19, 0)
Submod. (19, 1) Submod. (19, 1)
Submod. (19, 2) Submod. (19, 2)
Submod. (20, 0) Submod. (20, 0)
Submod. (20, 1) Submod. (20, 1)
Submod. (20, 2) Submod. (20, 2)
Submod. (20, 3) Submod. (20, 3)
Submod. (20, 4) Submod. (20, 4)
Submod. (21, 0) Submod. (21, 0)
Submod. (21, 1) Submod. (21, 1)
Submod. (21, 2) Submod. (21, 2)
Submod. (21, 3) Submod. (21, 3)
Submod. (22, 0) Submod. (22, 0)
Submod. (22, 1) Submod. (22, 1)
Submod. (22, 2) Submod. (22, 2)
Submod. (23, 0) Submod. (23, 0)
Submod. (24, 0) Submod. (24, 0)
Module Score
0
1
Module $Log_2FC$
-1
0
1
$Log_2FC$ not plotted
Module Score < 0.2

**Supplemental Figure 20: MTC Spatial submodules show region specificity and are altered in ALS.**

The average spatial submodule scores per motor cortex layer (left). $Log_2$ fold change of submodule scores in ALS donors relative to controls (right). Regions with submodule scores less than 0.2 are masked in $log_2$ fold change panels. Asterisks indicate significant differences relative to control donors (Welch's t-test, $p < 0.05$, FDR-BY).

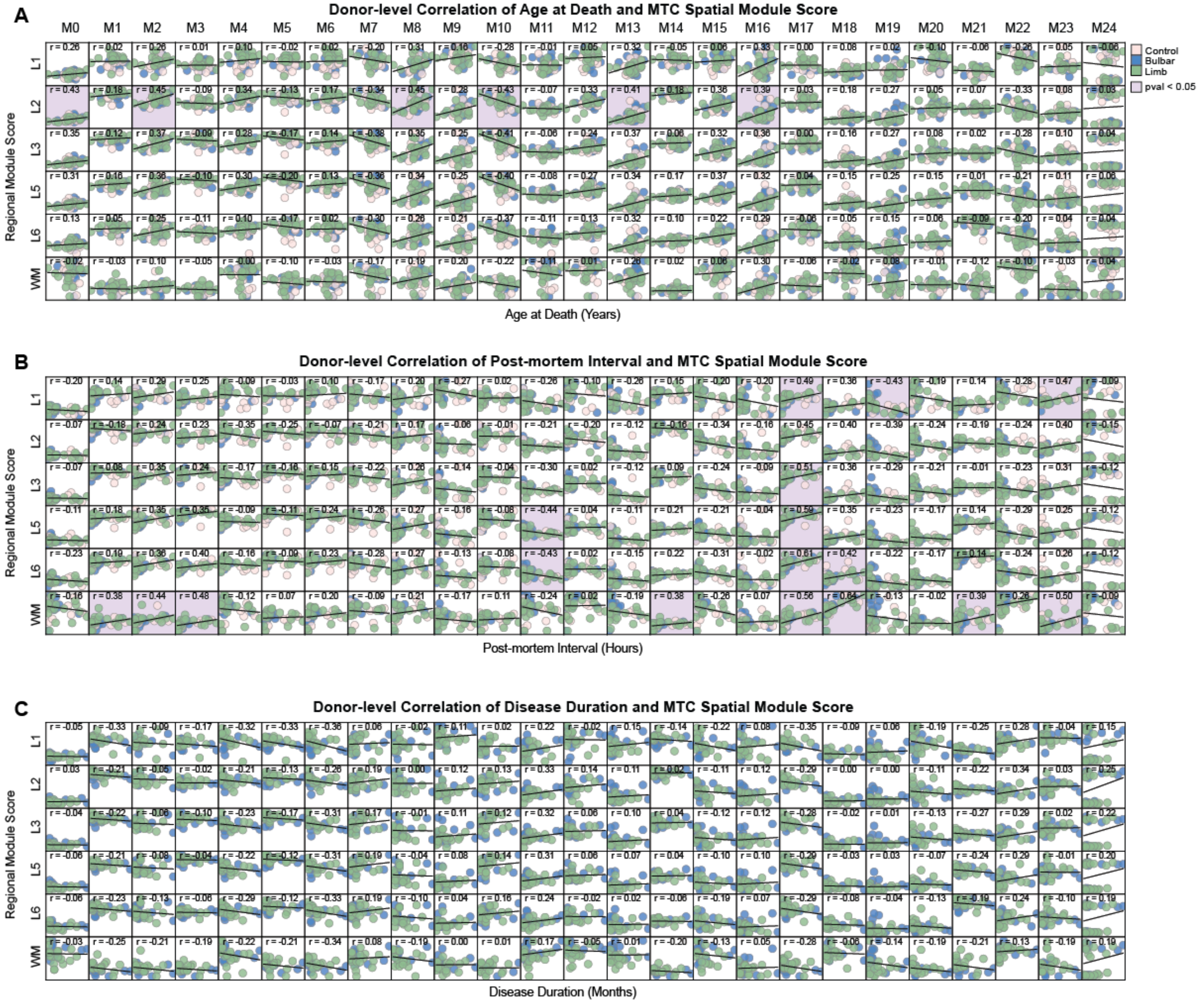
A
Donor-level Correlation of Age at Death and MTC Spatial Module Score
M0 M1 M2 M3 M4 M5 M6 M7 M8 M9 M10 M11 M12 M13 M14 M15 M16 M17 M18 M19 M20 M21 M22 M23 M24
L1 L2 L3 L5 L6 WM
Regional Module Score
Age at Death (Years)
Control
Bulbar
Limb
pval < 0.05
B
Donor-level Correlation of Post-mortem Interval and MTC Spatial Module Score
Regional Module Score
Post-mortem Interval (Hours)
C
Donor-level Correlation of Disease Duration and MTC Spatial Module Score
Regional Module Score
Disease Duration (Months)

**Supplemental Figure 21: MTC Spatial gene module 17 is affected by post-mortem interval.**

The averaged spatial modules scores per AAR for each donor sample in the MTC ST data, plotted on the y-axis, compared to **A)** donor age at death, **B)** post-mortem interval (PMI), and **C)** ALS disease duration plotted on the x-axis. Each dot represents an individual donor. Plots include a least squares regression line and Pearson correlation coefficient, and are colored violet when correlations are significantly different from 0 (Wald test, $p < 0.05$, FDR-BH).

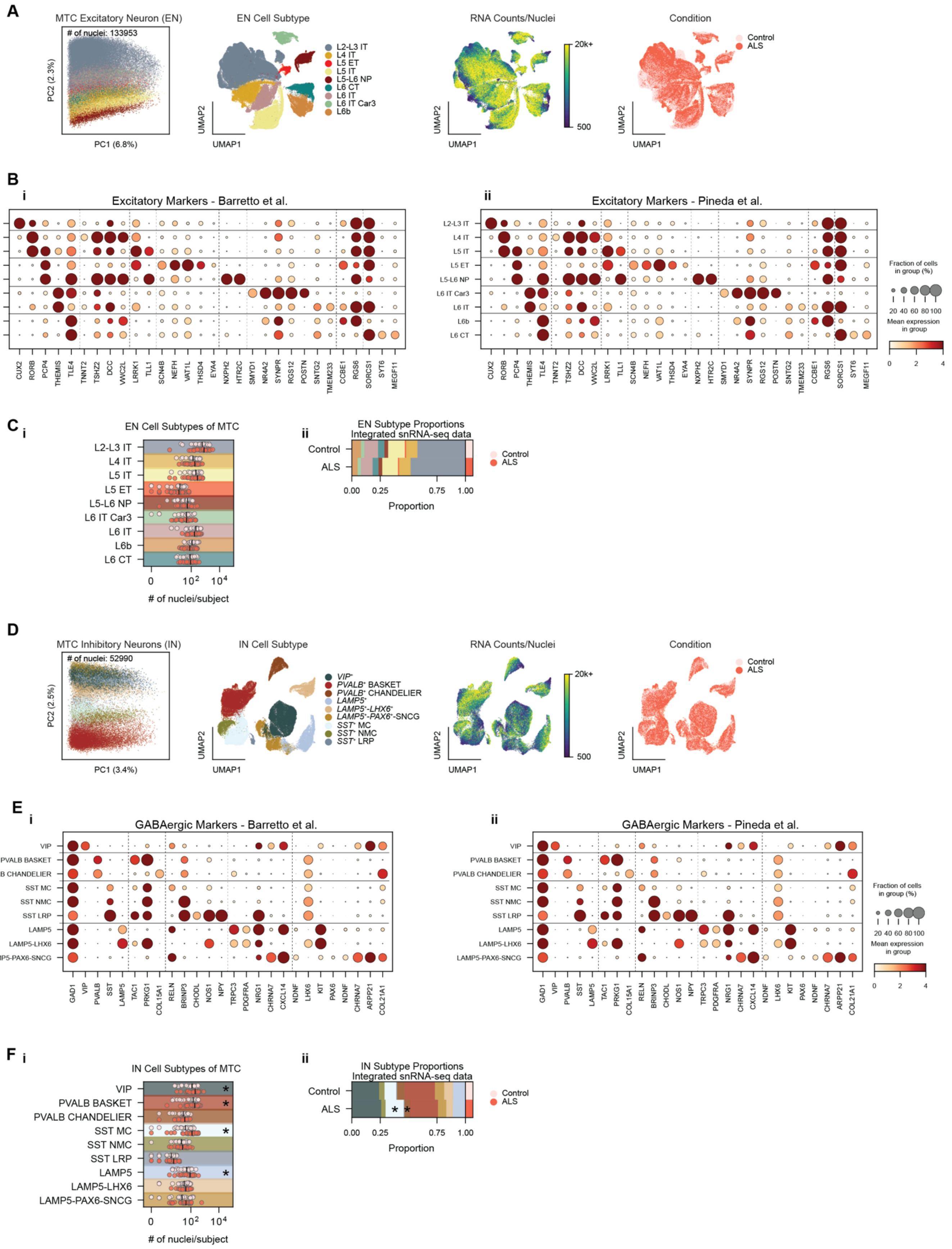
A
MTC Excitatory Neuron (EN)
### of nuclei: 133953
PC2 (2.3%)
PC1 (6.8%)
EN Cell Subtype
L2-L3 IT
L4 IT
L5 ET
L5 IT
L5-L6 NP
L6 CT
L6 IT
L6 IT Car3
L6b
UMAP1
UMAP2
RNA Counts/Nuclei
20k+
500
Condition
Control
ALS
B
i
Excitatory Markers - Barretto et al.
ii
Excitatory Markers - Pineda et al.
CUX2 RORB PCP4 THEMIS TLE4 TNNT2 TSHZ2 DCC VWC2L LRRK1 TLL1 SCN4B NEFH VAT1L THSD4 EYA4 NXPH2 HTR2C SMYD1 NR4A2 SYNPR RGS12 POSTN SNTG2 TMEM233 CCBE1 RGS6 SORCS1 SYT6 MEGF11
Fraction of cells in group (%)
20 40 60 80 100
Mean expression in group
0 2 4
C
i
EN Cell Subtypes of MTC
### of nuclei/subject
ii
EN Subtype Proportions
Integrated snRNA-seq data
Proportion
0.00 0.25 0.75 1.00
D
MTC Inhibitory Neurons (IN)
### of nuclei: 52990
PC2 (2.5%)
PC1 (3.4%)
IN Cell Subtype
VIP⁺
PVALB⁺ BASKET
PVALB⁺ CHANDELIER
LAMP5⁺
LAMP5⁺-LHX6⁺
LAMP5⁺-PAX6⁺-SNCG
SST⁺ MC
SST⁺ NMC
SST⁺ LRP
E
GABAergic Markers - Barretto et al.
GABAergic Markers - Pineda et al.
VIP
PVALB BASKET
PVALB CHANDELIER
SST MC
SST NMC
SST LRP
LAMP5
LAMP5-LHX6
LAMP5-PAX6-SNCG
GAD1 VIP PVALB SST LAMP5 TAC1 PRKG1 COL15A1 RELN BRINP3 CHODL NOS1 NPY TRPC3 PDGFRA NRG1 CHRNA7 CXCL14 NDNF LHX6 KIT PAX6 NDNF CHRNA7 ARPP21 COL21A1
F
IN Cell Subtypes of MTC
IN Subtype Proportions
*

**Supplemental Figure 22: Neuronal subtypes from integrated MTC snRNA-seq data.**

**A)** PCA and UMAP plot of excitatory neurons. Each data point represents one nucleus, color-coded by assigned cell subtype, RNA counts per nucleus, or condition.

**B)** Dot plot showing expression of key marker genes for each EN subtype in i) Barretto et al., and ii) Pineda et al. Color indicates mean gene expression, and dot size denotes the proportion of cells with non-zero counts, from 0-100%.

**C)** i) Number of nuclei annotated per EN subtype across all MTC snRNA-seq libraries, color-coded by condition. Black bars mark median nuclei counts; asterisks denote significant differences between ALS and control donors (Welch's t-test, $p < 0.05$, FDR-BH). ii) Averaged proportions of EN subtypes for controls and ALS donors. (Welch's t-test, $p < 0.05$, FDR-BH).

**D – F)** Same as B and C, shown for inhibitory neuron subtypes.

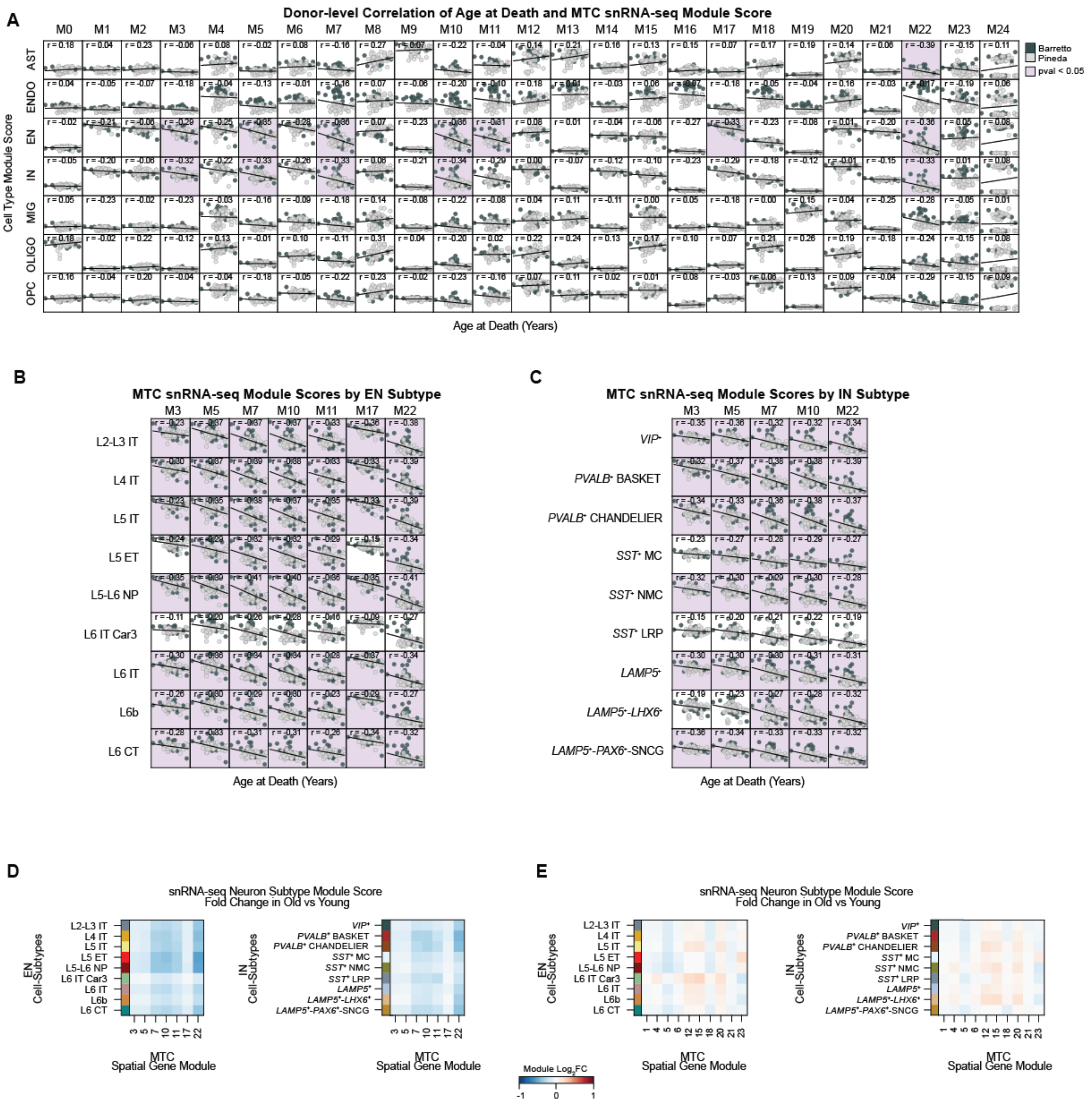
A
Donor-level Correlation of Age at Death and MTC snRNA-seq Module Score
M0 M1 M2 M3 M4 M5 M6 M7 M8 M9 M10 M11 M12 M13 M14 M15 M16 M17 M18 M19 M20 M21 M22 M23 M24
AST
ENDO
EN
IN
MIG
OLIGO
OPC
Cell Type Module Score
Barretto
Pineda
pval < 0.05
Age at Death (Years)
B
MTC snRNA-seq Module Scores by EN Subtype
M3 M5 M7 M10 M11 M17 M22
L2-L3 IT
L4 IT
L5 IT
L5 ET
L5-L6 NP
L6 IT Car3
L6 IT
L6b
L6 CT
Age at Death (Years)
C
MTC snRNA-seq Module Scores by IN Subtype
M3 M5 M7 M10 M22
VIP+
PVALB+ BASKET
PVALB+ CHANDELIER
SST+ MC
SST+ NMC
SST+ LRP
LAMP5+
LAMP5+-LHX6+
LAMP5+-PAX6+-SNCG
Age at Death (Years)
D
snRNA-seq Neuron Subtype Module Score
Fold Change in Old vs Young
EN Cell-Subtypes
IN Cell-Subtypes
MTC Spatial Gene Module
Module Log2FC
-1 0 1
E
snRNA-seq Neuron Subtype Module Score
Fold Change in Old vs Young
EN Cell-Subtypes
IN Cell-Subtypes
MTC Spatial Gene Module

**Supplemental Figure 23: MTC spatial gene module 5 alterations are driven by ALS, not age.**

**A)** Age at death plotted against snRNA-seq module scores per MTC cell type. Each dot represents an individual donor. Plots include a least squares regression line and Pearson correlation coefficient, and are colored violet when correlations are significantly different from 0 (Wald test, $p < 0.05$, FDR-BH).

**B)** Age at death plotted against snRNA-seq module scores per EN and **C)** IN subtype. The seven significant EN modules and four IN modules highlighted in panel A are shown. Each dot represents an individual donor. Plots include a least squares regression line and Pearson correlation coefficient, and are colored violet when correlations are significantly different from 0 (Wald test, $p < 0.05$, FDR-BH).

**D)** $Log_2$ fold change of spatial module scores correlated with age, shown for each snRNA-seq neuronal subtype. Donors were stratified into two age groups: young (34-67) and old (67+). Fold changes were calculated for old groups relative to young. Asterisks indicate significant differences from the young age group (Welch's t-test, $p < 0.05$, FDR-BH).

**E)** $Log_2$ fold change of spatial module scores altered in ALS, shown for each snRNA-seq neuronal subtype. Donors were stratified into two age groups: young (34-67) and old (67+). Fold changes were calculated for old groups relative to young. Asterisks indicate significant differences from the young age group (Welch's t-test, $p < 0.05$, FDR-BH).
